# Deep MALDI-MS Spatial ‘Omics guided by Quantum Cascade Laser Mid-infrared Imaging Microscopy

**DOI:** 10.1101/2023.12.14.571637

**Authors:** Lars Gruber, Stefan Schmidt, Thomas Enzlein, Huong Giang Vo, James Lucas Cairns, Yasemin Ucal, Florian Keller, Denis Abu Sammour, Rüdiger Rudolf, Matthias Eckhardt, Stefania Alexandra Iakab, Laura Bindila, Carsten Hopf

**Author notes:** these authors contributed equally (Lars Gruber, Stefan Schmidt). Corresponding author: Carsten Hopf, PhD.

## Abstract

In spatial ‘omics, highly confident molecular identifications are indispensable for the investigation of complex biology and for spatial biomarker discovery. However, current mass spectrometry (MS)-based spatial ‘omics must compromise between data acquisition speed and biochemical profiling depth, thus often leading to only “putative” molecular identifications. Here, we introduce fast quantum cascade laser mid-infrared imaging microscopy to guide MS imaging to confined tissue areas of high interest, *e.g.,* multicellular spheroid cores or kidney glomeruli, for spatial lipidomics profiling at maximized analytical depth utilizing magnetic resonance-MS imaging at >10^6^ resolution or prm-PASEF-MS^2^ fragmentation imaging. Instigating selective sulfatide accumulation in arylsulfatase A-deficient mice as ground truth concept, we demonstrate that deep QCL-infrared-guided *on-tissue* spatial ‘omics unequivocally identifies 120 sulfatides. This approach enables identifications of odd-chain sulfatides and studies of structure-ion mobility-relationships that provide chemical rationales for improvements to current ion mobility prediction algorithms. Workflows and data processing tools are provided as community resources.

## Introduction

Mass spectrometry imaging (MSI) is a fundamental label-free technology in spatial biology. It enables spatially resolved visualization, investigation and probabilistic mapping of lipids, metabolites, peptides, drugs or N-glycans in tissue sections in biomedical science as well as in clinical and industrial research^1–4^. Integration of whole tissue data sets sequentially acquired from the same or adjacent tissue sections by MSI and orthogonal technologies such as (mid-)infrared imaging (IRI), Raman imaging or spatial transcriptomics, often referred to as correlative spatial multi-omics^5–7^, have opened up new avenues for scientific inquiry. Recent high-end MSI platforms, including magnetic resonance (MR) - and trapped ion mobility spectrometry (TIMS) MSI, offer superior **<u>s</u>**peed, **<u>s</u>**ensitivity, **<u>s</u>**patial resolution, or molecular **<u>s</u>**pecificity^8, 9^. However, the four criteria in this 4S-paradigm of MSI performance are currently mutually exclusive, and very high molecular specificity and sensitivity can only be obtained by in-depth spatial chemical analysis at lower speed and at reduced image resolution, *i.e.* large pixel sizes^10^. Therefore, in practice, most MALDI-MSI studies with high-performance instruments today appear to use MR-MSI with mass resolving power below 100,000 or timsTOF-MSI in TIMS-off/qTOF mode without using collisional cross section (CCS) information. Because of time constraints and despite the lack of HPLC separation in MSI, analytical capabilities for spatially-resolved accurate mass determination, ultra-high resolution analysis of isotope fine structures (IFS; both in MR-MSI), on-tissue fragmentation analysis, and ion mobility separation of isobaric compounds (both in TIMS-MSI) are often not used. Without this information, lipids/metabolites cannot be considered confidently identified.

To overcome instrument limitations, “smart” data processing and “smart” sampling methods (reviewed in^11^) such as histology- or IRI-guided MSI have been suggested^12, 13^. In the latter case, a non-destructive, label-free and fast “guiding” imaging modality determines the composition of tissue specimens by mid-infrared vibrational spectroscopy. It captures collective molecular information as a rough molecular sketch, *i.e.,* relative lipid-, nucleic acid-, carbohydrate- and protein content and allows for computational segmentation of the imaging data. This enables effective IRI-based definition of regions of interest (ROI), *e.g.,* the cerebellar granular layer in mouse brain^12^. ROI information is then transferred to the slower imaging mass spectrometer for focused MSI analysis of well-defined, often small tissue areas. This saves time and data volume, which could, in principle, be spent for advanced chemical bioanalysis with LC-MS-like analytical depth, *i.e.* with ultra-high resolving power provided by MR-MS and/or by default use of TIMS in MSI studies.

However, IRI-guided MSI has remained a mere concept so far, as available Fourier transform (FT-IR) imaging instruments have not been fast enough for spatially focused tissue analysis at cellular resolution. In contrast, Quantum Cascade Laser (QCL)-based mid-IR imaging microscopes feature a tunable coherent IR source with high power-density for high sensitivity and higher sample throughput in biological systems^14–18^. This permits the selective acquisition of IRI data for user-defined single wavenumbers or full hyperspectral data. However, validated methods and dedicated computational tools for information-rich and high-throughput IRI-guided MSI are lacking^7, 15, 19–21^.

In addition to these challenges, method development and validation in MSI have generally been hampered by the lack of reliable analytical ground truths for segmentation and molecular identities^22, 23^, as the spatial and molecular composition of investigated tissues is typically unknown. To this end, synthetic datasets^2^, expert crowdsourcing^22^ single-cell fluorescence^24^, or histopathology annotations^25^ have been proposed as ground truths. To address this key challenge in MSI method development and validation, we propose that genetic mouse models with defined alterations in metabolism be used as qualitative ground truth: In case of QCL-IRI-guided MSI workflows we employed arylsulfatase A-deficient (ARSA-/-) mice, a model of human metachromatic leukodystrophy (MLD). In these mice, sulfatides, a family of sulfated glycosphingolipids, selectively accumulate in kidneys and other organs^26–28^. Because of this well-understood biology of ARSA-/- mice, we could use kidney sulfatides, whose masses, chemical sum formulae and structures (but not their quantities) are known, as qualitative ground truth for statistical evaluation of IRI-guided MSI methods, for method validation and for benchmarking against 4D LC-TIMS-MS sulfatide lipidomics, against previous sulfatide MSI studies and against CCS value prediction models^26, 28–41^.

Using a hybrid mid-IRI instrument for point-wise FT-IR spectroscopy and rapid QCL-IRI microscopy, we developed QCL-IRI-guided MSI workflows and computational tools for spatially resolved deep lipidomics profiling and make them and extensive spatial lipidomics data available as a community resource. On the one hand, this concept enables IRI-guided in-depth spatial chemical analysis of tissue ROIs using highly accurate ultra-high resolution MRMS imaging (R>10^6^ at *m/z* 800; mass accuracy <0.2 ppm) for improved use of IFS in lipid identification. These workflows also allow for IRI-guided detailed spatially resolved *on-tissue* MS^2^ of lipids in QCL-IRI-defined ROIs by means of parallel reaction monitoring with parallel accumulation and serial fragmentation (prm-PASEF) using ion mobilograms with optimized resolution of CCS values for precursor ion selection.

## Results

### Quantum cascade laser mid-infrared microscopy imaging to guide spatially focused data acquisition in MALDI imaging

Disentangling the molecular complexity of biological specimens by HPLC separation and thorough structural elucidation by MS^2^ fragmentation analysis have been the hallmarks of LC-MS-based lipid/metabolite analysis for decades. However, LC-MS-like analytical depth, sensitivity and confidence of identification is currently lacking in spatial MSI lipidomics.

As a solution to this conundrum, we proposed to spend more MSI analysis time and depth on user-defined morphological structures of interest, *i.e.,* on fewer pixels than those of entire tissue sections. Previously, we suggested mid-IR vibrational tissue imaging (FT-IRI) for ROI definition, followed by MSI restricted to these ROIs^12^. However, in the past, such an IRI-guided MSI workflow lacked IRI acquisition speed, provided insufficient molecular specificity or assessed few vibrational bands. It was therefore not feasible^42^. Meanwhile, QCL-IRI instruments with a focal plane array detector offer scanning capabilities with microscopy quality and improved sensitivity due to high power density and much higher speed for acquisition of full spectra in the “fingerprint” region (950–1800 cm^-1^; **Fig. 1a(i)**). The QCL-IRI microscope used within this study can analyze 5 million 5×5 µm^2^-sized pixels in 10 min (∼8750 pixels per sec versus 50 pixels per sec with FTIR imaging^12^) compared to 7h for 175,000 10×10 µm^2^- sized pixels in TIMS-MSI. Available coherence reduction ensures effective suppression of sample-dependent phase shifts for structured tissues where the objects of interest, *e.g.,* cells, have the same dimension as the wavelength of the light source (∼5 µm)^43^.

**Fig. 1.**
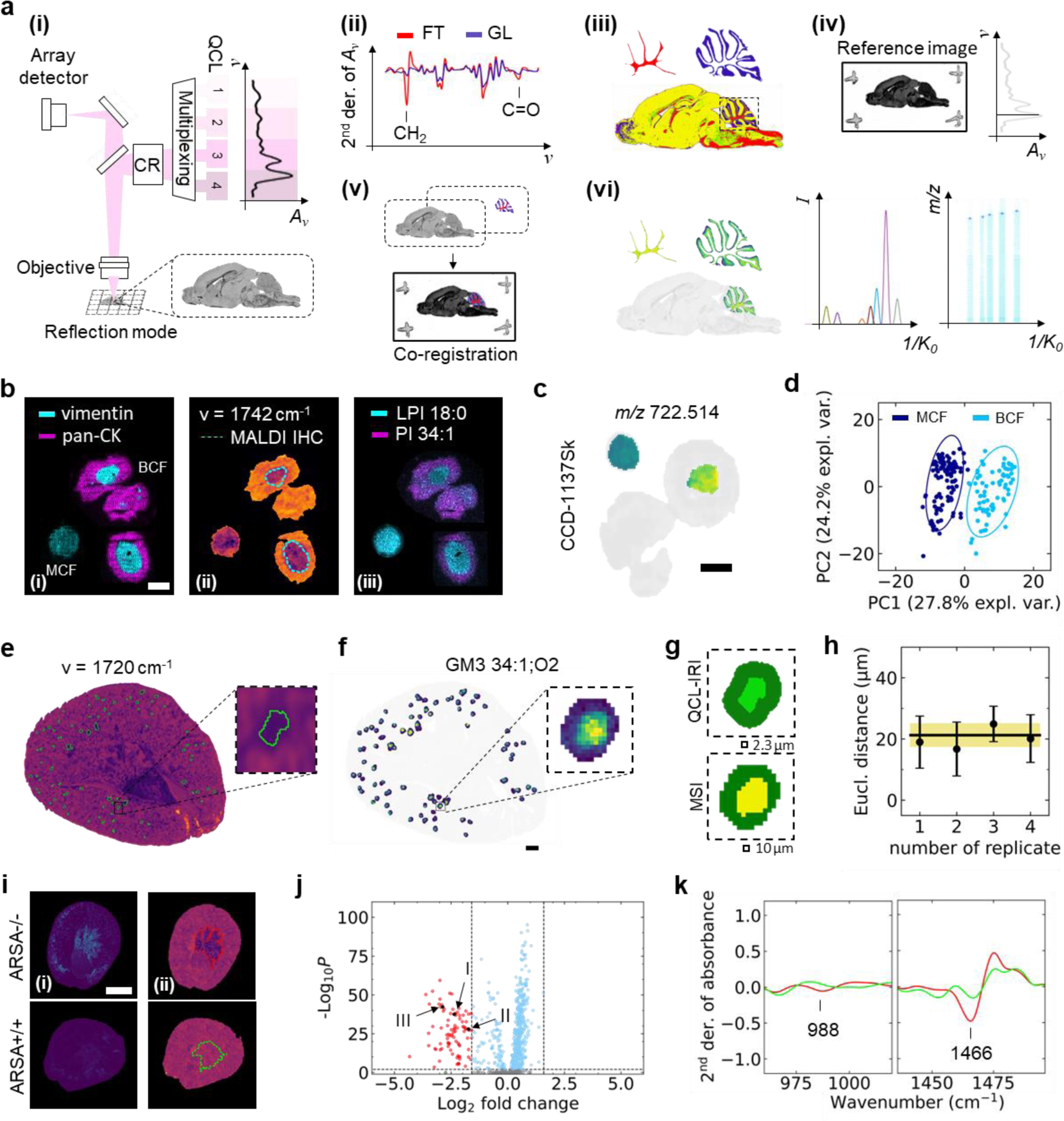
QCL-IRI microscopy to guide spatially focused data acquisition in MALDI MSI. **a,** Schematic overview of QCL microscope **(i)**, and QCL-IRI-guided MSI workflow on fresh-frozen biological specimen in reflection mode. A_ν_: absorbance, CR: coherence reduction. **(ii)** Spectral features (2^nd^ derivative of absorbance (2^nd^)) for cerebellar fiber tracts (FT) and granular layer (GL) are selected: 1466 cm^-1^ (CH_2_ bending vibration) and 1742 cm^-1^ (C=O vibration). **(iii)** Feature-selective image segmentation and region-of-interest (ROI) definition. **(iv)** *Single wavenumber* (1656 cm^-1^) data acquired as reference image and **(v)** co-registered with QCL-IRI dataset (i). **(vi)** Cerebellar ROI-focused MSI incl. mobility-based MS^2^ using prm-PASEF. **b,** Comparison of CCD-1137Sk fibroblast/HT-29 cancer biculture (BCF) versus monoculture fibroblast (MCF) spheroids. Scale bar, 200 µm. **(i)** Ion images of multiplex-MALDI-IHC^45^ using anti-vimentin (*m/z* 1230.84; fibroblast) and anti-pan-CK (*m/z* 1288.71; cancer) antibodies. Mass window ±10 ppm. **(ii)** QCL-IR data (1742 cm^-1^; 2^nd^) of same spheroid section, fibroblast core outline derived by clustering (k=2) of MALDI-IHC data. **(iii)** Ion images for *m/z* 835.54 (PI 34:1[M-H]^-^; fibroblasts) and *m/z* 599.32 (lyso-PI 18:0[M-H]^-^; cancer). **c**, QCL-IRI-guided ion image of *m/z* 722.51 (PE(P-36:4[M-H]^-^) discriminating MCF and BCF. **d**, Principal component analysis (PCA) of MSI data distinguishes MCF (dark blue) and BCF (cyan). **e,** QCL-IRI of ARSA-/- mouse kidney (1720 cm^-1^; 1^st^) with putative glomerular ROI (green outline). **f**, QCL-IRI-guided ion image of *m/z* 1151.71 (GM3 34:1;O2[M-H]^-^; mass window ±10 ppm) with highlighted glomerular ROI. Scale bar, 300 µm. **g**, Comparison of dilated QCL-IRI-ROI (top) and MSI-feature-based clustering (k=2; bottom). **h**, Mean Euclidean distance (21±4 µm) between centers-of-gravity of QCL-IRI-ROI and glomeruli cluster (MSI; yellow; *n*=4). **i**, Sulfatide accumulation in ARSA-/- mice by MSI and QCL-IRI. **(i)** Sum intensity distribution (87 sulfatides^28^). **(ii)** QCL-IRI at 988 cm^-1^ (2^nd^, C_β_-O vibration). Superimposed kidney inner segment of outer medulla (ISOM) ROI determined by clustering of MSI data (red and green dashed lines). Scale bar, 2 mm. **j**, Volcano scatter plots of qTOF-MSI data for ARSA-/- vs ARSA+/+ reveals accumulation of sulfatides^28^ for ARSA-/- (red dots), e.g., **I)** SM4 34:1;O2[M-H]^-^, **II)** SM4 38:1;O3[M-H]^-^, and **III)** SM3 42:1;O2[M-H]^-^. **k**, QCL-IRI spectra, highlighting features at 988 cm^-1^ and 1466 cm^-1^ (2^nd^).

To maximize analytical depth per pixel, we set out to develop experimental workflows and IT tools for spatially focused MSI data acquisition on fewer pixels preselected by QCL-IRI (**Fig. 1a; Supplementary Fig. 1**): As demonstrated for murine brain tissue sections, mid-IR “fingerprint” absorbance spectra are first recorded in reflection mode using indium tin oxide (ITO) glass slides (**Supplementary Fig. 2**). QCL-IRI spectral features discriminating between tissues, tissue morphologies, or cell types are then extracted from the IR “fingerprint” spectra^21^ and used to segment QCL-IRI data, *e.g.* by performing 2^nd^ derivative spectroscopy^21^ (**Fig. 1a(ii)**), and to define ROIs (**Fig. 1a(iii)**). A separate single wavenumber (1656 cm^-1^) whole-slide QCL-IR reference image is co-registered with the “fingerprint” data set to generate the MSI data acquisition file, thus effectively transferring QCL-IRI-defined ROIs to the TIMS- or MR-mass spectrometer for ROI-targeted MSI with maximized analytical depth (**Fig. 1a(iv) to Fig. 1a(vi)**).

To validate this workflow, we first examined whether photonic interaction of the coherent QCL light with the tissue’s molecules may cause lipid alterations^7^. However, no marked lipid changes (*m/z* 600–1700) were observed in murine brain tissue sections irradiated with the high-power laser for 15 min in either *single wavenumber* or *sweep scan* mode (**Supplementary Fig. 3 and 4**).

To ensure wide applicability of the QCL-IRI-guided MSI approach, we investigated three different biomedical examples. First, we examined a two-cell types-two ROIs *in-vitro* 3D-cell culture model of cancer-like aerobic glycolysis and reverse Warburg effect^44^. This model features biculture spheroids (300-500 µm diameter) consisting of two human cell lines, vimentin-positive CCD-1137Sk fibroblasts forming the core and pan-cytokeratin (pan-CK)-positive HT-29 colon cancer cells that engulf them, as confirmed by multiplex-MALDI-immunohistochemistry (IHC)^45^ (**Fig. 1b(i)**). Analysis of hyperspectral QCL-IRI “fingerprint” data revealed the lipid-associated bands at wave-numbers 1466 cm^-1^ (CH_2_ bending vibration) and 1742 cm^-1^ (C=O vibrations) as molecular features capable of distinguishing between the two cell types, as indicated by the precise match of the fibroblast core outlines determined by MALDI-IHC and QCL-IRI (**Supplementary Fig. 5a; Fig. 1b(ii)**). Consistent with recent observations in (brain) tumor patient samples where transcripts of glycerophospholipid (GPL) remodeling enzymes were overexpressed compared to surrounding non-tumor tissue^2^, GPLs such as phosphatidylinositol PI 34:1 were more prominent in cancer cells, whereas lyso-GPLs, *e.g.* LPI 18:0, were more abundant in fibroblasts (**Fig. 1b(iii)**).

As a feasibility study, we compared monoculture fibroblast (MCF) spheroids with the QCL-IRI-guided fibroblast core of bicultures (BCF), to investigate lipidomic reprogramming of BCF (compared to MCF) induced in the latter by being surrounded and separated from culture medium by cancer cells^46^.

Principal component analysis on 105 MCF- and 72 BCF spheroids (mean MSI profiles, technical replicates) of the MSI data indicated unique lipidomic profiles for both 3D-cellular systems (**Fig. 1d)**. Chemometrics- and machine learning-based feature extraction (LASSO regression) revealed lipid candidates, *e.g.,* the ether phosphatidyl-ethanolamine PE(P-36:4), with different abundance in BCF than MCF (**Fig. 1cd; Supplementary Fig. 6 and 7; Supplementary Tables 1–9**).

Next, we explored small tissue morphologies like glomeruli in murine kidney to further validate QCL-IRI-based hyperspectral tissue segmentation. Gangliosides as markers of these functional filtration units are well-characterized, and autofluorescence-directed MSI has recently been used for their detailed molecular analysis^38^. Using ganglioside GM3 34:1;O2 (both negative mode ion images and molecular probabilistic maps (MPMs)^2^) as a marker, we compared MALDI-qTOF-MSI and QCL-IRI for entire dried kidney cryosections by correlative MSI-IRI imaging (**Supplementary Fig. 8a and 8b**). Utilizing the spectral region at around 1720 cm^-1^ (1^st^ derivative of transmittance) between the amide I and C=O vibrations discriminated between glomerular structures and surrounding kidney cortex. Additionally, ROIs were chosen such that the number of objects identified by both QCL-IRI and MSI compared to the latter alone was maximized, thus rather accepting false-negatives, but avoiding false-positive detections in QCL-IRI (**Fig. 1e; Supplementary Fig. 8c**). On average, more than 85% of QCL-IRI-defined ROI contained glomeruli, as defined by GM3 34:1;O2 presence (**Supplementary Fig. 9**). To compensate for possible errors in image co-registration, IRI-defined ROIs were computationally dilated. To evaluate the magnitude of these effects, the Euclidean distance between the centers of the dilated QCL-IRI-defined ROI and the GM3 34:1;O2-defined MSI area determined the off-set (**Fig. 1fg; Supplementary Fig. 9a-d**). Averaged across all glomeruli in four independent data sets, the spatial shifts were 21±4 µm compared to QCL-IRI- (*i.e.,* the taught reference image) and MSI pixel sizes of 2.3 µm and 10 µm, respectively (**Fig. 1g and 1h; Supplementary Fig. 9a-d).** Focusing MSI data acquisition on these glomeruli-containing ROIs instead of full tissue MSI reduced data acquisition time by >95% (8483 pixels instead of 216,411). This allowed for subsequent redirection of time and effort into unequivocal identification of ten gangliosides in these ROIs by MRMS accurate mass analysis and ion mobilogram-based on-tissue fragmentation using prm-PASEF in TIMS-MSI (**Supplementary Fig. 9e-i; Supplementary Table 10**).

### Genetically engineered mice with defined metabolic alterations as a qualitative ground truth in MSI method development and validation

Using our QCL-IRI-guided MSI toolbox, we endeavored to chemically characterize an entire lipid class, sulfatides, as comprehensively and completely as possible, since it is neither sufficiently covered in public databases (SwissLipids knowledgebase, https://swisslipids.org/; LIPID MAPS Structure Database, https://www.lipidmaps.org/) nor in instrument vendor software. Furthermore, we aimed to provide methods for ultra-high resolution (R >10^6^) MR-MSI and prm-PASEF MS^2^ with tailored ion mobility separation, *i.e.* using long ramp times, to analyze QCL-IRI-defined ROIs and to generate an extensive comparative MSI- and LC-TIMS-MS data resource for the MSI and lipidomics communities.

To this end, we introduce the concept of using knock-out mice to approach analytical ground truths in complex tissue analytics. To this end, we used kidneys of arylsulfatase A-deficient mice (ARSA-/-) that we first analyzed in 2011 using low-resolution MALDI-TOF MSI incapable of accurately identifying lipids.^28^ (**Supplementary Table 11**). In kidneys of ARSA-/- mice, sulfatides of the SM4, SM3, SM2a and SB1a subclasses accumulate primarily in medulla and papilla^26–28, 47^ (**Fig. 1i and j**; **Supplementary Fig. 10** for sulfatide metabolism), where they are known to be critical for urinary pH and ammonium excretion and have been characterized at single intercalated cell level^35, 39^.

In brain, they promote neurodegeneration in MLD. Sulfatide accumulation was observed consistently across MSI and QCL-IRI (**Supplementary Fig. 11**), and the inner segment of outer medulla (ISOM) and inner medulla/papilla (IMP) ROIs of kidney could readily be segmented using a well-defined subset of spectral features in the “fingerprint” region, *i.e.,* the lipid-associated CH_2_ bending vibration at 1466 cm^-1^ and the C_β_-O vibration of the 3-sulfogalactosyl head group at 988 cm^-1^ (**Fig. 1k; Supplementary Fig. 5b**). Comparison of segmentations based on either imaging modality and with reference histology suggested that QCL-IRI did not alter coverage of the ISOM and IMP ROIs (**Supplementary Fig. 12**).

### QCL-IRI-guided ultra-high resolution magnetic resonance-MSI improves accuracy, utilization of isotope fine structures and FDR-controlled sulfatide identification

To investigate the analytical benefits of reinvesting data acquisition time saved by the QCL-IRI-guided MSI approach in MR-MSI at very mass resolution, we established a ground truth for sulfatide identification in ARSA-/- mice. First, we considered 156 structural configurations for each of the sulfatide subclasses SM4, SM3, SM2a, SM1a and SB1a, *i.e.* all sphingoid base-FA compositions from 32:x to 46:x with x = 0,1,2,3 for O2,O3 and x = 0,1 for O4 (in total 150) plus six sphingosine-compositions (**Fig. 2a**). As these were within the mass range *m/z* 540-1510, we considered them potentially detectable by MSI. SB1a is detected as SM1a in MALDI-MSI, due to the in-source loss of a sulfate^28^. Then we studied kidneys of two 12-week-old and two 60-week-old ARSA-/- mice *in-depth* by LC-based 4D-TIMS-PASEF (**Supplementary Table 12**) and identified 90-92 sulfatides (with structural confirmation by PASEF; **Table 1**)^48^. Finally, we compared conventional MR-MSI of whole kidney slices at mass resolution of R_1_∼77,000 at *m/z* 800 (1s free induction decay (FID) time; 14,331 40 µm pixels; 5h) in the FT-ICR with QCL-IRI-guided analysis focused to the ISOM and IMP ROIs but using R_2_∼1,230,000 at *m/z* 800 (16s FID time; 2672 40 µm pixels; 11.6 h) and benchmarked it against the ground truth (**Fig. 1jk; Fig. 2**; **Table 1; Supplementary Fig. 13; Supplementary Table 13**).

**Fig. 2.**
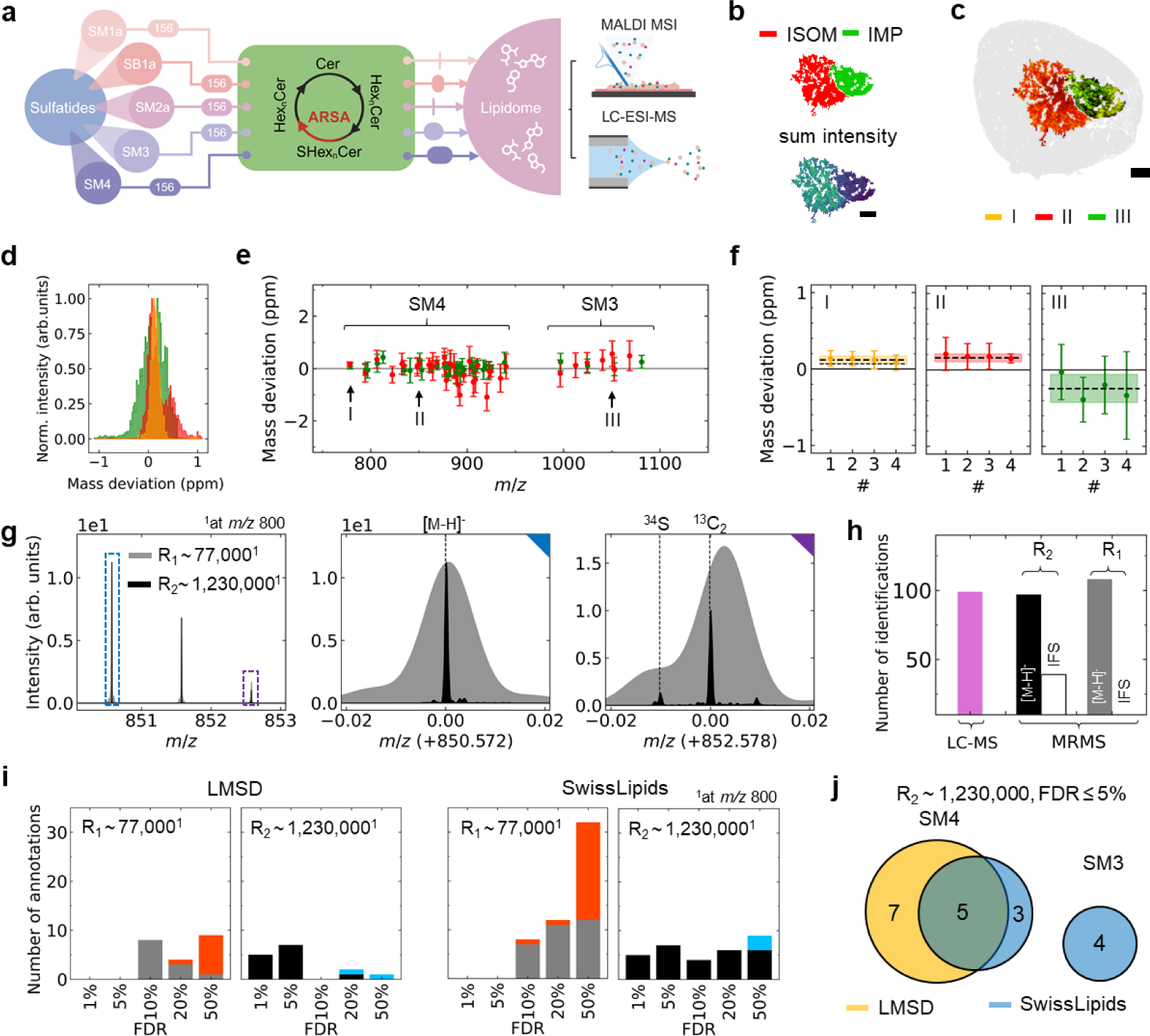
Deep sulfatide profiling via QCL-IRI-guided ultra-high resolution MR-MSI in ARSA-/- mouse kidney. **a,** Introduction of ground truth (GT) concept. For each sulfatide subclass, 156 theoretical configurations were hypothesized and evaluated for their real detectability in the lipidome via deep LC-ESI-MS and MALDI-MSI profiling. **b,** Inner medulla/papilla (IMP; green) and ISOM (red) identified as ROIs via QCL-IRI (top) and sulfatide distribution in MSI (sum intensities; bottom). Scale bar, 500 µm. **c**, Overlaid QCL-IRI ion image of **I)** *m/z* 778.5146 (SM4 34:1;O2[M-H]^-^; orange), **II)** *m/z* 850.5721 (SM4 38:1;O3[M-H] ^-^; red), and **III)** *m/z* 1052.6923 (SM3 42:1;O2[M-H] ^-^; green) within kidney (grey). Mass window ±3 ppm. **d,** Histogram of sum intensities in ISOM and IMP for I), II), and III) measured with a mass resolution of ∼1,230,000 (1230k) at *m/z* 800. **e,** Mean mass deviation (±σ of intensity distribution in d) of 47 sulfatides (*m/z* present in ≥50 pixels) at R_2_∼1230k with higher ion intensity in IMP (green) or ISOM (red). *m/z* values are shifted by +0.2 (IMP) or −0.2 (ISOM) for visualization. **f,** Weighted mean (dotted line) and uncertainty (*n*=4) presented as internal error^49^ (filled area) for mass deviation for I), II), and III). **g,** Ultra-high resolution MRMS data acquired with a mass resolution of R_1_∼77k (gray) and R_2_∼1230k (black). Isotopic fine structure (IFS) of SM4 38:1;O3[M-H]^-^ incl. ^13^C_2_ (M+2) and ^34^S isotopic peaks, normalized to the monoisotopic peak. **h,** Putative identifications of sulfatides by LC-TIMS-MS and by MALDI MR-MSI for R_1_∼77k (grey; whole kidney MSI) versus R_2_∼1230k (black; QCL-IRI-guided MSI) mass resolution. IFS is resolved for 33 sulfatides (R_2_∼1230k). **i,** Number of annotations in Metaspace (Swiss Lipids and Lipid Maps databases) of molecules containing a sulfate group (cyan and red bars) and of manually curated sulfatides (gray and black bars) based on MALDI MSI data. **j,** Venn diagram of SM3/SM4 isoforms identified by ultra-high resolution MR-MSI in the two databases that very incompletely cover sulfatides.

**Table 1.**
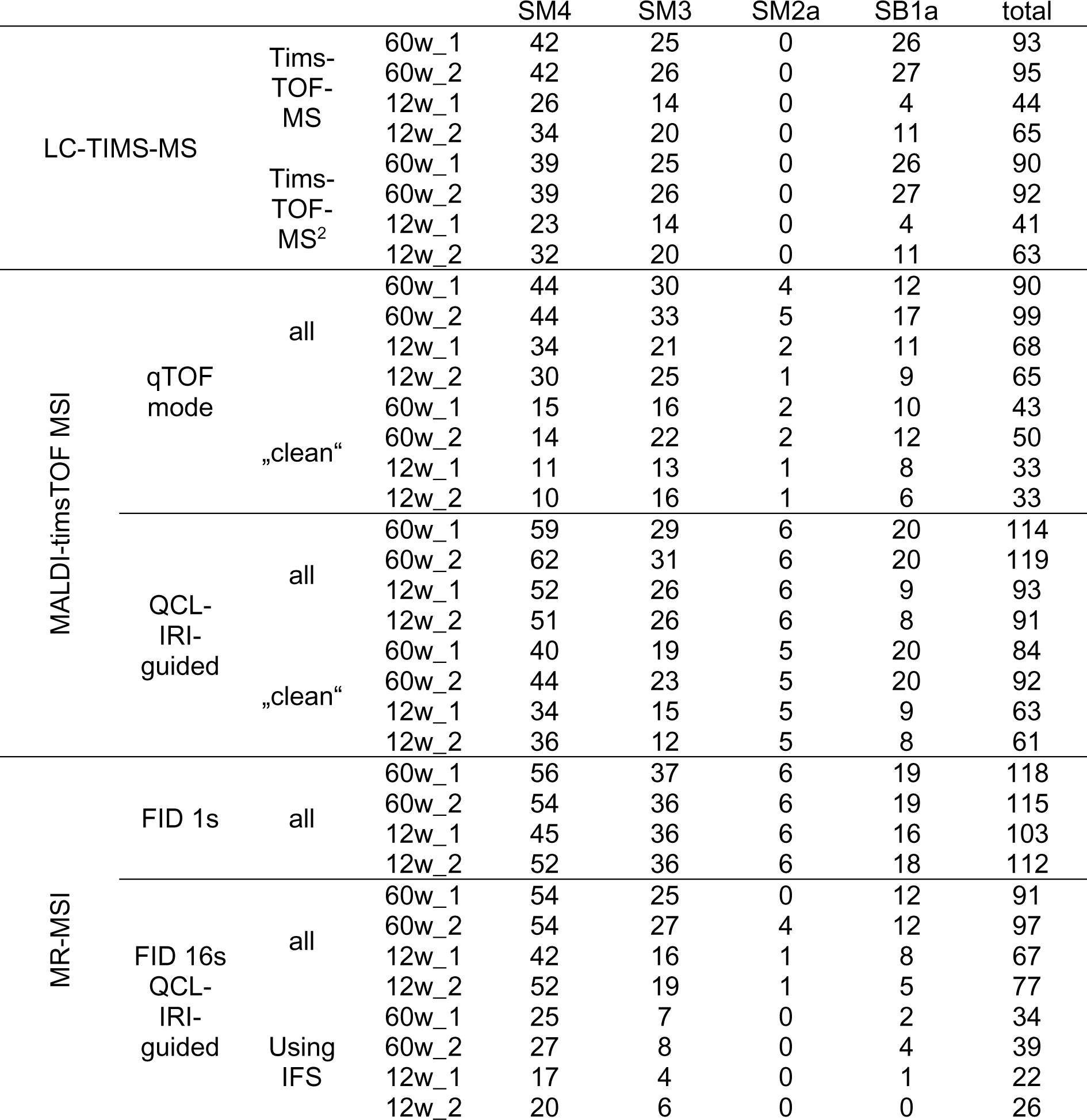
Cumulative numbers of sulfatide subclass isoforms identified in ARSA -/- mouse kidney by LC-MS or MALDI-MSI using various analytical workflows. Whole kidney sections of 12- or 60-week-old ARSA-/- mice (*n*=2 each) were analyzed by LC-tims-TOF-MS or –MS^2^, by MALDI-qTOF-MSI or by MALDI-MR-MSI with a mass resolution R_1_∼77k. “Clean” peaks fulfilled higher quality standards, defined as peaks that do not feature at least one other peak within a mass window of ±1.1 Da (qTOF mode MSI) and a mobility window of ±0.005Vs/cm^2^ (TIMS-MSI), and were manually curated. QCL-IRI-guided MSI focused on the kidney’s ISOM and IMP ROIs only. In many cases of MR-MSI with mass resolution of R_2_∼1230k, isotope fine structures (IFS) could be used for added confidence.

In this example, measurement time was reduced 5.6-fold (11.6h instead of 65h) by measuring ISOM/IMP instead of the entire tissue section with ultra-high mass resolution. Sulfatide sum intensities were about 2-3-fold higher in ISOM than in IMP (**Fig. 2b**). As ion suppression is likely different in these two areas of salt-rich kidney, ion intensities do not directly translate to quantities^28, 38, 47^. Nevertheless, cross-modality (semi-)quantitative analysis (**Supplementary Fig. 11**) showed consistent relative accumulation of sulfatides across the different ARSA conditions. Three sulfatides serve as examples of compounds with similar intensities in both ROIs (**I,** *m/z* 778.5146 (SM4 34:1;O2[M-H]^-^)), more intense in ISOM (**II,** *m/z* 850.5721 (SM4 38:1;O3[M-H]^-^)), or more intense in IMP (**III,** *m/z* 1052.6923 [SM3 42:1;O2[M-H]^-^); **Fig. 2c**). The maximum mass deviation was ∼1 ppm for the two less intense sulfatide ions **I** and **III** and <0.2 ppm for ion **II** with mass resolution R_1_∼77,000, but it was improved to <0.2 ppm for **I** and **II** (∼0.5 ppm for **III**) with a mass resolution of R_2_∼1,230,000.

Mass deviation was constant across the *m/z* range, reproducible in four mice, and the uncertainty of *m/z* values was reduced by a factor of about 3-6 for data acquired using ultra-high resolution FT-ICR. (**Fig. 2d to f; Supplementary Fig. 13a-d**). Furthermore, the IFS, particularly the peak attributed to ^34^S, was very well resolved by about 10-times FWHM (**Fig. 2g**). Due to predominant presence of some sulfatides in kidney cortex, the number of candidate sulfatide identifications with the ultra-high-mass resolution method (91 and 97 for two 60-week ARSA-/- mice) was lower than with the “conventional” whole-tissue method (118; 117). The identification confidence was, however, much improved, as 34 and 39 ultra-high-mass resolution spectra were supported by IFS information (**Fig. 2hi**; **Table 1**). Identification was performed manually, since IFS is currently not properly utilized in commercial sum formula IT tools that match experimental against theoretical isotope patterns^50^. In the sulfatide case, the ^34^S isotope peak was not automatically recognized as such, but S and N as constituents of the molecule-of-interest had to be predefined in the tool for successful searches. A case in point, this emphasizes the importance of the proposed ground truth model. For comparison, LC-TIMS-MS yielded 93 and 95 identifications (**Supplementary Table 12**).

More importantly, the number of decoy database-controlled sulfatide identifications with low false-discovery rate (FDR) in Metaspace (http://www.metaspace2020.eu) searches^51^ against the LipidMaps and SwissLipids databases was improved considerably, as none were identified within the 77k mass resolution data with FDR ≤5% (**Fig. 2i**). As these databases do not yet sufficiently cover sulfatides, only a small subset of about 15% (FDR ≤5%; gray and black bars in **Fig. 2i**) of all sulfatides identified from the ground truth was database-annotated. Combining the results from both databases, in total 19 sulfatides were assigned with FDR ≤5% (**Fig. 2j**). In addition, the number of improbable annotations with FDR ≥20% (red and blue part of the bars in **Fig. 2i**) was reduced using ultra-high mass resolution. FDR-based quality measures have recently been critically evaluated in the much more mature technology field of proteomics, as i) FDR determination is often a black box, and it is often unclear ii) at what level FDR is determined^52^. For MSI, the FDR-control process in Metaspace is rather transparent^51^. However, comparison of our experimental sulfatide data with the ground truth of conceivable or 4D-TIMS-PASEF verified sulfatides demonstrates that even ultra-high mass resolution MSI (MS^1^) does not guarantee unequivocal lipid identifications, since even mass determination with maximum accuracy is limited to the MS^1^ level.

### QCL-IRI-guided trapped ion mobility-MSI enables extensive fragmentation analysis on tissue and sulfatide identifications on par with LC-based 4D-TIMS-PASEF

TIMS-TOF-MSI, in principle, offers substantial capabilities for deep lipidomics profiling^48, 53^. However, as the use of TIMS separation slows down MSI data acquisition up to 10-fold, in practice, most imaging studies refrain from using the TIMS capabilities and are conducted in qTOF/TIMS-off mode instead. Hence, we investigated the gains in deep spatial sulfatide profiling that can be achieved by in-depth analysis of smaller ARSA-/- kidney ROIs (ISOM and IMP), as defined by QCL-IRI (compare **Fig. 2b**). The spatial distribution of the three example sulfatides **I** to **III** measured by TIMS-MSI resembled closely the ones obtained by MR-MSI (**Supplementary Fig. 14**; compare **Fig. 2c**). Then we comprehensively compared conventional qTOF-mode MSI of whole kidney slices (including on-tissue MS^2^ based on *m/z* values for precursor selection alone) with QCL-IRI-guided analysis focused to the ISOM and IMP ROIs but using TIMS-MSI with 2D-mobilogram-based prm-PASEF fragmentation, and benchmarked it against the ground truth (**Fig. 3**; **Table 1; Supplementary Table 12 and 13**).

**Fig. 3.**
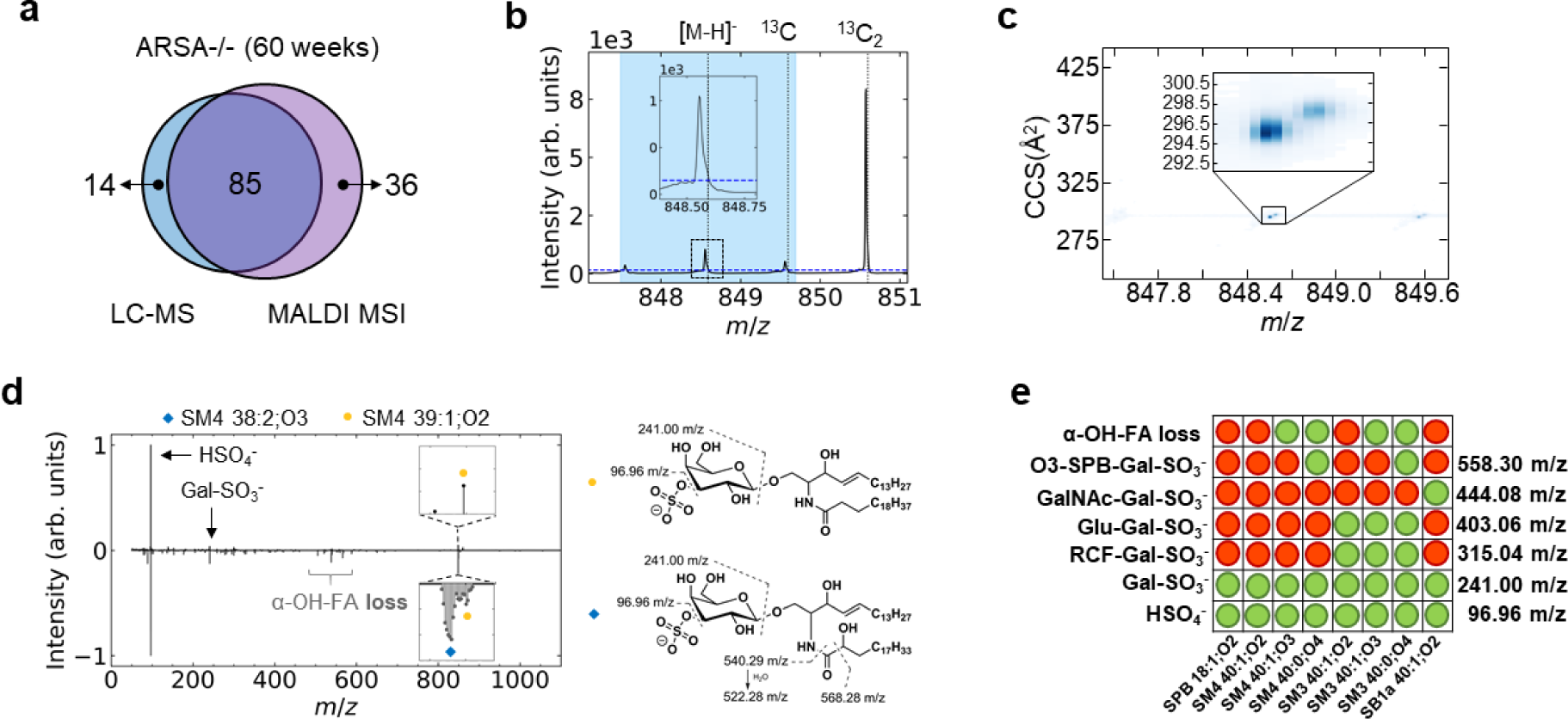
Ground truth evaluation and deep ion mobility profiling via QCL-IRI-guided tims-TOF MSI in ARSA-/- kidney. **a,** Sulfatides identifications by LC-TIMS-MS (96 of 99 confirmed by PASEF) and MALDI-TIMS-MSI (46 of 121 confirmed by prm-PASEF), both on timsTOF MS platforms across different laboratories. **b**, Example of a “non-clean” peak: *m/z* 848.557 (SM4 38:2;O3[M-H]^-^) displays an asymmetric peak shape because of non-resolved *m/z* 848.591 (SM4 39:1;O2[M-H]^-^). Precursor isolation window of ±1.1 Da for MS^2^ is depicted as light blue area. The blue horizontal dashed line (inset) refers to the intensity value at ^13^C of [M-H]^-^. **c,** Ion mobility heat map with inset highlighting two SM4 isoforms of *m/z* 848.557, 295.4 Å² (SM4 38:2;O3[M-H]^-^; left peak) and *m/z* 848.591, 297.6 Å² (SM4 39:1;O2[M-H]^-^; right peak). **d,** Butterfly plot of prm-PASEF-derived MS^2^ spectra (top) and conventional on-tissue MS^2^ without ion mobility separation (bottom) for *m/z* 848.591 (SM4 39:1;O2[M-H]^-^). The insets visualize the inability of conventional on-tissue MS^2^ to isolate SM4 39:1;O2 without SM4 38:2;O3 at *m/z* 848.557. Elucidated fragment patterns for SM4 38:2;O3[M-H]^-^ and SM4 39:1;O2[M-H]^-^. Fragments at *m/z* 568.28, *m/z* 540.29, and *m/z* 522.28 refer to the loss of the α-OH-FA. **e**, Fragments observed (green dots) in prm-PASEF MS^2^ analysis of selected sulfatides (columns).

For example, in non-guided whole kidney qTOF-mode MSI, 54,100 pixels were measured in 1:22h; in QCL-IRI-guided TIMS-MSI 9600 pixels were assessed in 1:19h (both 20 µm lateral step size). For prm-PASEF at 40 µm lateral step size 883 pixels were investigated in IMP (8 min) and 1554 pixels in ISOM (13 min). MALDI-MSI and 4D-LC-TIMS-MS lipidomics were on par in this comparison: In two 60-week-old mouse kidneys, 93 and 95 sulfatides were identified by 4D-TIMS-PASEF (90 and 92 with extra MS^2^ validation), 90 and 99 by TIMS-MSI in qTOF mode and 114 and 119 by QCL-IRI-guided TIMS-MS (**Fig. 3a**; **Table 1**). Four sulfatide subclasses were detected in method-specific manner (**Supplementary Fig. 15**). SM2a isoforms were only identified with the latter method, as they may have been too dilute in whole-kidney LC-TIMS-MS analysis, as in QCL-IRI-guided MSI (MR and TIMS) their signal-to-noise ratios were below 10. SB1a isoforms were only observed by LC-TIMS-MS, but due to sulfate loss in the MALDI ion source^28^ they were observed as SM1a in MSI instead. In total, we found about 120 sulfatides in the ARSA-/- mouse model out of 780 structural configurations considered as ground truth. Contrary to our expectation, we identified additional sulfatides in MSI that were not observed in LC-TIMS-MS. However, our findings validate the use of mutant mice with known patterns of metabolite accumulation in distinct tissue regions as ground truth in spatial lipidomics/metabolomics.

To aid statistical quality assessment, we defined “non-clean” sulfatide peaks for *m/z* spectra in qTOF-mode MSI as the ones that featured at least one other peak within a mass window of ±1.1 Da that exceeded the intensity of the first sulfatide isotope peak; the reason being that this interfering peak would prevent unequivocal precursor ion selection and on-tissue MS^2^ sulfatide identification in “conventional” qTOF mode. In TIMS-MSI operation, peaks were classified as “non-clean” if the spectra contained a peak within a mass window of ±1.1 Da and a mobility window of ±0.005 Vs/cm^2^. All peaks were manually classified as “clean” or “non-clean” (**Fig. 3bc**; **Table 1; Supplementary Fig. 16**). Of the 90 and 99 sulfatides seen in qTOF-mode MSI and the 114 and 119 detected in TIMS-mode MSI, about half (43 and 50) and about 75% (84 and 92) were considered “clean”, respectively (**Table 1**).

QCL-IRI-guided TIMS-MSI with prm-PASEF enabled new applications in deep spatial lipidomics, *e.g.,* the spatial profiling of odd-chain sulfatides (OCS), a lipid class that is still underexplored. Odd-chain fatty acids (FAs) can be formed by elongation of gut-/microbiome-derived propionyl-CoA but also via α-oxidation of 2-OH-FA by 2-hydroxy acyl-CoA lyase in mammalian cells^54, 55^. The example of *m/z* 848.557 (SM4 38:2;O3[M-H]^-^) and *m/z* 848.591 (SM4 39:1;O2[M-H]^-^) demonstrated that non-resolved (“non-clean”) odd-/ even-chain sulfatides could be separated by ion mobility spectrometry and subsequently identified by prm-PASEF fragmentation analysis (**Fig. 3b and 3c; Supplementary Fig. 19; Supplementary Table 14**). The ability to unequivocally identify odd-chain membrane lipids also applied to PI isomers like PI 33:1 in the spheroid example (**Supplementary Fig. 17**).

Altogether, QCL-IRI-guidance to ISOM and IMP ROIs allowed for extensive prm-PASEF fragmentation analysis and structure elucidation of odd- and even chain-sulfatides directly on-tissue (**Fig. 3d; Supplementary Fig. 18 and 19**). Surprisingly and in contrast to even-chain sulfatides like SM4 38:2;O3[M-H]^-^, ion intensities for OCS like SM4 39:1;O2[M-H]^-^ were unaltered in ARSA-/- kidneys, as indicated by ion images, volcano plots and example spectra (**Supplementary Fig. 18a**). As prm-PASEF analysis unequivocally identified OCS, this finding may suggest that odd-chain galactosylceramides and lactosylceramides can be sulfated by cerebroside sulfotransferases (**Supplementary Fig. 10 and 18b**), but that an arylsulfatase other than ARSA may catalyze their de-sulfation. This observation may be pursued in a separate study, as arylsulfatases constitute a growing enzyme family whose functions are not fully characterized yet^56^. In similar fashion large numbers of sulfatides could be structurally elucidated by TIMS-MSI with prm-PASEF (**Fig. 3e; Supplementary Fig. 19, Supplementary Table 14**).

### Structure-CCS-relationships based on experimental deep profiling spatial lipidomics data

This vast resource of spatial MALDI-MSI and corresponding LC-MS data, both acquired on TIMS-MS platforms, permitted deep inquiries into the (spatial) sulfoglycolipidome. First, experimental CCS values (LC-MS and MSI) differed profoundly from those predicted by recent models LipidCCS, DeepCCS and AllCCS2^30, 32, 34, 57^ (**Fig. 4a**). In contrast, experimental CCS values were very consistent and independent of ion source and instrument usage at different sites (Mannheim and Mainz; R^2^ = 0.9988; mean relative error 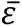 of 0.5%; **Fig. 4b; Supplementary Table 12 and 13**). LipidCCS was accurate (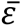 of 1%) for SM4, but inaccurate for all other sulfatide subclasses, whereas DeepCCS predicted all subclasses with a mean relative error of 8%. AllCCS2 yielded the most accurate prediction (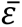of 1% for most subclasses, but 2.48% for SM2a; **Fig. 4ac; Supplementary Fig. 20**).

**Fig. 4.**
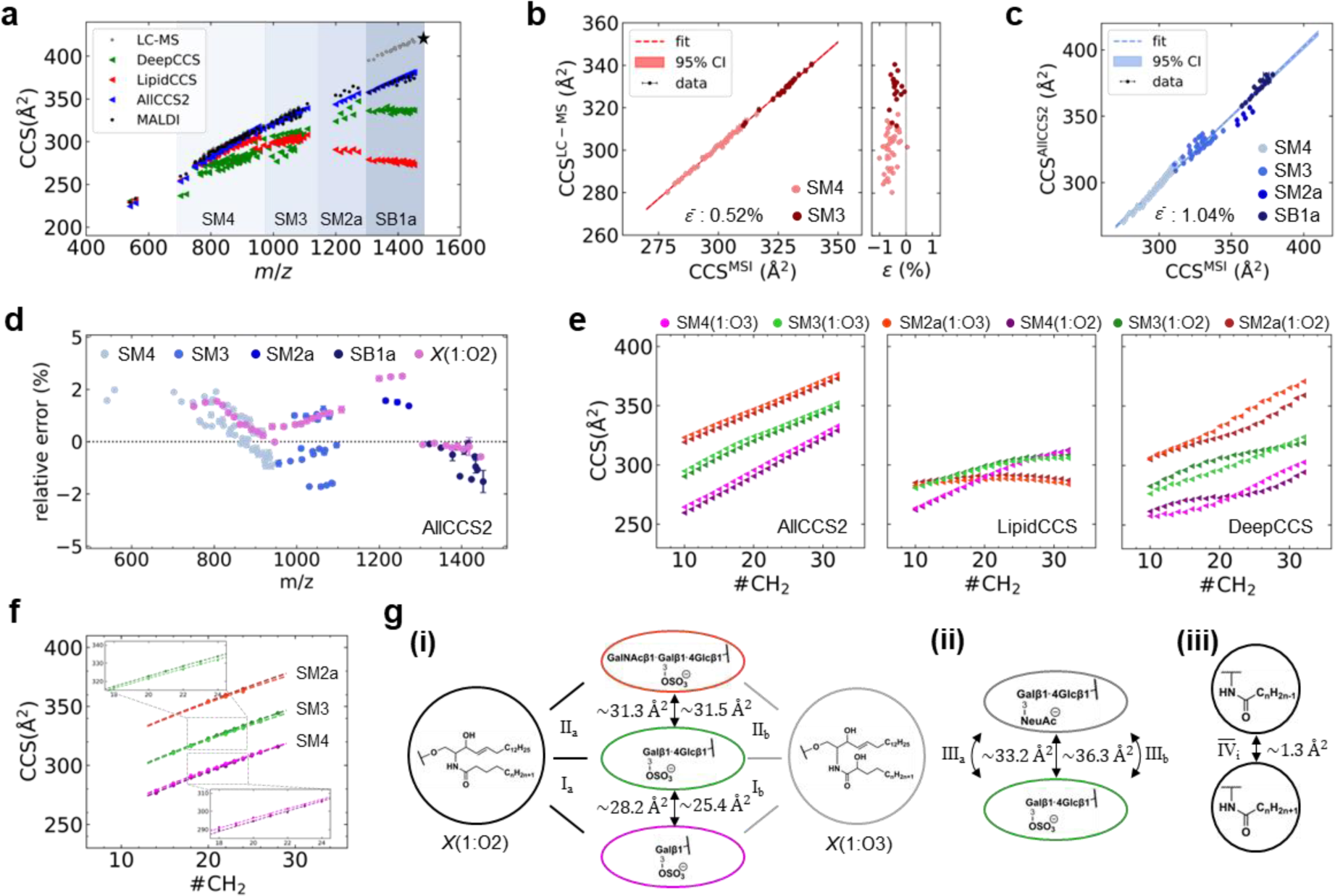
Structure-CCS-relationships for sulfatides. **a,** Comparison of experimental (gray dots: LC-TIMS-MS; black dots: MALDI-TIMS-MSI) and predicted CCS values, modeled by LipidCCS^57^ (red triangle), AllCCS2^30^ (blue triangle) and DeepCCS^32^ (green triangle). LC-MS-derived CCS values of SB1a[M-2H]^2-^ /SB1a[M-HSO_3_]^-^ are marked with a star. **b**, Strong correlation (R² = 0.9988; linear fit (red); 95% confidence interval (CI)) of LC-MS-derived and MALDI-MSI-derived ion mobility data. Mean relative error 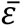= 0.5%. **c**, Correlation of experimental (MALDI-MSI) and predicted (AllCCS2) CCS values. Mean relative error 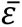 is 1.0%. **d**, Relative error *ε* reveals inconsistent deviations per subclass of the predicted CCS values using AllCCS2 against CCS values obtained by MALDI-MSI. **e**, CCS values of sulfatide subclasses as a function of fatty acid (FA) chain length for three different prediction tools, AllCCS2, LipidCCS and DeepCCS yielding ambiguous relative structural relationships. **f**, Experimental CCS values and 2^nd^ order polynomial fit to evaluate the contribution of the degree of glycosylation in the sulfated head group and the α-hydroxylation of the N-acyl FA. The relative positions are highlighted in the insets. **g**, Visualization of structural relationships for the data presented in **f (i)**, relative to ganglioside series **(ii)** and the contribution of saturated FA and mono-unsaturated FA. Surprisingly, the difference between SM3(O3) and SM4(O3) is reduced compared to the non-α-hydroxylated counterpart suggesting that interaction between the α-OH group and the head group may influence the three-dimensional structure.

Mean errors were neither a function of subclass nor of FA chain lengths (**Fig. 4d**), as highlighted by the inconsistent behavior of the relative error of the subclass *X*(1:O2) predicted by AllCCS2. These discrepancies between prediction and experiment, revealed by QCL-IRI-guided TIMS-MSI together with an ambiguous trend in the relative position of the homologous series of subclasses across various prediction tools (**Fig. 4e**), prompted us to analyze structure-CCS-relationships (SCR) in more detail.

To compare series of sulfatides in structural subclasses such as the α-hydroxylated (*X* 18+n:1;O3) and non-α-hydroxylated (*X* 18+n:1;O2) sulfatides, experimental CCS of a homologous series were better modeled by 2^nd^ order polynomial fits than by linear fits or *y* = *b*_*CCS*_*x*^2/3^ + *a*_*CCS*_ fits that have been used to describe the progression of CCS values of polymers^58^ (**Supplementary Fig. 21**). First, we assessed the contribution of glycosyl head groups and chain lengths of *N*-acyl-linked FA to experimental CCS values for both the α-hydroxylated (*X* 18+n:1;O3) and non-α-hydroxylated (*X* 18+n:1;O2) sulfatides compared to predicted CCS (**Fig. 4f**).

Only AllCCS2 correctly predicted the strictly monotonous increase with FA chain length, but it did not predict a subtle difference between the O2- and O3-subseries: In both MSI and LC-MS data (**Supplementary Fig. 22**), CCS_O2_>CCS_O3_ was observed for complex SM2a and SM3 sulfatides, but not for SM4, whereas inconsistent trends were observed for different prediction tools. Especially the most accurate predictor investigated here, AllCCS2, showed CCS_O3_>CCS_O2_ for all classes. Employing a parallel line model (see **Methods**), we determined constant differences ΔCCS between SM2a and SM3 isoforms of identical FA chain length of 31.3±0.2 Å^2^ and 31.5±0.5 Å^2^ for the O2- and O3-subseries, respectively (**Fig. 4f and 4g; Supplementary Fig. 21; Supplementary Table 15**). However, ΔCCS between SM3 and SM4 isoforms was 28.2±0.1 Å^2^ and 25.4±0.2 Å^2^ for the O2- and O3-subseries, respectively (**Fig. 4g(i)**).

This significant difference suggests that an interaction between the α-OH-group and the glycosyl head group may influence the three-dimensional structure. The finding is also supported by our LC-TIMS-MS data (**Supplementary Table 12**). This effect seems to be reversed for lipid classes with unrelated glycosyl head groups like sulfatides and GM3 gangliosides (**Fig. 4g(ii), Supplementary Fig. 23, Supplementary Table 15**). In contrast, single sites of FA unsaturation that, because of the cis-configuration of double bonds in FA, introduce a kink in their three-dimensional structure leading to a reduction of the CCS values by 1.3±0.2 Å^2^ **(Fig. 4g(iii)**, **Supplementary Fig. 24**). The position of the double bond was experimentally not determined. Our data indicated that phytosphingoid base-containing sulfatides exhibit the same trend as the SM3 and SM4 isoforms for the O2- and O3-subseries, *i.e.* that the relative difference in CCS is reduced for *X*(0:O4) compared to *X*(0:O3) (**Supplementary Fig. 25, Supplementary Table 15**).

In conclusion, with this study we make workflows and computational tools for QCL-IRI-guided high-performance MSI available for the scientific community. This versatile platform is applicable for many biomedical research topics ranging from single-cell metabolomics to 3D cell cultures and to well-defined functional tissue areas in disease research, and it offers enhanced bioanalytical depth while simultaneously reducing measurement time by focusing MSI on relevant ROIs. Consequently, it enables deep spatial profiling of odd-chain lipids, evaluation and improvement of prediction tools or investigation of structure-CCS-relationships. Genetically modified mice can serve as analytical ground truths in MSI, besides their established role as disease models. Altogether, the presented concepts pave the way for deeper spatial investigations of complex biological processes in general and of sulfatide biochemistry in particular with a high level of confidence in molecular identifications.

## Methods

### Materials

All chemicals and solvents were of HPLC-MS grade. Conductive indium tin oxide (ITO)-coated glass slides were purchased from Diamond Coatings (West Midlands, UK) or Bruker Daltonics ([ITO slides and MALDI IntelliSlides], Bremen, Germany). SuperFrost Plus Adhesion slides and BioGold Microarray Slides were obtained from Thermo Fisher Scientific (Schwerte, Germany). MALDI matrix 2,5-dihydroxyacetophenone (DHAP) was purchased from Thermo Fisher Scientific (Waltham, Massachusetts, USA), 1,5-diaminonaphtalene (DAN), and α-cyano-4-hydroxycinnamic acid (α-CHCA) from Merck KGaA (Darmstadt, Germany). Acetonitrile (ACN), ethanol (EtOH), LC-MS water, 2-propanol (IPA), ammonium sulfate (AmS) and glass cover slips were obtained from VWR Chemicals (Darmstadt, Germany). Hydroxypropyl methylcellulose: polyvinyl pyrrolidone (HPMC:PVP, 1:1, w/w) was prepared in-house as described elsewhere^59^.

For external calibration of the trap unit of the timsTOF fleX and timsTOF Pro mass spectrometer (Bruker Daltonics), ESI-L Low Concentration Tuning Mix (Agilent Technologies, Waldbronn, Germany) was used. Sulfatide standard C_17_ mono-sulfo galactosyl(β) ceramide d18:1/17:0 (SM4 35:1;O2), and ganglioside standard C18:0 GM3-d_5_ were purchased from Avanti Polar Lipids (Birmingham, USA). Trifluoroacetic acid (TFA), Mayer’s hemalum solution, hydrochloric acid, sodium bicarbonate, magnesium sulfate, eosin Y-solution 0.5%, xylene, and eukitt were purchased from Merck KGaA. Anti-pan-cytokeratin (panCK) and anti-vimentin antibodies labeled with photocleavable mass-tags (PC-MT) were purchased from AmberGen (Billerica, USA).

### Mouse studies

The arylsulfatase A (ARSA) mutant mouse line that was generated using an ARSA gene targeted embryonic stem cell clone^60^ has been described previously^28^. Here, mice that had been backcrossed with C57BL/6J mice for 12 generations were used. ARSA-deficient (ARSA-/-), heterozygous (ARSA+/-) and wild-type (ARSA+/+) mice were obtained from heterozygous breeding pairs. Mice aged 12 or 60 weeks were sacrificed by cervical dislocation, organs were removed and immediately frozen on dry ice. Frozen organs were stored at −80 °C and shipped on dry ice. Animal experiments (breeding and maintaining of ARSA-/- mice) were approved by the Landesamt für Natur, Umwelt und Verbraucherschutz Nordrhein-Westfalen (reference: 84-02.04. 2014.A117).

### 3D-cell culture models, tissue and spheroid slice preparation

Monoculture and biculture spheroids of CCD-1137Sk human fibroblasts and HT-29 human colon cancer cells (both LGC Standards, Wesel, Germany) were prepared, embedded, frozen and cut as described previously^44, 59^. Briefly, spheroids were harvested after 3 days and 12 hours after seeding at a density of 1 x 10^6^ cells/T75 flask for HT-29 cells and 1.5 x 10^6^ cells/T75 flask for CCD-1137Sk. All spheroids of the same type were collected in the same Eppendorf tube. Excess culture media was removed, and 1 mL of PBS was added for washing. For embedding, spheroids were transferred to HMPC-PVP filled channels inside a gelatin cryo-mold^59^. Fresh-frozen mouse kidneys and embedded spheroids (–80 °C) were sectioned at 10 and 20 μm thickness, respectively, with a Leica CM1950 cryostat (Leica Biosystems, Nussloch, Germany) at –18 °C chamber- and specimen head temperature. Sections were thaw-mounted onto ITO-coated glass slides from Diamond Coating or IntelliSlides and either stored at –80°C until further use or dried for a minimum of 15 min. in a desiccator. Dried ITO slides were put in a slide mailer and vacuum-sealed to avoid environmental influences on the samples.

### Quantum cascade laser (QCL) mid-infrared imaging microscopy

QCL mid-infrared imaging (IRI) microscopy was conducted on a Hyperion II-ILIM FT-IR and QCL microscope (Bruker Optics, Ettlingen, Germany) equipped with a 300 x 300 focal plane array detector and spatial coherence reduction technology ^43^. For rapid data acquisition of large specimens, a 3.5x (0.15 numerical aperture (NA)) objective was used, resulting in a nominal pixel size of 4.66 µm. In addition, high resolution images were recorded with either a 15x (0.4 NA) or a 20x (0.6 NA) objective, yielding nominal pixel sizes of 1.15 µm and 0.86 µm, respectively. The optical resolution of the instrument at wavenumber 1500 cm^-1^ can be estimated as 22.2 µm (0.15 NA), 8.3 µm (0.4 NA) and 5.5 µm (0.6 NA). All measurements were performed in reflection mode using ITO-slides (**Supplementary Fig. 1**). Prior to data acquisition, a background spectrum was collected on a clean part of the slide. Focus was adjusted manually.

#### QCL-IRI microscopy in single wavenumber mode

For image registration and teaching of the mass spectrometers, infrared data was recorded in *single wavenumber mode* at 1656 cm^-1^ (amide I band used for best contrast) for whole slide scans, and images were exported from the OPUS software v8.8 (Bruker Optics) via a python interface as a *.tiff* file.

#### QCL-IRI microscopy in sweep scan mode for tissue segmentation and definition of regions-of-interest (ROI)

For tissue segmentation, mid-IR hyperspectral imaging data of tissue specimens was recorded in *sweep scan mode* within a spectral range of 950 – 1800 cm^-1^, covered by four QCL modules, with a spectral sampling interval of 4 cm^-1^. Individual image tiles consist of 250 x 250 pixels. Hyperspectral data cubes were exported from the OPUS software v8.8 (Bruker Optics) via a python interface as *.pickle* file. Hyperspectral data was then imported into an in-house Python tool. Absorbance spectra underwent asymmetric least square smoothing to correct baseline distortions and light-scattering effects^12^. QCL IRI data, typically >> 10^6^ spectra per data set, was processed by pixelwise spectral differentiation (1^st^ or 2^nd^ order of absorbance or transmittance)^21^.

For visualization, spectra of the 2^nd^ derivative of the absorbance were interpolated using a cubic interpolation function. For simplicity, we consider the data obtained from spectral differentiation, usually given in units of 1/cm^-1^ (1^st^ derivative) and or 1/cm^-2^ (2^nd^ derivative), as unit-less.

#### Feature-selective image segmentation for definition of regions-of-interest (ROI)

A binary image created by a Gaussian Mixture Model (GGM; two clusters) on the amide peaks usually serves as mask that distinguishes tissue from background. For feature-selective image segmentation, in step1 use case-specific sets of wavenumbers were defined based on literature or comparison of 2^nd^ derivative data. In step2, these sets of wavenumbers were employed for *k*-means clustering to define ROIs. <u>Use case 1 –</u> <u>segmentation of brain regions in ARSA-/- mice.</u> For image segmentation the lipid-associated 1466 cm^-1^ and 1740 cm^-1^ (2^nd^ derivative of absorbance), were utilized, respectively. <u>Use case 2 - 3D cell biculture models:</u> Similarly, to distinguish CCD-1137Sk and HT-29 cell lines in biculture spheroids, the same bands at 1466 cm^-1^ and 1740 cm^-1^ were used (**Supplementary Fig. 5a**). For each monoculture spheroid an individual measurement area was used. Objects were recognized using *random walker* segmentation in Python v3.8 during final assignment of MSI measurement regions, in case multiple spheroids were not spatially separated^61^. <u>Use case 3 – kidney glomeruli</u> <u>in ARSA-/- mice:</u> For identification of glomeruli-containing tissue, blob-detection was performed on a mean QCL-IR image created from spectral features of the transmittance data centering around 1726 cm^-1^ and 1142 cm^-1^ (1^st^ derivative of transmittance) (**Supplementary Fig. 8c**). Initially, the cortex of the kidney was selected by a donut-shaped binary mask preserving the outline of the tissue region. Threshold filtering followed by size exclusion and eccentricity filtering yielded ROIs, the relevance of which was later confirmed by MSI (**Supplementary Fig. 8ab**). Typically, values for image thresholds were in the range of 30-60 (8-bit image), 40-60 and 1000 pixels for the lower and the upper limits, respectively, of the bandpass filter for size exclusion (3.5x objective, 4.66 µm pixel size) and 0.94 for eccentricity. <u>Use case 4 - kidneys’inner segment of outer medulla (ISOM) and inner medulla/papilla (IMP) in ARSA-/- mice:</u> QCL-IRI data was segmented based on characteristic lipid-associated spectral features^62^ representing the accumulation of sulfatide lipids. The following features were selected: 1466 cm^-1^ (CH_2_ bending vibration), 988 cm^-1^ (C_β_-O vibration of the 3- sulfogalactosyl head group), 1740 cm^-1^ (C=O stretching vibration) as well as the protein-associated features at 1548 cm^-1^ and 1656 cm^-1^ (amide II and I bands) (**Supplementary Fig. 5b**). Subsequently, unsupervised *k*-means clustering of tissue-specific regions was conducted on the masked imaged^12^. The number of clusters *k* was directed by calculation of the Calinski-Harabasz-Score implemented in the *yellowbrick* python package^63^.

#### Transfer of QCL-IRI-defined ROIs to mass spectrometer and teaching

MSI data acquisition was initially set up in flexImaging v5.0 (MR-MSI) or v7.2 (TIMS-MSI) (Bruker Daltonics), while using the whole-slide single 1656 cm^-1^ wavenumber IR reference image (absorbance) to teach the particular MS device. Hereby, the single-wavenumber image is modified by affine transformation and further used to generate a data acquisition file (*.mis*). The corresponding image at 1656 cm^-1^ (absorbance) from the hyperspectral cube is co-registered with the modified reference image by means of *SimpleITK*^64^ and *simple Elastix*^65^ yielding an affine transformation matrix, which is further used to transfer the ROI information into the frame of the modified image. Advanced Mattes mutual information was used as a similarity metric for with linear interpolation. QCL-IRI-defined ROIs were further processed in Python v3.8 to account for use case-specific demands, e.g. by removal of small holes (all use cases), size reduction (spheroids, ISOM and IMP) or expansion (glomeruli) by a layer of one or two MALDI pixels using respective functions from the *scikit-image* or by an in-house written iterative approach for hole opening to account for *donut-shaped* measurement regions (ISOM and IMP)^66^. Finally, ROIs were imported as polygonal areas into the *.mis* file.

#### Image co-registration for multimodal imaging

Image co-registration for multimodal data analysis was performed as previously described^67^. To estimate performance of segmentation/identification of small objects like glomeruli-containing tissue areas, we compared the number of identified objects in a multimodal approach, *i.e.* by comparing IRI and MSI alone versus combined IRI/MSI (**Supplementary Fig. S8**). Parameters for blob-detection were optimized to yield a high ratio between the numbers of objects identified in both IRI/MSI modalities vs. IRI alone. For identification of glomerular regions in MSI, we applied our previously described concept for spatial probabilistic mapping of metabolites^2^ to generate glomerular hotspots. For visualization purposes and incorporation of ROIs SCiLS Lab (Version 2024a Pro, Bruker Daltonics) and the SCiLS Lab API v6.3.115 has been used.

### MALDI-MR- and MALDI-TIMS-TOF Mass Spectrometry Imaging

#### Calibration

Prior to MSI data acquisition, external mass and ion mobility calibration was achieved (via ESI source) using ESI-Low Concentration Tuning Mix (Agilent Technologies, Santa Clara, USA) and a linear calibration model. Final mass calibration via the MALDI source was performed using red phosphorus (RedP) clusters P_n_ (n = 13–61 in intervals of 4) and either an enhanced-quadratic (timsTOF fleX) or quadratic (solariX) calibration model. During data acquisition, the internal standard (IS) SM4 35:1;O2 (C41H79NO11 S, [M-H]^-^; *m/z* 792.530107) was used for internal lock-mass calibration.

#### MALDI magnetic resonance MSI

Ultra-high resolving power data was acquired on a solariX 7T XR Fourier Transform Ion Cyclotron Resonance (FT-ICR) MS (Bruker Daltonics), equipped with a smartbeam II 2 kHz laser and ftms control 2.3.0 software (Bruker Daltonics, Build 92). Mass spectra were acquired in negative ion mode (*m/z* range 401.29–2600) and a time domain of acquisition of 8 M, resulting in a very long free induction decay (FID) of 15.7 s and a mass resolution of 1,230,000 at *m/z* 800. Reducing the number of data points in the time domain to 512k resulted in an FID of 0.98 s and a mass resolution of R ∼ 77,000. Ion optics settings were constant for all measurements: funnel RF amplitude (150 Vpp), source octopol (5 MHz, 350 Vpp), and collision cell voltage: 1.5 V, cell: 2 MHz, 1200 Vpp). The source DC optics was also constant for all measurements (capillary exit: - 200 V, deflector plate: −220 V, funnel 1: −150 V, skimmer 1: −15 V), as well as the ParaCell parameters (transfer exit lens: 30 V, analyzer entrance: 10 V, sidekick: 0 V, side kick offset: 1.5 V, front/back trap plate: −3.4 V, back trap plate quench: 30 V). Sweet excitation power for ion detection was set to 14 %, and ion accumulation time was 0.05 s. For ISOM and IMP, the transfer optics were as follows: time of flight: 1 ms, frequency: 4 MHz, and RF amplitude: 350 Vpp. The laser parameters were laser power: 32 %, laser shots: 20, and laser frequency: 200 Hz, and laser focus: medium, at a lateral step size of 40 µm. For glomeruli, the transfer optics were adjusted to 1.2 ms, 4 MHz, and 350 Vpp. The laser power was reduced to 28 % with minimum focus at a lateral step size of 20 µm.

#### MALDI trapped ion mobility spectrometry MSI

MALDI TIMS-MSI was carried out on a tims-TOF fleX system (Bruker Daltonics) equipped with a smartbeam 3D 10 kHz laser and TimsControl 4.1 and flexImaging v7.2 software (Bruker Daltonics). Data was acquired in negative ion mode (*m/z* range of 300–2000) with 240 laser shots per pixel, 2 kHz laser frequency and lateral step size 20 µm. The Ion Transfer parameters were as follows: MALDI Plate Offset 50 V, Deflection 1 Delta −70 V, Funnel 1 RF 350 Vpp, isCID Energy −0.0V, Funnel 2 RF 350 Vpp, and Multipole RF 320 Vpp. Collision Cell parameters: Collision Energy 8 eV, and Collision RF 1600 Vpp. Quadrupole parameters: Ion Energy 5 eV, and Low Mass *m/z* 320. Focus Pre TOF parameters: Transfer Time 85 µs, and Pre Pulse Storage 10 µs. For acquisition of full kidney data sets, the qTOF mode was utilized, and laser frequency was 10 kHz. TIMS-ON-MSI data was acquired in a range of 0.8–1.87 Vs/cm^2^, with ramp time 480 ms, and accumulation time 120 ms (resulting duty cycle 25 %). Tims-offsets: Δt1 (Deflection Transfer -> Capillary Exit): 20.0 V; Δt2 (Deflection Transfer -> Deflection Discard): 120.0 V; Δt3 (Funnel 1 In -> Deflection Transfer): −80.0 V; Δt4 (Accumulation Trap -> Funnel 1 in): −100.0 V; Δt5 (Accumulation Exit -> Accumulation Transfer): 0.0 V; Δt6 (Ramp Start -> Accumulation Exit): −100.0 V; Collision Cell Inlet: −225.0 V. TIMS-ON-MSI data of glomeruli was acquired at a lateral step size of 10 µm. Transfer time and pre-pulse storage were increased to 120 µs and 15 µs, respectively. The range was adjusted to 1.20–2.05 Vs/cm^2^, and offsets were as follows: Δt1 (Deflection Transfer -> Capillary Exit): 20.0 V; Δt2 (Deflection Transfer -> Deflection Discard): 120.0 V; Δt3 (Funnel 1 In -> Deflection Transfer): −85.0 V; Δt4 (Accumulation Trap -> Funnel 1 in): −150.0 V; Δt5 (Accumulation Exit -> Accumulation Transfer): 0.0 V; Δt6 (Ramp Start -> Accumulation Exit): −150.0 V; Collision Cell In: −225.0 V.

#### On-tissue lipid/metabolite fragmentation using prm-PASEF

Parallel reaction monitoring with parallel accumulation and serial fragmentation (prm-PASEF) of lipids/metabolites has so far only been outlined for TIMS-MS without spatial resolution^53^. Here, we used a prototypic version of Bruker software for spatially resolved prm-PASEF data acquisition in the *m/z* range 50–2000 with a quadrupole low mass of *m/z* 50, a collision cell energy of 4 eV and collision RF of 500 Vpp. The laser frequency was 5 kHz, the transfer and the pre-pulse storage times were 65 µs and 8 µs, respectively for kidney ISOM and IMP, and 70 µs and 10 µs, respectively, for glomeruli. Fragmentation energies were modulated for kidney ISOM and IMP (for 1/k_0_ = 1.25 Vs/cm^2^, collision energy 65 eV; 1.30 and 72; 1.35 and 80; 1.40 and 85; 1.45 and 95; 1.60 and 100) and also for glomeruli (1.20 and 55; 1.70 and 75; 1.80 and 80).

#### Multiplex-MALDI-MS-Immunohistochemistry (IHC)

using two PC-MT antibody probes (pan-cytokeratin [panCK] and vimentin) was carried out as described^45^, with the following modifications for fresh-frozen spheroid sections: 2x 3 min ice-cold acetone, 30 min 1% paraformaldehyde fixation; 10 min PBS; 2x 3 min acetone; 3 min Carnoy’s solution; 2x 2 min 100% ethanol, 3 min 95% ethanol, 3 min 70% ethanol, 3 min 50% ethanol and 10 min TBS. Antigen retrieval (100x Tris-EDTA Buffer, pH 9) was performed for 30 min using a water bath at 95 °C in a coplin jar (VWR Chemicals). Spheroid sections were blocked with Tissue Blocking Buffer (2% [*v/v*] normal mouse serum and 5% (*w/v*) BSA in TBS-octyl-beta-glucoside (OBG; 0.05% [*w/v*]). For the PC-MT antibody treatment, the slide was incubated with 3 µg/mL of panCK and 2 µg/mL of vimentin each antibody diluted in Tissue Blocking Buffer at 4 °C overnight in a humidified, light-protected environment. Next, the slide was washed with 3x 5 min TBS; 3x 2 min 50 mM ammonium bicarbonate for 2 min each, and dried in a vacuum desiccator for 2 hours. Probes were photo-cleaved at 365 nm for 10 min in a UV illumination box (Ambergen). Then α-CHCA matrix (10 mg/mL in 70% ACN with 0.1% TFA) was applied using a HTX TM sprayer (HTX Technologies) with the following parameters: flow rate=0.1 mL/min, velocity=1,350 mm/min, number of passes=8, nozzle height=40 mm, pattern CC, temperature=60 °C, nitrogen gas pressure 10 psi; followed by recrystallization at 55 ⁰C with 1.0 mL 5% isopropyl alcohol in LC-MS water for 1 min. Data was acquired using a timsTOF fleX mass spectrometer. PC-MTs were analyzed in qTOF mode with the following ion transfer parameters: MALDI Plate Offset 30 V, Deflection 1 Delta 80 V, Funnel 1 RF 500 Vpp, isCID Energy −0.0V, Funnel 2 RF 500 Vpp, and Multipole RF 1200 Vpp. The Collision Cell parameters were: Collision Energy 25 eV, and Collision RF 4000 Vpp. The quadrupole parameters were: Ion Energy 15 eV, and Low Mass *m/z* 900. The Focus Pre TOF parameters were: Transfer Time 140 µs, and Pre Pulse Storage 10 µs.

### Analysis of MALDI-MS imaging data

#### Selection of hypothetical sulfatide configurations

Based on previous empirical data^28^, we considered the fatty acid compositions from 32:x to 46:x, with x = 0,1,2,3 for O2,O3 and x = 0,1 for O4, and six lyso-sulfatide compositions as potentially detectable with our MSI settings, resulting in a total number of 156 theoretical compositions for each sulfatide subclass.

#### MR-MSI high resolution metabolic profiling

Data was saved as profile spectrum with a data reduction factor of 97% in addition to the centroided mass spectra (SQLite peaks list) generated during data acquisition. Data was imported into R using a custom function reading the SQLite peak list. It was converted into *MALDIquant* MassPeaks-Objects^68^. For each reference mass list entry, the closest peak (1 ppm tolerance) per spectrum was selected and the relative mass deviation of measured and theoretical masses was computed. To determine region-wise mass deviation, a Gaussian distribution was fitted to the mass deviation using *fitdistrplus* package^69^, and the mean value and standard deviation of the fitted distribution were reported. To determine the global mass deviation across the *m/z*-axis, the pixel-wise mass deviation values were filtered to exclude observations outside mean ±3* standard deviations before again determining the mean (corrected mass deviation). Molecular probabilistic maps (MPMs) of metabolite distributions were calculated and plotted by the R package *moleculaR*^2^. For visualization of high resolution MR-MSI data (**Fig. 2g**), centroided data was exported from Data Analysis v6.1 (Bruker Daltonics) including peak resolution information and modelled by a Gaussian distribution. For annotating MR-MSI data via Metaspace, data was RMS-normalized, exported from SCiLS Lab (Version 2024a Pro, Bruker Daltonics) as imzML file, uploaded to Metaspace (www.metaspace2020.eu) and annotated using the following settings: *m/z* tolerance 3 ppm, Analysis Version v2.20230517 (META-SPACE ML https://www.biorxiv.org/content/10.1101/2023.05.29.542736v1), database SwissLipids - 2018-02-02. Annotations were downloaded and visualized using R/Python. Hereby, annotations were filtered for molecules containing a sulfate group in a first step and compared against experimentally identified sulfatides.

#### Classification of sulfatide peaks in qTOF mode

Each sulfatide peak that showed at least one additional peak within a mass window of ±1.1 Da that exceeded the intensity of the first sulfatide isotope peak was classified as “non-clean”, since this “interfering” peak would prevent an unequivocal assignment of an on-tissue MS^2^ spectrum (**Fig. 3a**). Sulfatide peaks without interfering peaks were classified as “clean” (**Supplementary Fig. 13**). *Classification of sulfatide peaks in TIMS-ON mode:* Each sulfatide peak that shows a colocalizing peak within a mass window of ±1.1 Da and a mobility window of ±0.005 Vs/cm² that exceeds the intensity of the first sulfatide isotope peak was classified as non-clean, since this interfering peak would prevent a clear assignment of a prm-PASEF MS^2^ spectrum. Sulfatide peaks without interfering peaks were classified as clean (**Fig. 3b**). *timsTOF* fleX *TIMS-ON mode:* CCS values obtained from QCL IRI-guided MSI of four biological replicates were averaged and the standard deviation was calculated and given as uncertainty. To obtain CCS values from prediction tools AllCCS2^30^, DeepCCS^32^ and LipidCCS^57^, simplified molecular input line entry specification (SMILES) strings were provided to these tools. All chemical structures and all SMILES strings were generated in ChemDraw 21.0.0.38 (PerkinElmer, Waltham, US). To benchmark the predicted values CCS^pred.^ against experimental values CCS^MSI^, the mean relative error 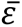 in % was calculated by

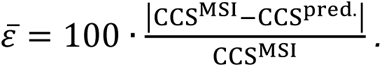

For the evaluation of relative differences in CCS values, *e.g.,* of the degree of glycosylation in the sulfated head group and the α-hydroxylation of the *N*-acyl-linked FA in sulfoglycosphingolipids, a global least square model fitting was applied. Hereby, a 2^nd^ order polynomial fit was performed by keeping the amplitudes of the linear and quadratic term fixed, parallel line model, when describing the data for two (or three) similar sulfatide subclasses, *e.g.,* SM3 18+n:1;O2 and SM4 18+n:1;O2. On the level of our experimental accuracy, we found that this parallel line model can be applied to describe the data for similar subgroups, enabling us to deduce relative contributions of chemical modifications to CCS values.

### LC-TIMS-TOF-MS

#### Sulfatides extraction from ARSA-/- kidney tissues

Two 12-week-old and two 60-week-old ARSA-/-, two heterozygous (ARSA+/-) and two wild-type (ARSA+/+) kidney samples underwent two rounds of homogenization for 30 seconds each using 400 μL of 200 mM Na_2_CO_3_ (pH=9.3). Subsequently, all samples were subjected to overnight freeze-drying at −56 ^°^C and 1 mbar.

The extraction of kidney tissues was carried out utilizing a modified Folch method. Kidney tissues, spiked with 5 μL GM3-d5 (100 μg/ml in methanol) and 5 μL SM4 35:1;O2 (100 μg/mL in chloroform/methanol, 2:1[*v/v*]) as internal standards, were mixed and sonicated with 9 mL of chloroform/methanol (2:1[*v/v*]) for 15 min. Subsequently, 840 μL of water were added, and the resulting mixture was thoroughly mixed before being centrifuged at 3000 rpm for 5 min at RT. The chloroform layer at the bottom was collected and evaporated using a stream of nitrogen. Finally, the samples were re-dissolved in 200 μL methanol/water (4:1[*v/v*])

#### LC-TIMS-MS

4D LC-TIMS-MS experiments were conducted using timsTOF PRO instrument interfaced with an Elute UHPLC system (both Bruker Daltonics). Analysis of sulfatides was performed in negative ion mode. The LC-TIMS-MS method was adapted from the protocol of Lerner *et al.* (2023)^47^ with modifications. The LC column was a C_18_ Kinetex column (100 x 2.1 mm x 2.6 μm) (Phenomenex, Germany). The PASEF scan mode was performed on a mass scan range of *m/z* 200–2000 Da for both MS and MS^2^ acquisition. The collision energy was 90 eV.

#### LC-TIMS-MS data analysis

Data processing including identification and annotation of the sulfatides was performed using Metaboscape 2021b (Bruker Daltonics). The acquired data from ARSA-/- and ARSA+/+ (ARSA +/- for 12-week-old mice) samples was subsequently uploaded to Metaboscape 2021b in a single bucket table. The processing method parameters included: *i)* feature detection was performed using an intensity threshold of 100 counts, *ii)* recursive feature extraction was conducted for features found in at least 1 out of 8 analyses (to ensure features are found in both or either ARSA-/- and ARSA+/+ (ARSA +/- for 12-week-mice) samples), and *iii)* the feature was only included in the bucket tables if it was found in 1 analysis after recursive feature extraction. The established Sulfatides Analyte list containing 4D-descriptors (*m/z*, retention time (R_T_), CCS, and MS^2^ spectra) was created by manual structural annotation and elucidation and further used for sulfatide automated annotation. Tolerances and scoring adjustments for automatically annotating sulfatides using Sulfatides List include *m/z* between 2 and 5 mDa, R_T_ between 0.1 and 0.5 min, mSigma between 10 and 500, MS/MS score between 500 and 900, and CCS between 0.1% and 2.0%. After manual verification and further curation, we identified 93–95 sulfatide species in two 60-week-old and 44-65 sulfatide species in two 12 week-old ARSA-/- kidney samples.

## Data availability

Extensive data (4D-LC-TIMS-MS; MR-MSI data; TIMS-MSI-MS and MS^2^ data incl fragment spectra and images) will be made publicly available in line with requirements by the accepting journal.

## Code availability

Code will be made publicly available in line with requirements by the accepting journal.

## Acknowledgements

We thank Lucas Gast for performing automated H&E staining, Marten Seeba and Nils Kröger-Lui (Bruker Optics) for introduction to the Hyperion II ILIM instrument and computational support, Ethan Yang (Bruker Daltonics) for fruitful discussions, Hans-Christian Koch (Bruker Optics) for access to the Hyperion II instrument, and Arne Fütterer (Bruker Daltonics) for access to prm-PASEF prototype software. C.H. acknowledges support by the Ministerium für Wissenschaft, Forschung & Kunst (MWK) Baden-Württemberg „Mittelbauprogramm“. This work was supported by the BMBF (German Federal Ministry of Research) as part of the Innovation Partnership “Multimodal Analytics and Intelligent Sensorics for the Health Industry” (M^2^Aind), projects “Drugs4Future” (grant 12FH8I05IA) and “DrugsData” (grant 13FH8I09IA) to C.H. and R.R., within the framework FH-Impuls. Acquisition of the solarix 7T XR was supported by DFG (Project 262133997) to C.H. Acquisition of the timsTOF flex mass spectrometer was supported by BMBF as part of the MSCorSys SMART-CARE (grant 161L0212F) to CH.

## Author contributions

L.G. and S.S. designed and conducted all QCL-IRI experiments, designed all MSI experiments, performed ion mobility analysis and generated the Figures. L.G. performed all MSI experiments. S.S. wrote python code for multimodal image registration and for QCL-IRI data analysis and analyzed MR-MSI and IRI data. L.G. conducted all MSI experiments, analyzed the data and interpreted all MSI fragmentation spectra for ground truth evaluations. T.E. analyzed high-resolution MR-MSI data, wrote R code and performed MSI feature identification. D.A.S. conducted generation of molecular probabilistic maps for multimodal analysis and benchmark of glomerular structures. J.L.C. performed statistical analysis and provided input for ion mobility analysis. H.G.V. and L.B. conducted and analyzed all LC-TIMS-MS experiments and generated corresponding tables. F.K. and R.R. performed 3D cell culture experiments and analyzed data. S.A.I. performed MSI 3D cell culture experiments and analyzed data. Y.U. performed multiplex-MALDI-IHC experiments and analyzed corresponding MSI data. R.R., L.B. and C.H. provided infrastructure. M.E. genotyped and provided ARSA-/- mouse organs and insights into sulfatide biochemistry. C.H. conceived and managed the overall study and wrote the first draft of the manuscript. L.G., S.S. and C.H. wrote the final manuscript – with input from all co-authors.

