## Supplementary Information for "Deep MALDI-MS Spatial ‘Omics guided by Quantum Cascade Laser Mid-infrared Imaging Microscopy"

+ these authors contributed equally.

Supplementary Figure 21. Modelling of experimental CCS values as a function of the chain length of the *N*-acyl-linked fatty acid (FA). 24
Supplementary Figure 22. Experimental (MALDI-TIMS-MSI and LC-MS) and predicted CCS values as a function of *N*-acyl-linked fatty acid chain length: 25
Supplementary Figure 23. Comparison of experimental MALDI-TIMS-MSI CCS values for SM3 and GM3 subclasses. 25
Supplementary Figure 24. Evolution of the relative CCS values between selected SM3 and SM4 subclasses incorporating either a saturated FA ($C_nH_{2n+1}$), or a mono-unsaturated FA ($C_nH_{2n-1}$). 26
Supplementary Figure 25. Evolution of the relative CCS values between selected SM3 and SM4 subclasses. 27
**Supplementary Tables** 27
Supplementary Table 1. Overview of significant features (as determined by lasso method) in MCF and BCF. 27
Supplementary Table 2. Identified fragment ions of lyso-PI 18:0$[M-H]^-$ (*m/z* 599.317) 28
Supplementary Table 3. Identified fragment ions of PE 32:1$[M-H]^-$ (*m/z* 688.489). 28
Supplementary Table 4. Identified fragment ions of PE P-36:4$[M-H]^-$ (*m/z* 722.511) 28
Supplementary Table 5. Identified fragment ions of PA 38:2$[M-H]^-$ (*m/z* 727.523). 29
Supplementary Table 6. Identified fragment ions of PI 33:1$[M-H]^-$ (*m/z* 821.517). 29
Supplementary Table 7. Identified fragment ions of PI 34:1$[M-H]^-$ (3$^{rd}$ carbon isotope ($^{13}C_3$), *m/z* 838.541). 30
Supplementary Table 8. Identified fragment ions of PI 36:3[M-H]- (m/z 859.530) 31
Supplementary Table 9. Identified fragment ions of PI 34:1$[M-H]^-$ (2$^{nd}$ carbon isotope ($^{13}C_2$), *m/z* 915.593). 32
Supplementary Table 10. Overview of GM3-series gangliosides identified in ARSA-/- and ARSA+/+ kidney**.** 32
Supplementary Table 11. List of all (kidney) sulfatides reported in ARSA-/- mice based on low-resolution MALDI-TOF-MSI (Marsching et al., 2011)[5]. 33
Supplementary Table 12. Overview of sulfatides identified in ARSA-/- kidney by LC-ESI-TIMS-TOF MS. 35
Supplementary Table 13. Overview of sulfatides identified in ARSA-/- kidney by MALDI-MSI. 37
Supplementary Table 14. Prm-PASEF fragmentation patterns of sulfatide isoforms identified by MALDI-timsTOF-MSI. Color code represents the degree of hydroxylation. 39
Supplementary Table 15. Relative values for structure-CCS-relationships of sulfatide subclasses. 40
**References** 41

### Supplementary Methods

#### Haematoxylin & Eosin Staining

Matrix removal was performed by submerging the slide in 70 % EtOH for 1 min. For subsequent H&E staining, the SunTissuePrep System (SunChrom, Friedrichsdorf, Germany) was used. The staining procedure consists of the following steps as described previously[1]: 1.5 min hemalum (Mayer's hemalum solution, Sigma-Aldrich), 2 min tap water, 1 min deionized water, 1 min acidic (350 mL ethanol + 150 mL $H_2O$ + 1.5 mL HCl) ethanol (absolute EMPLURA®, Merck; HCL Titripur, Merck), 45 s deionized water, 2 min bluing solution (2 g $NaHCO_3$ + 20 g $MgSO_4$ in 1L $H_2O$) ($NaHCO_3$, Merck) ($MgSO_4$, VWR Chemicals), 1 min deionized water, 2 min Eosin (Eosin Y-solution 0.5 % aqueous, Merck), 45 s deionized water, 1 min 80 % EtOH, 2 min 96 % EtOH, 2 min 100 % EtOH, and 3 min xylene. The slides were cover-slipped with Eukitt (Sigma-Aldrich) and a glass cover slide (VWR Chemicals) for long-term preservation.

#### MALDI-FT-ICR- and MALDI-TIMS-TOF Mass Spectrometry Imaging

##### *Matrix spray-coating*

10 mg/mL DHAP was dissolved in 70% ACN with 125 mM ammonium sulfate. After sonication, 0.1% TFA and 3 µM of SM4 35:1;O2 (100 µg/mL (=157.41 µM) in MeOH/chloroform 2:1) as internal standard (IS) were added. Matrix was applied with an M5 TM-Sprayer (HTX Technologies, Chapel Hill, USA). Temperatures of the spray nozzle and tray were 75 °C and 35 °C, respectively. The spraying parameters were as follows: Spray Nozzle Velocity: 1200 mm/min; Flow Rate: 0.1 mL/min; No. of Passes: 10; Track Spacing: 2 mm; Pattern: HH; Pressure: 10 psi; Gas Low Rate: 2 L/min; Nozzle Height: 40 mm; Drying Time: 0s. Dissected spheroids were coated with DAN (10 mg/mL) in ACN/water 7:3 (*v/v*). The spraying parameters were as follows: Spray Nozzle Velocity: 1350 mm/min; Flow Rate: 0.07 mL/min; No. of Passes: 6; Track Spacing: 2 mm; Pattern: HH; Pressure: 10 psi; Gas Low Rate: 2 L/min; Nozzle Height: 40 mm; Drying Time: 10s. For the specified measurements of PC-MTs, the slides were spray-coated with α-CHCA (10 mg/mL) in ACN/water/TFA 7:3:0.01 (*v/v/v*) with the following parameters: Spray Nozzle Velocity:

1350 mm/min; Flow Rate: 0.1 mL/min; No. of Passes: 8; Pattern: HH; Pressure: 10 psi; Gas Low Rate: 2 L/min; Nozzle Height: 40 mm; Drying Time: 10s.

### Supplementary Figures

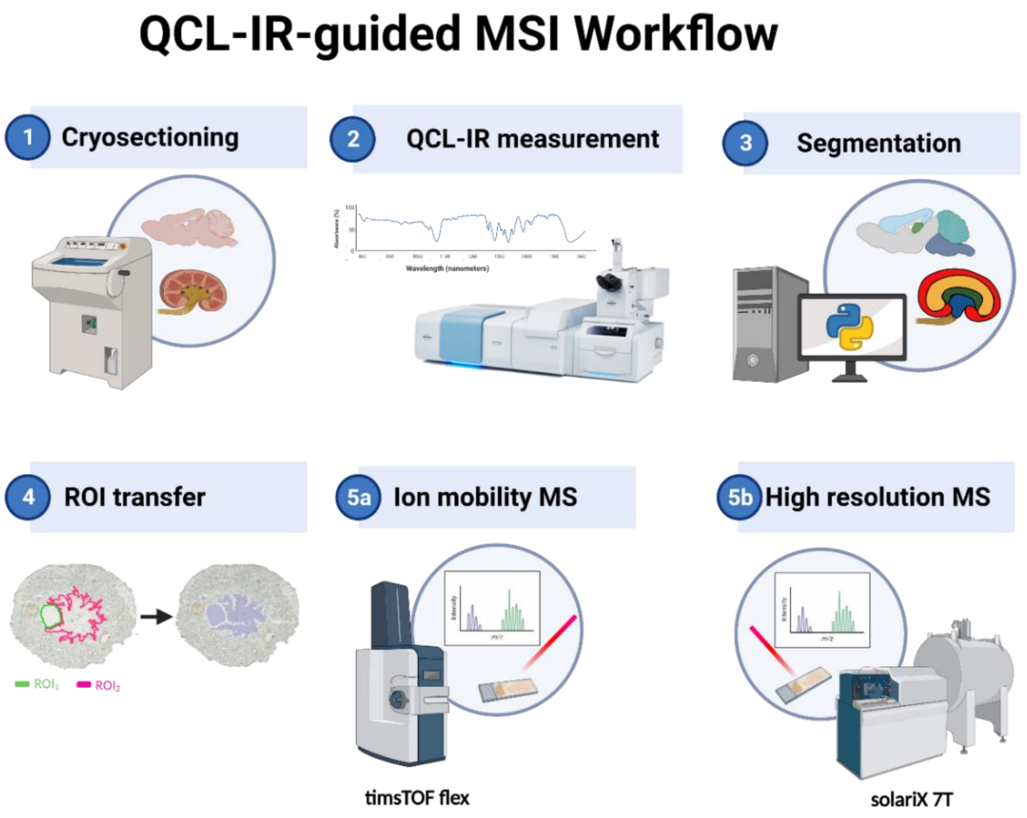

Supplementary Fig. 1. Schematic workflow of QCL Mid-infrared Imaging Microscopy guided MALDI MS Imaging (QCL-IRI guided MSI). **1**, Fresh-frozen tissue is cut and mounted onto indium tin oxide (ITO)-coated glass slides and dried in a desiccator. **2**, Mid-IR spectra covering the full "fingerprint" region (950–1800 cm$^{-1}$) are rapidly recorded using a quantum cascade laser (QCL) mid-IR microscope in sweep-scan mode. **3**, QCL-IRI datasets are segmented utilizing unsupervised methods like k-means clustering on most distinctive wavenumbers or wavenumber bands. **4**, Segments are identified as regions of interest (ROIs) and co-registered with a reference whole-slide, *single wavenumber* infrared image. **5a**, ROI information is transferred to the data acquisition file of either a trapped ion mobility spectrometry - time of flight (timsTOF; left) or **5b,** a magnetic resonance (MR) Fourier transform-ion cyclotron resonance (FT-ICR) mass spectrometer, in order to restrict MSI to ROIs defined by QCL-IRI. This figure was created with BioRender.com.

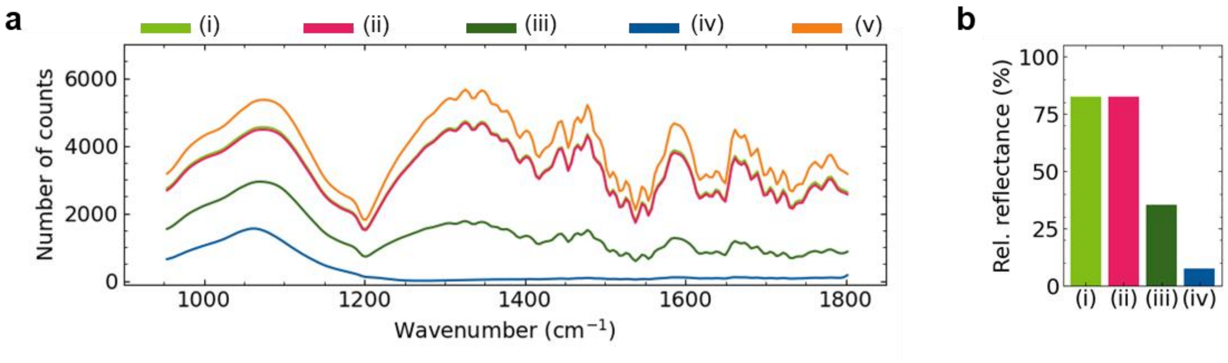

Supplementary Fig. 2. Mid-infrared (MIR) optical properties of various surface-coated glass slides. **a,** The relative reflectance across the mid-infrared fingerprint region was measured as the number of counts on the detector from a single channel measurement on the Hyperion II QCL-IRI microscope for various ITO-coatings (i), (ii) and (iii), for normal glass slide (iv), and for a gold-coated surface (v). **b,** Spectral responses for ITO-coated slides (relative to gold coating (v)) were averaged over the entire spectra ranging from 950-1800 $cm^{-1}$. The mean relative reflectance ranged from 82% for both the Bruker MALDI IntelliSlide (i) and the Diamond Coating ITO-coated glass slide (ii) to 35% for the Bruker ITO-coated glass slide (iii) to 7% for the non-coated SuperFrost Plus Adhesion glass slide. As a result, for biomedical specimen analysis with the presented workflows, Diamond Coating ITO-coated slides (ii) were mainly used, since they allow for transmitted-light microscopy and visual inspection during sample preparation. All mass spectrometry-related methods and protocols were optimized for this slide type.

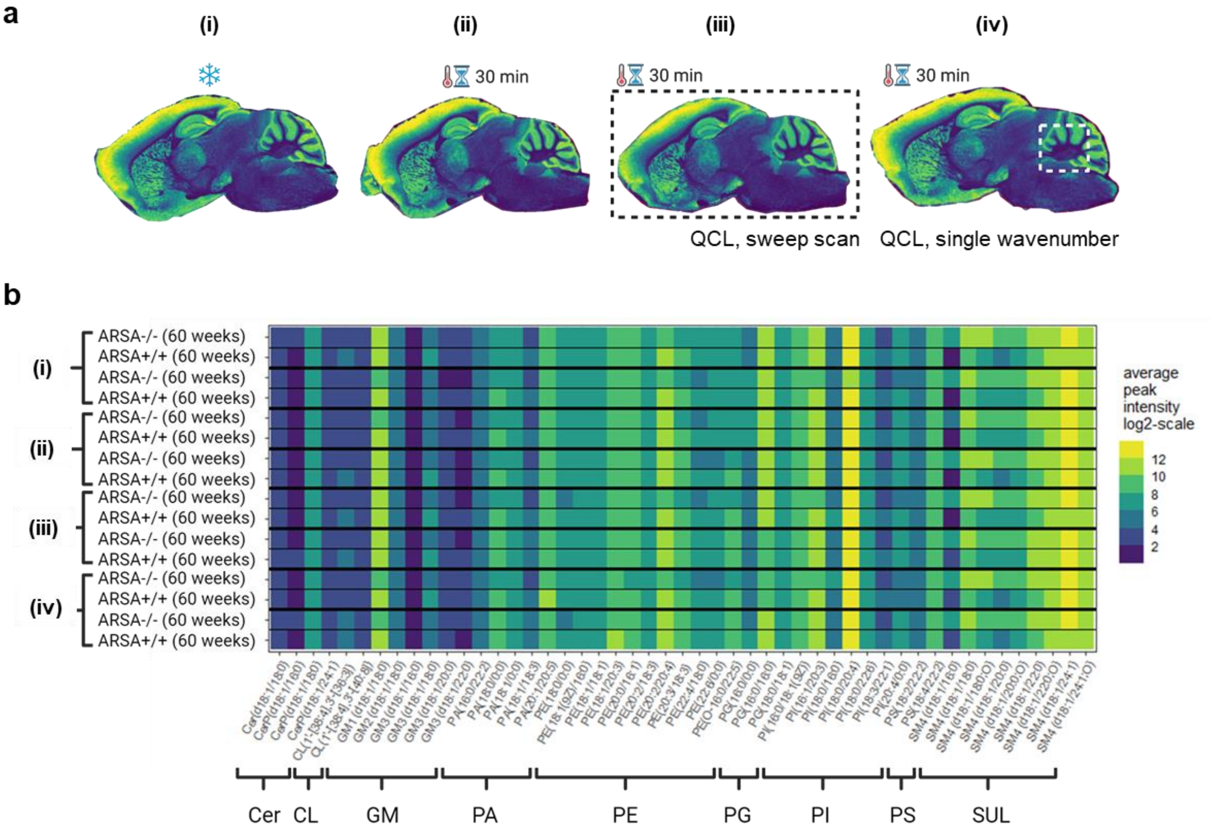

**Supplementary Fig. 3. MSI assessment of lipid alterations induced in mouse brain cryo-sections by pre-analytical stress or by laser light.** **a,** Four different stress conditions prior to MALDI MSI (timsTOF fleX operated in qTOF mode) were investigated: **(i)** standard workflow where the samples are stored at -80 °C after sectioning and before matrix deposition, **(ii)** the specimen is kept at room temperature (RT) and standard pressure (SP) for about 30 min, **(iii)** the entire tissue section (dashed black box) is exposed to QCL-IRI scanning for 15 min in sweep scan mode and kept at RT and SP for in total of about 30 min, and **(iv)** where a defined 1.2 mm x 1.2 mm region of the tissue sections (dashed white box) is exposed to infrared light for 15 min at a constant wavenumber of 1656 $cm^{-1}$ (amide I) used for generation of the reference image. **b,** Average peak intensity in negative ion mode MALDI MSI for several lipid classes across an *m/z* range of 600 - 1700. For each of the stress conditions **(i)** to **(iv)** in **a**, the procedure was repeated for four different 60-week-old mice, two wild-type (ARSA+/+) mice and two arylsulfatase A-deficient (ARSA-/-) mice. Lipid assignment was done by *m/z*-based annotation in Metaspace (www.metaspace2020.eu). No indication of environment- or laser-induced lipid alterations was observed in the MSI data. Differences in peak intensity were within the expected range, considering biological variability and known batch effects in MALDI MSI[2]. Abbreviations: Cer: Ceramides, CL: cardiolipin, GM: ganglioside, PA: phosphatidic acid, PE: phosphatidylethanolamine, PG: phosphatidylglycerol, PI: phosphatidylinositol, PS: phosphatidylserine and SUL: sulfatide.

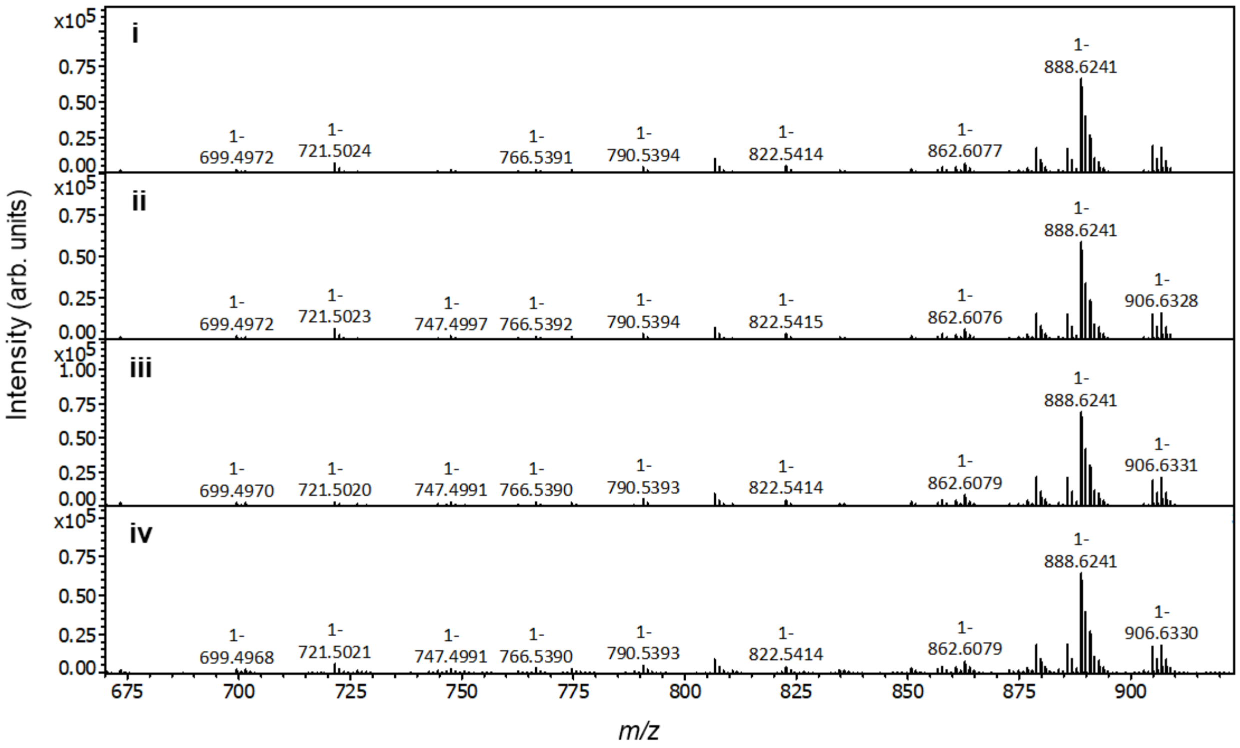

Supplementary Fig. 4. Lipid MS profiles of stressed brain tissue sections. Representative MALDI-qTOF-MSI (timsTOF fleX) average spectra (*m/z* 675–920) for brain slices of 60 weeks-old ARSA-/- mice following treatment under conditions presented in **Suppl. Fig. 3**. **(i)** standard workflow where the samples are stored at -80 °C after sectioning and before matrix deposition, **(ii)** the specimen is kept at room temperature (RT) and standard pressure (SP) for about 30 min, **(iii)** the entire tissue section (dashed black box) is exposed to QCL-IR scanning for 15 min in sweep scan mode and kept at RT and SP for in total of about 30 min, and **(iv)** where a defined 1.2 mm x 1.2 mm region of the tissue sections (dashed white box) is exposed to infrared light for 15 min at a constant wavenumber of 1656 $cm^{-1}$. Intensity scale is identical for **(i–iv)**. No lipid alterations were observed.

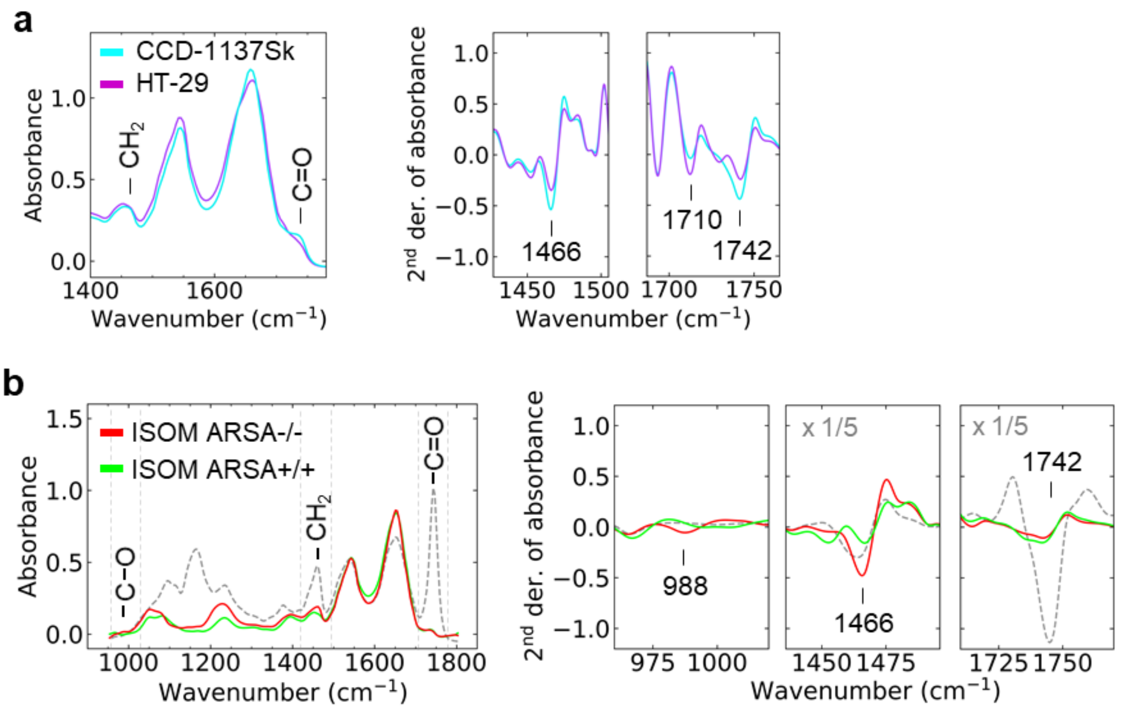

Supplementary Fig. 5. Infrared spectral data for biculture spheroids and ARSA-/- mouse kidney. **a,** Mean cell type-specific QCL-IR absorbance spectra (Hyperion II, 20x objective) for CCD-1137Sk fibroblasts and HT-29 colon cancer cells in spheroids. Lipid-associated bands at 1466 $cm^{-1}$ and 1740 $cm^{-1}$ are reduced in HT-29 cells. 2nd derivative of the mean absorbance spectra for 1466 $cm^{-1}$ and 1740 $cm^{-1}$ demonstrates that the transition (1710 $cm^{-1}$) between the amide I and ester bands is discriminative for the two cell lines. **b**, Mean absorbance spectra and 2nd derivative of absorbance (Hyperion II, 3.5x objective) of the ISOM and IMP region as in **Fig.1i and j**. In addition, the grey dotted line corresponds to the absorbance spectra of a tissue region of high fat content present within the kidneys showing partial overlap with discriminant features of the sulfatide fingerprint, e.g. at 1466 $cm^{-1}$. Dashed vertical lines represent the spectra regions highlighted for the 2nd derivative data.

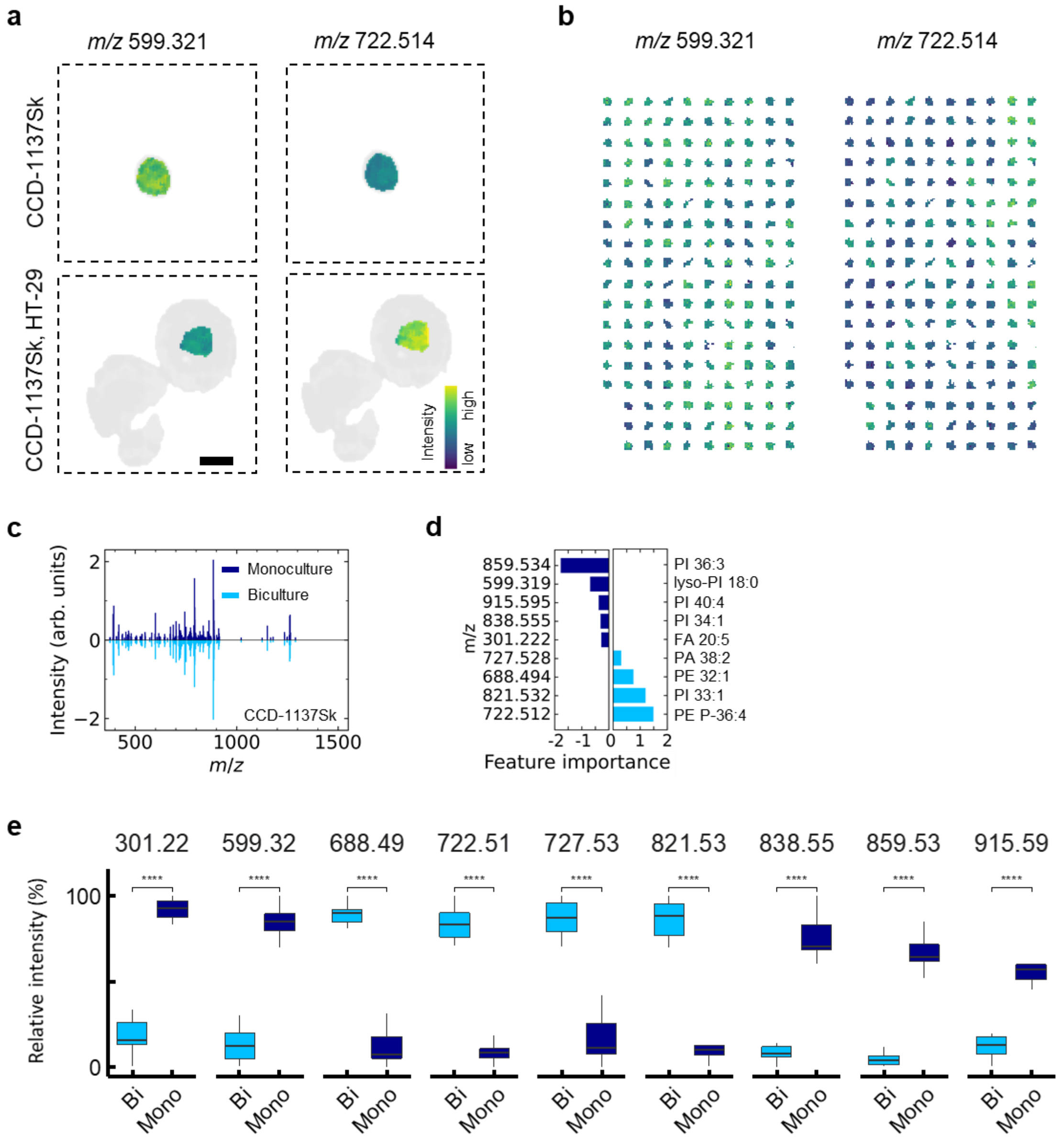

Supplementary Fig. 6. Profiling of fibroblasts in mono- and biculture spheroids. **a**, QCL-IRI-guided TIMS-MSI-derived ion images (timsTOF fleX) for *m/z* 599.321 (lyso-PI 18:0[M-H]$^-$) and *m/z* 722.514 (PE P-36:4[M-H]$^-$; both within a ±10 ppm mass window) in monoculture fibroblast (MCF) CCD-1137Sk spheroids and the core fibroblast region of a biculture spheroid (BCF) containing of CCD-1137Sk and HT-29 colon cancer cells. Both *m/z* values are part of a discriminative feature list for monoculture versus biculture fibroblasts. Scale bar, 200 µm. **b**, Overview of ion images for *m/z* 599.321 (lyso-PI 18:0[M-H]$^-$) and *m/z* 722.514 (PE P-36:4[M-H]$^-$) from CCD-1137Sk cells of 105 MCF- and 72 BCF spheroid technical replicates. **c**, Butterfly plot of the mean intensity from 105 MCF- and 72 BCF spheroids**. d,** Machine learning-based feature extraction (LASSO (Least Absolute Shrinkage and Selection Operator) regression). Feature importance reveals discriminative *m/z* values between mono- and biculture fibroblasts. **e,** Boxplots of relative signal intensities for the extracted *m/z* features. ****Benjamini-Hochberg-adjusted p-value of < 0.001.

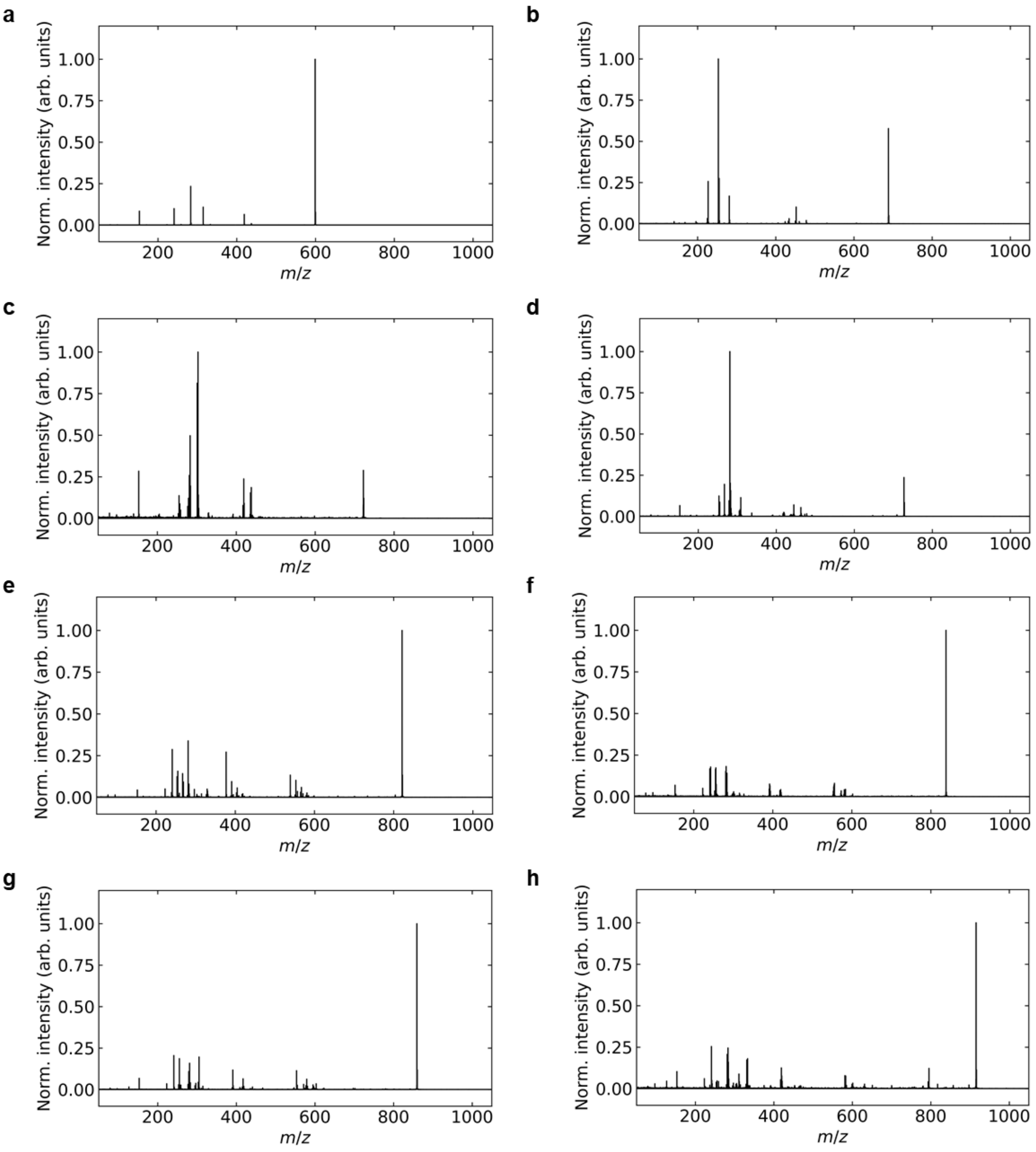

**Supplementary Fig. 7. QCL-IRI-guided partial reaction monitoring with parallel accumulation serial fragmentation (prm-PASEF)-MS$^2$ spectra of *m/z* features discriminating between MCF and BCF.** All MS$^2$ spectra were recorded on timsTOF flex. **a,** *m/z* 599.317 (lyso-PI 18:0[M-H]$^-$) isolated at $1/K_0$ = 1.171 Vs/cm$^2$ and fragmented with -40.0 eV. **b,** *m/z* 688.489 (PE 32:1[M-H]$^-$) at $1/K_0$ = 1.281 Vs/cm$^2$, fragmented -40.0 eV. **c,** *m/z* 722.511 (PE P-36:4[M-H]$^-$) at $1/K_0$ = 1.317 Vs/cm$^2$, fragmented -43.4 eV. **d,** *m/z* 727.527 (PA 38:2[M-H]$^-$) at $1/K_0$ = 1.330 Vs/cm$^2$, fragmented -44.5 eV. **e,** *m/z* 821.517 (PI 33:1[M-H]$^-$) at $1/K_0$ = 1.416 Vs/cm$^2$ and fragmented with -53.5 eV. **f,** *m/z* 838.541 (PI 34:1[M-H]$^-$, $^{13}C_3$) at $1/K_0$ = 1.427 Vs/cm$^2$, fragmented -54.5 eV. **g,** *m/z* 859.530(PI 36:3[M-H]$^-$) at $1/K_0$ = 1.438 Vs/cm$^2$, fragmented -55.3 eV. **h,** *m/z* 915.593 (PI 40:4[M-H]$^-$, $^{13}C_2$) at $1/K_0$ = 1.494 Vs/cm$^2$, fragmented -59.1 eV. Detailed fragment ion identifications can be found in **Supplementary Tables 2-9**.

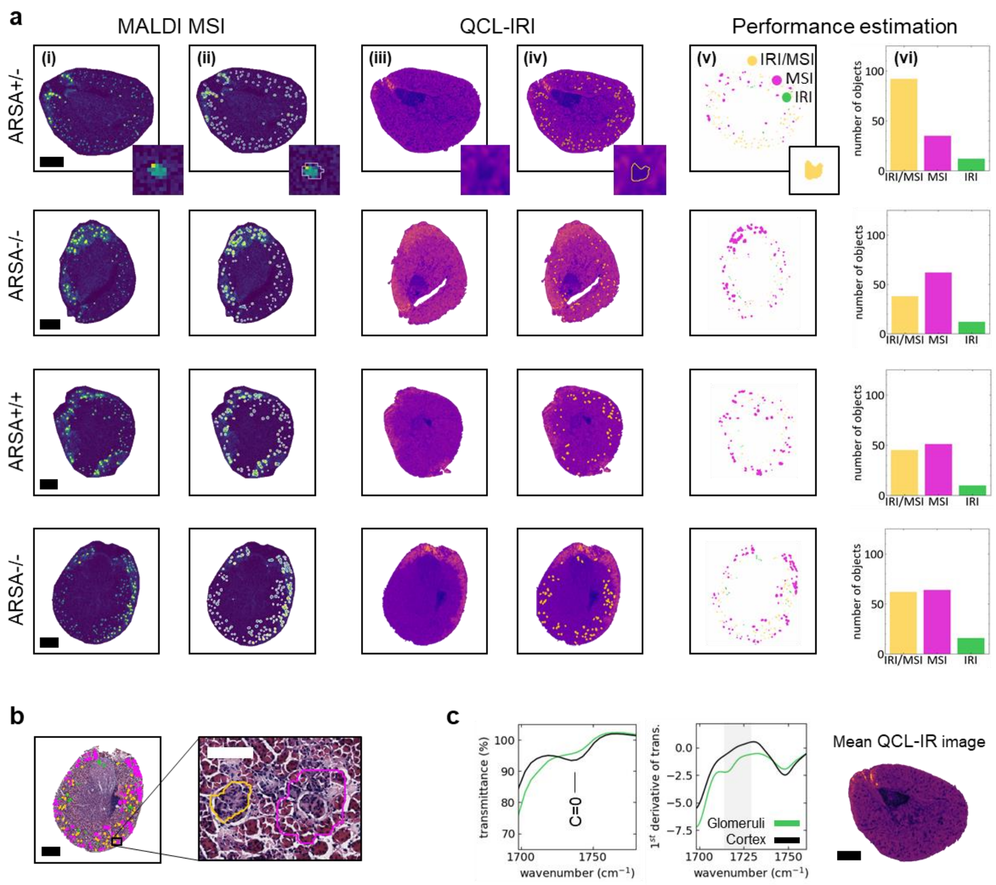

Supplementary Fig. 8. Focused segmentation of glomeruli-containing ROIs in kidney. **a**, Comparison of glomeruli-containing kidney regions in wild-type (ARSA+/+; 60 weeks), ARSA+/- (12 weeks) and ARSA-/- mice (12 and 60 weeks) by MALDI MSI (timsTOF fleX) and QCL-IRI microscopy (Hyperion II, 3.5x objective): **(i)**, ion image of *m/z* 1151.71 (GM3 34:1;O2[M-H]$^{-}$), ±10 ppm mass window; **(ii)**, Molecular probabilistic mapping (MPM) hotspot[3] for ganglioside GM3 34:1;O2 to aid probabilistic MSI segmentation of glomeruli; **(iii)**, QCL-infrared image at 1724 cm$^{-1}$ (selected from full spectrum of fingerprint region; 1$^{st}$ derivative of transmission, Hyperion II, 3.5x objective); **(iv)**, assignment of ROIs by QCL-IRI-based detection of glomeruli; **(v)**, Comparison of probabilistic MSI-MPM-based (magenta) and QCL-IRI-based (green; or both modalities: yellow) segmentation of glomeruli-containing kidney regions. Parameters for QCL-IRI-based identification were optimized to yield high ratios between the numbers of objects identified in both modalities vs. QCL-IRI alone (see method section). Note that identification of glomeruli-containing ROIs by QCL-IRI can be hampered by high fat content in tissue causing a dominant peak at 1740 cm$^{-1}$ (C=O vibrational band) (**Supplementary Fig. 5**) that limits spectral assignment. 40-80 glomeruli per tissue section were recognized by both modalities. Scale bars, 1 mm. **b**, Haematoxylin and eosin (H&E)-stained histological image of the ARSA-/- (60 weeks) section from **a**. Example regions identified as glomeruli-containing by MSI (magenta) and by both modalities (yellow) are superimposed. Scale bars, 1 mm. **c**, Mean transmittance (Hyperion II, 3.5x objective) and its 1$^{st}$ derivative at around 1720 cm$^{-1}$. The grey area highlights a distinct spectral region used to discriminate the glomerular from cortex region, as indicated by the mean QCL-IR image. Scale bar, 1 mm.

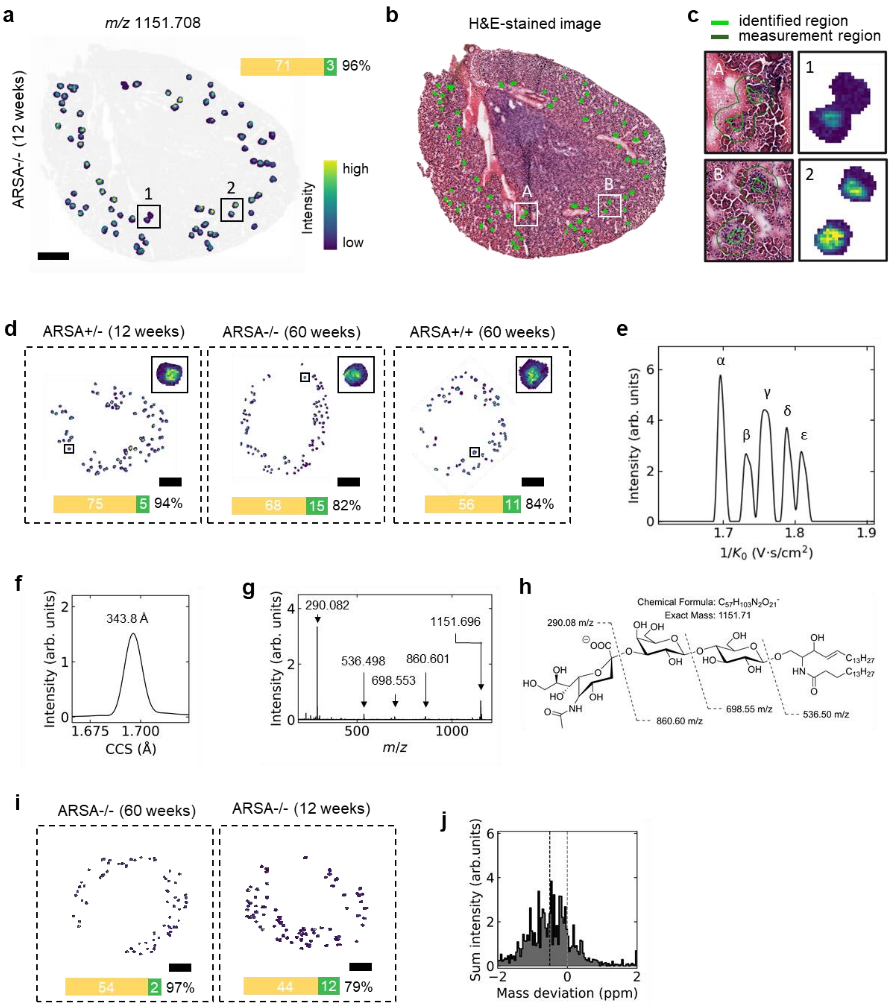

Supplementary Fig. 9. QCL-IRI-guided MSI of glomeruli-containing regions in kidney. **a,** QCL IRI-guided TIMS-MSI (timsTOF fleX) data for *m/z* 1151.708 (GM3 34:1;O2$[M-H]^-$), ±10 ppm mass window, and overlay on H&E-stained histological image (**b**). Number of QCL-IRI-segmented regions where characteristic glomeruli gangliosides[4] were identified by MSI (yellow part of the bar chart) and the number of regions below the threshold set for MSI signal intensity (green part), namely 20% of the maximum signal intensity of *m/z* 1151.708 (GM3 34:1;O2$[M-H]^-$). **c,** Magnified examples from **a** and **b**. **d,** Example data for tissue sections from three different mice (wild-type, heterozygotes and knock-out). **e,** Ion mobilogram for five gangliosides α: GM3 34:1;O2$[M-H]^-$, β: GM3 36:1;O2$[M-H]^-$, γ: GM3 38:1;O2$[M-H]^-$, δ: GM3 40:1;O2$[M-H]^-$, ε: GM3 42:1;O2$[M-H]^-$ identified by subsequent prm-PASEF $MS^2$analysis. **f**, Extracted ion mobilogram (343.8 $Å^2$) for GM3 34:1;O2$[M-H]^-$. **g**, prm-PASEF $MS^2$ spectrum of *m/z* 1151.696 (GM3 34:1;O2$[M-H]^-$). **h**, Chemical structure of GM3 34:1;O2$[M-H]^-$ including fragment assignment of. **i,** MRMS ion images for *m/z* 1151.7082 (GM3 34:1;O2$[M-H]^-$, 7T MRMS mass spectrometer), a ±3 ppm mass window. Glomerular structures where annotated as positive (yellow bar) if at least 10% of pixels contained the signal of *m/z* 1151.7082 (GM3 34:1;O2$[M-H^-]$), ±1 ppm mass window. **j,** Mass deviation of GM3 34:1;O2 as a function of the pixel index of the measurement regions in ARSA-/- (60 weeks) MRMS data. The red dotted line denotes the mean value determined from a Gaussian fit to the data.

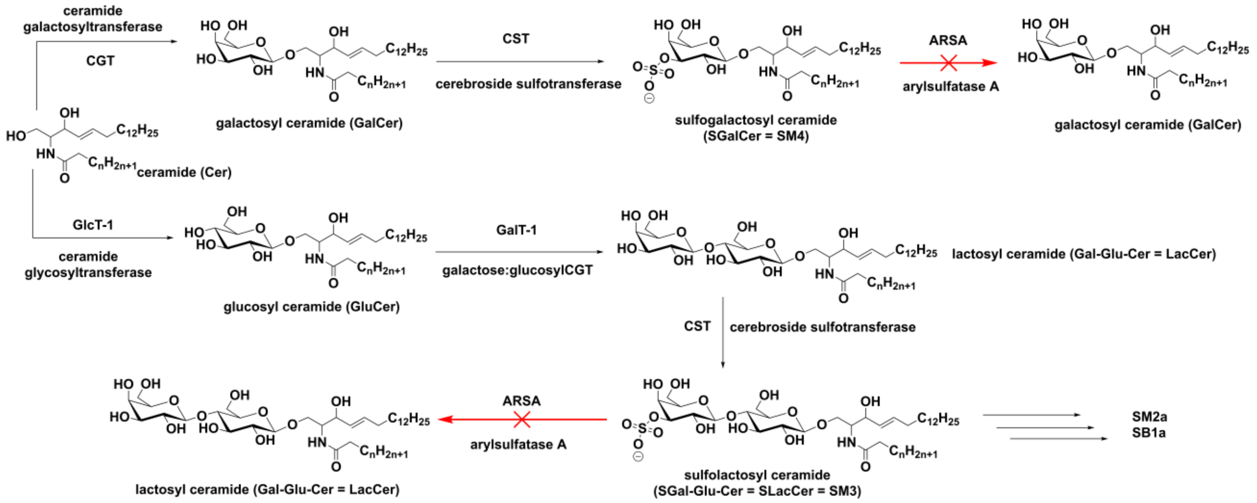

Supplementary Fig. 10. Biosynthesis and degradation of sulfo-glycosphingolipids in the ARSA-/- mouse model of human metachromatic leukodystrophy (MLD). Sulfatide biosynthesis utilizes ceramides for initial enzymatic β-glycosidic linkage of a hexose (galactose [Gal] or glucose [Glu]). Subsequent steps involve either the coupling of another hexose (Gal) to obtain lactosyl ceramides (LacCer) or of a sulfate group in the 3O-position of Gal to obtain sulfogalactosyl ceramides (SGalCer = **SM4**). For LacCer the sulfate group is coupled to the 3O-position of the terminal Gal, leading to sulfolactosyl ceramides (SLacCer = **SM3**). Complex sulfatides like SM2a or SB1a are generated from SM3. Due to the ARSA deficiency in the mouse model, hydrolytic removal of the sulfate group is blocked for SM4 and SM3, thus causing the accumulation of these lipids in multiple organs including kidney and brain.

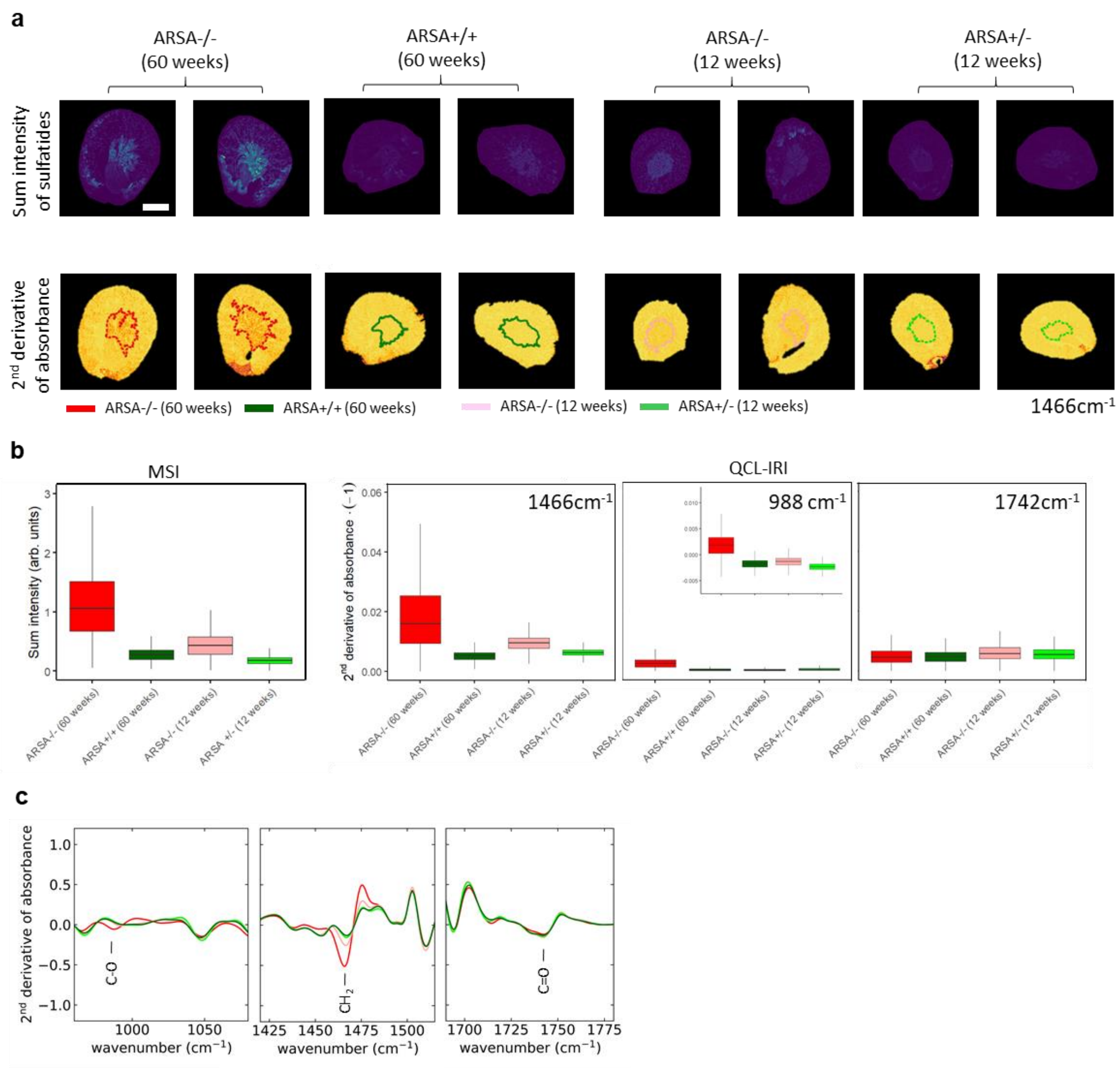

**Supplementary Fig. 11. (Semi-)quantitative analysis of sulfatide accumulation in the ISOM by MSI and QCL-IRI. a,** Sum intensity distribution of 87 sulfatides[5] (internal standard normalized) obtained by timsTOF-MSI in TIMS-off/qTOF mode for ARSA-/- (60 weeks) vs. ARSA+/+ (60 weeks) and ARSA-/- (12 weeks) vs. ARSA+/- (12 weeks) mice (top). QCL-IRI at the lipid associated band at 1466 $cm^{-1}$ ($CH_2$ vibration) (bottom). Contours of kidney inner segment of outer medulla (ISOM) ROI determined by clustering of MSI data (colored solid lines) are highlighted. **b**, Box-plots (n=2) of sum intensity of individual MSI and 2$^{nd}$ derivative of absorbance (linear to the concentration of molecular species[6]) of QCL-IRI of the ISOM region. QCL-IRI data is presented for the glycolipid-specific spectral band at 988 $cm^{-1}$, as well as the in general lipid associated but not sulfatide specific bands at 1466 $cm^{-1}$ and 1742 $cm^{-1}$. Both modalities show consistently (semi-)quantitative accumulation of sulfatides in the ISOM region. As expected, no difference between the different conditions of ARSA is observed for the C=O vibrations. **c,** Corresponding mean QCL-IRI spectra of the 2$^{nd}$ derivative of absorbance.

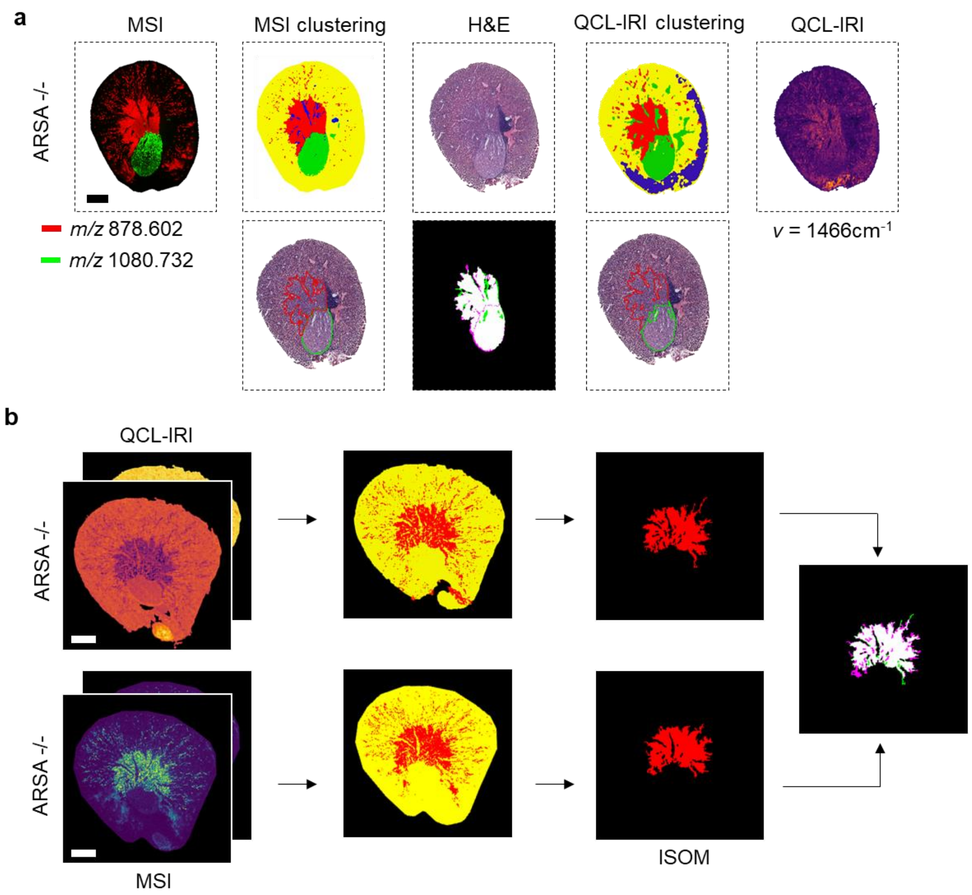

Supplementary Fig. 12. Comparable kidney segmentation with MSI and QCL-IRI. **a,** Top row, left: QCL-IRI-guided TIMS-MSI-derived ion images (timsTOF fleX, both represented within a ±10 ppm mass window) for *m/z* 878.602 (SM4 40:1;O3[M-H]$^-$) and 1080.732 (SM3 44:1;O2[M-H]$^-$) ; MSI image segmentation based on 60 selected *m/z* features obtained by bisecting k-means clustering (k=4); top row, middle: H&E-stained tissue section, QCL-IRI clustering based on 5 selected features with k=4; top row, right: IRI of single lipid-associated band at v=1466 cm$^{-1}$ (2$^{nd}$ derivative of absorbance, Hyperion II, 3.5x objective). The blue region in the IRI clustering may indicate tissue with a high fat content. Bottom row: Visualization of the regions of interest (ROIs) for the kidney's inner segment of the outer medulla (ISOM) and for the kidney inner medulla/papilla (IMP), as defined by MSI or IRI, superimposed on the H&E-stained image. A comparison of the respective IMP and ISOM regions defined by MSI and QCL-IRI yielded a Dice-Sorensen coefficient[7] of 93%. Scale bar, 1 mm. **b,** Sulfatide distributions in ARSA -/- mouse kidney. Top row, left: High-resolution QCL-IRI images (2$^{nd}$ derivative data, Hyperion II, 15x objective) for the lipid-associated bands at 990 cm$^{-1}$ (C-O vibration of the 3-sulfogalactosyl head group) and 1466 cm$^{-1}$ ($CH_2$ bending vibration)[8]; middle: corresponding clusters for k-means clustering with k=2; top row, right: selected ISOM segment/ROI. Bottom row, left: Ion images (timsTOF fleX, qTOF mode) acquired with 20 µm lateral step-size and clustering for the two abundant *m/z* values (878.606 [M-H]$^-$) and 880.612 [M-H]$^-$; ±10 ppm mass window). Comparison of the respective ISOM regions based on MSI and QCL-IRI yields a Dice-Sorensen coefficient of 87%. Scale bar, 1 mm.

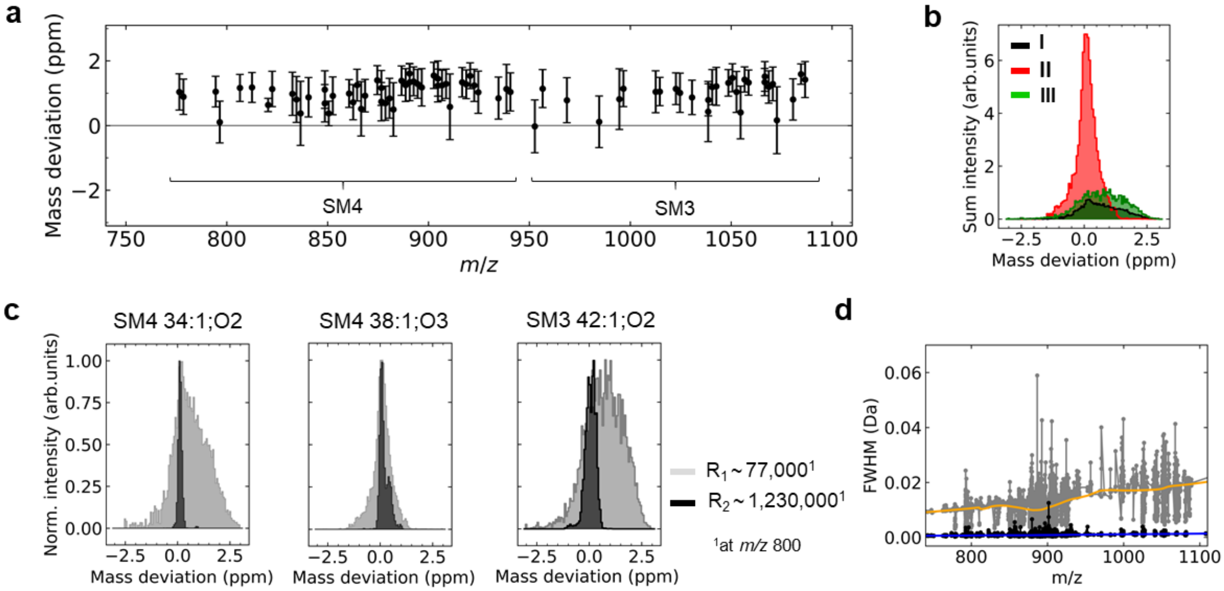

Supplementary Figure 13. Comparison of QCL-guided MR-MSI (16s FID) and conventional MRMS MSI (1s FID). **a**, Mean mass deviation for sulfatides identified by MR-MSI (solariX 7T XR) with a mass resolution of $R_1$~77,000 at *m/z* 800. Means and standard deviations for each sulfatide are obtained from the sum intensity-histograms in b, which results from the mean *m/z* value (determined across all pixels) of a given sulfatide. **b**, Histogram of the sum intensity for the three sulfatides **I** SM4 34:1;O2[M-H]$^-$, **II** SM4 38:1;O3[M-H]$^-$, and **III** SM3 42:1;O2[M-H]$^-$. **c**, Histogram of the normalized sum intensities from b and **Fig. 2c** for two different cases of $R_1$~77,000 and $R_2$~1,230,000. Direct comparison shows a 3- to 6-times more precise determination of the *m/z* value of the corresponding sulfatide. **d**, Full width at half maximum (FWHM) as a function of *m/z* value for two different free induction decay (FID) times of 1s and 16s. The orange and blue curves result from a locally estimated scatterplot smoothing (loess)[3] and are plotted to guide-the-eye. On average the ratio of FWHM with $R_2$ to FWHM with $R_1$ agrees well with the expected mass resolution determined by the relative duration of the FID times.

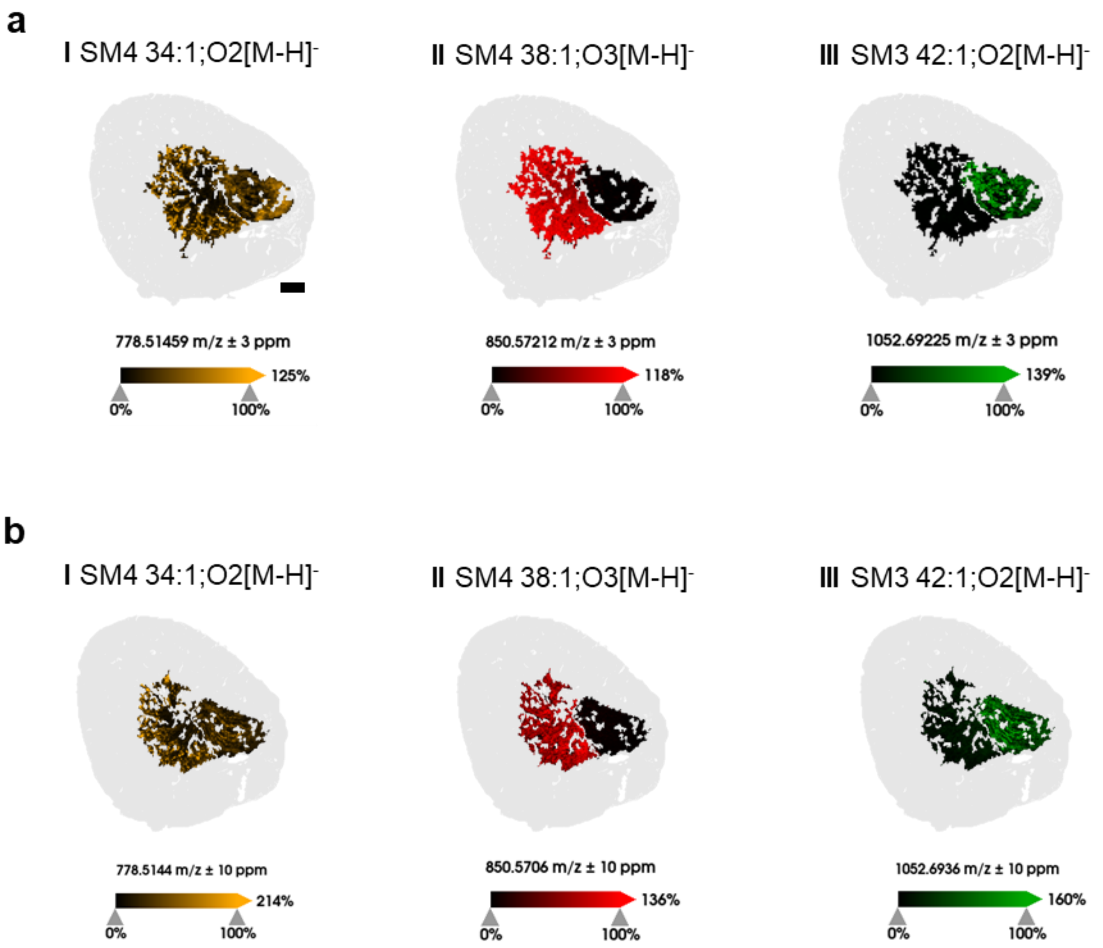

**Supplementary Figure 14. QCL-IRI-guided MR-MSI- (a) and tims-on-MSI (b) derived ion images of kidney ISOM and IMP.** Images are given for three molecules and conditions: i) *m/z* 778.5145 (SM4 34:1;O2[M-H]$^-$), showing a similar ion intensity in both regions, ii) *m/z* 850.5720 (SM4 38:1;O3[M-H]$^-$), showing much stronger ion intensity in ISOM, and iii) *m/z* 1052.6925 (SM3 42:1;O2[M-H]$^-$) showing much stronger ion intensity in IMP. All ion images in **(a)** are presented within a mass window of ±3 ppm (±10 ppm in **(b)**). Scale bar, 500 µm.

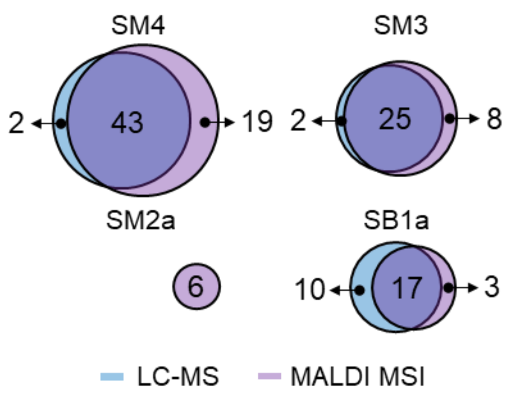

Supplementary Figure 15. Venn diagram of sulfatide subclasses identified by LC-TIMS-MS (blue) and MALDI-TIMS-MSI (purple). Sulfatide identifications are compared for the subclasses SM4, SM3, SM2a and SB1a. Note that SB1a isoforms are measured as SB1a[M-2H]$^{2-}$ in LC-MS and as SB1a[M-HSO3]$^{-}$ in MSI.

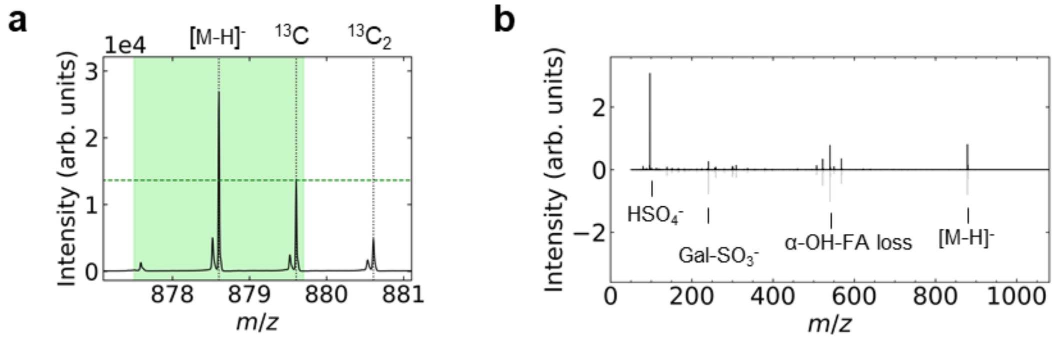

Supplementary Figure 16. Representative MS1 and MS2 spectra for the sulfatide SM40:1;O3[M-H]$^{-}$. **a,** Average MS spectrum obtained by QCL-IRI guided MSI (timsTOF fleX) of an ARSA-/- mouse IMP region. The vertical, black dotted lines indicate the theoretical *m/z* values for the mono-isotopic mass ([M-H]$^{-}$), the first carbon isotope ($^{13}C$) and the second carbon isotope ($^{13}C_2$). The green area indicates a typical isolation window of ±1.1 Da. The green horizontal dotted line represents the ion intensity at the position of the theoretical *m/z* value of the first carbon isotope. Within the isolation window, there is no additional *m/z* peak unrelated to SM4 40:1;O3[M-H]$^{-}$ with a signal intensity higher than the peak of the first carbon isotope. Hence, the spectrum is considered as "clean". **b,** Butterfly plot of two MS$^2$ spectra of SM40:1;O3[M-H]$^{-}$. The black spectrum (top half) was obtained by prm-PASEF, the grey spectrum (lower half) "conventionally" by MS$^2$ without prior ion mobility separation ("on-tissue"). Main fragments were assigned as $HSO_4^-$ (*m/z* 96.96), Gal-$SO_3^-$ (*m/z* 241.00), and α-hydroxy fatty acid (α-OH-FA) loss-related.

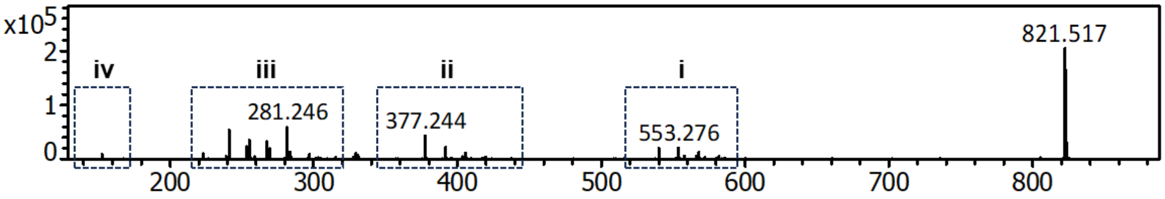

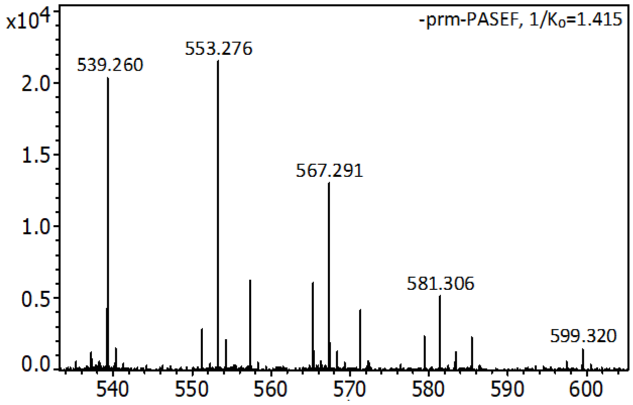

| *m/z* measured | Fragment ion | Sum formula |
|---|---|---|
| **821.517** | [M-H]$^-$ | $C_{42}H_{78}O_{13}P^-$ |
| **599.320** | [M-15:1ketene-H]$^-$ | $C_{27}H_{52}O_{12}P^-$ |
| **597.303** | [M-15:0ketene-H]$^-$ | $C_{27}H_{50}O_{12}P^-$ |
| **585.306** | [M-16:1ketene-H]$^-$ | $C_{26}H_{50}O_{12}P^-$ |
| **583.286** | [M-16:0ketene-H]$^-$ | $C_{26}H_{48}O_{12}P^-$ |
| **581.306** | [M-FA15:1-H]$^-$ | $C_{27}H_{50}O_{11}P^-$ |
| **579.292** | [M-FA15:0-H]$^-$ | $C_{27}H_{48}O_{11}P^-$ |
| **571.285** | [M-17:1ketene-H]$^-$ | $C_{25}H_{48}O_{12}P^-$ |
| **567.291** | [M-FA16:1-H]$^-$ | $C_{26}H_{48}O_{11}P^-$ |
| **565.280** | [M-FA16:0-H]$^-$ | $C_{26}H_{46}O_{11}P^-$ |
| **557.309** | [M-18:1ketene-H]$^-$ | $C_{24}H_{46}O_{12}P^-$ |
| **553.276** | [M-FA17:1-H]$^-$ | $C_{25}H_{46}O_{11}P^-$ |
| **551.261** | [M-FA17:0-H]$^-$ | $C_{25}H_{44}O_{11}P^-$ |
| **539.260** | [M-FA18:1-H]$^-$ | $C_{24}H_{44}O_{11}P^-$ |
| **537.243** | [M-FA18:0-H]$^-$ | $C_{24}H_{42}O_{11}P^-$ |

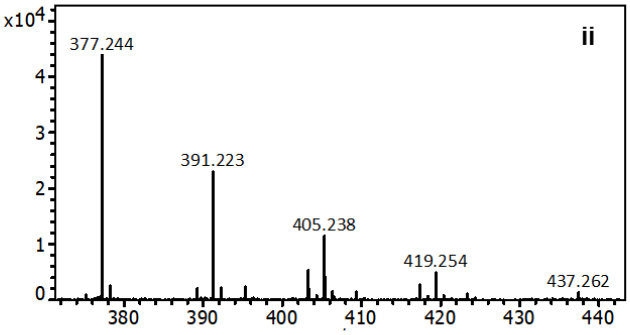

| *m/z* measured | Fragment ion | Sum formula |
|---|---|---|
| **437.262** | [M-15:1ketene-inositol-H]$^-$ | $C_{21}H_{42}O_7P^-$ |
| **423.248** | [M-16:1ketene-inositol-H]$^-$ | $C_{20}H_{40}O_7P^-$ |
| **419.254** | [M-15:1FA-inositol-H]$^-$ | $C_{21}H_{40}O_6P^-$ |
| **417.237** | [M-15:0FA-inositol-H]$^-$ | $C_{21}H_{38}O_6P^-$ |
| **409.234** | [M-17:1ketene-inositol-H]$^-$ | $C_{19}H_{38}O_7P^-$ |
| **405.238** | [M-16:1FA-inositol-H]$^-$ | $C_{20}H_{38}O_6P^-$ |
| **403.225** | [M-16:0FA-inositol-H]$^-$ | $C_{20}H_{36}O_6P^-$ |
| **395.253** | [M-18:1ketene-inositol-H]$^-$ | $C_{18}H_{36}O_7P^-$ |
| **391.223** | [M-17:1FA-inositol-H]$^-$ | $C_{19}H_{36}O_6P^-$ |
| **389.208** | [M-17:0FA-inositol-H]$^-$ | $C_{19}H_{34}O_6P^-$ |
| **377.244** | [M-18:1FA-inositol-H]$^-$ | $C_{18}H_{34}O_6P^-$ |
| **375.230** | [M-18:0FA-inositol-H]$^-$ | $C_{18}H_{32}O_6P^-$ |

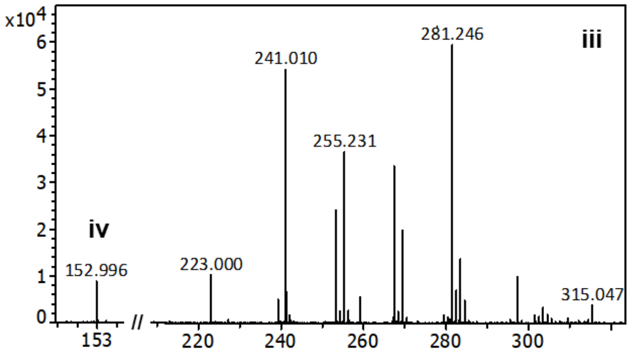

| *m/z* measured | Fragment ion | Sum formula |
|---|---|---|
| **315.047** | [M-FA-ketene-H]$^-$ | $C_9H_{16}O_{10}P^-$ |
| **297.036** | [M-FA-FA-H]$^-$ | $C_9H_{14}O_9P^-$ |
| **283.261** | [FA18:0-H]$^-$ | $C_{18}H_{35}O_2^-$ |
| **281.246** | [FA18:1-H]$^-$ | $C_{18}H_{33}O_2^-$ |
| **269.246** | [FA17:0-H]$^-$ | $C_{17}H_{33}O_2^-$ |
| **267.230** | [FA17:1-H]$^-$ | $C_{17}H_{31}O_2^-$ |
| **259.090** | [inositolPO$_3$H$_2$-H]$^-$ | $C_6H_{12}O_9P^-$ |
| **255.231** | [FA16:0-H]$^-$ | $C_{16}H_{31}O_2^-$ |
| **253.215** | [FA16:1-H]$^-$ | $C_{16}H_{29}O_2^-$ |
| **241.216** | [FA15:0-H]$^-$ | $C_{15}H_{29}O_2^-$ |
| **241.010** | [inositolPO$_3$H$_2$-H$_2$O-H]$^-$ | $C_6H_{10}O_8P^-$ |
| **239.199** | [FA15:1-H]$^-$ | $C_{15}H_{27}O_2^-$ |
| **223.000** | [inositolPO$_3$H-2H$_2$O-H]$^-$ | $C_6H_8O_7P^-$ |
| **152.996** | [glycerolPO$_3$H$_2$-H]$^-$ | $C_3H_6O_5P^-$ |

Supplementary Figure 17. Unequivocal identification of odd-chain PI 33:1[M-H]$^-$ (m/z 821.517) by MALDI-TIMS-MSI with prm-PASEF. Isolated at 1/$K_0$ = 1.416 Vs/cm$^2$ and fragmented with -53.5 eV. Inlet **i** represents a detailed view for *m/z* 530–610, **ii** for *m/z* 370–445, **iii** for *m/z* 210–310, and **iv** for *m/z* 151–154. The detailed descriptions of the elucidated fragment ions are displayed in tables together with chemical sum formulae. Steric effects made the neutral loss (NL) of the first FA more favorable at the *sn*-2 position[9]. Highest intensities were observed for the NL of FA18:1 and FA17:1. As a consequence, PI 15:0/18:1 and PI 16:0/17:1 were identified as the predominant isomeric structures[9].

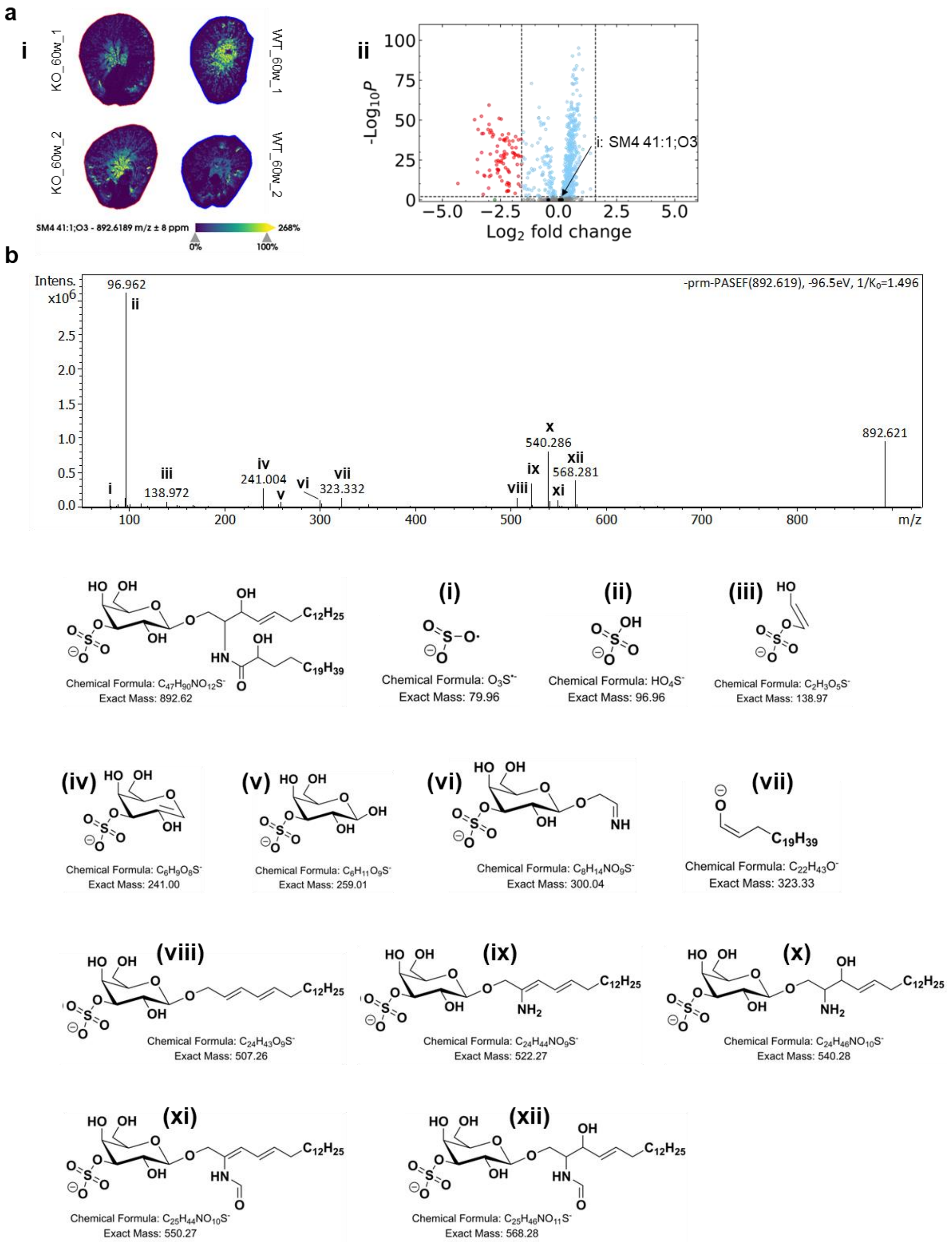

Supplementary Figure 18. Unequivocal identification of odd-chain sulfatides by MALDI-TIMS-MSI with prm-PASEF. **a, (i)** MALDI-MSI-derived ion images (timsTOF flex, qTOF mode) of *m/z* 892.619 (SM4 41:1;O3 $[M-H]^-$) for two data sets of ARSA-/- (KO, left) and ARSA+/+ (WT, right). **(ii)** MSI-derived volcano scatter plot suggests the non-significance for the accumulation of odd-chain sulfatides (black dots), in contrast to significant features (red dots). **b,** prm-PASEF-derived $MS^2$ spectrum of *m/z* 892.621 (SM4 41:1;O3$[M-H]^-$), isolated at $1/K_0$ = 1.496 Vs/cm² and fragmented at -96.5 eV. The structures of the identified fragments are shown below.

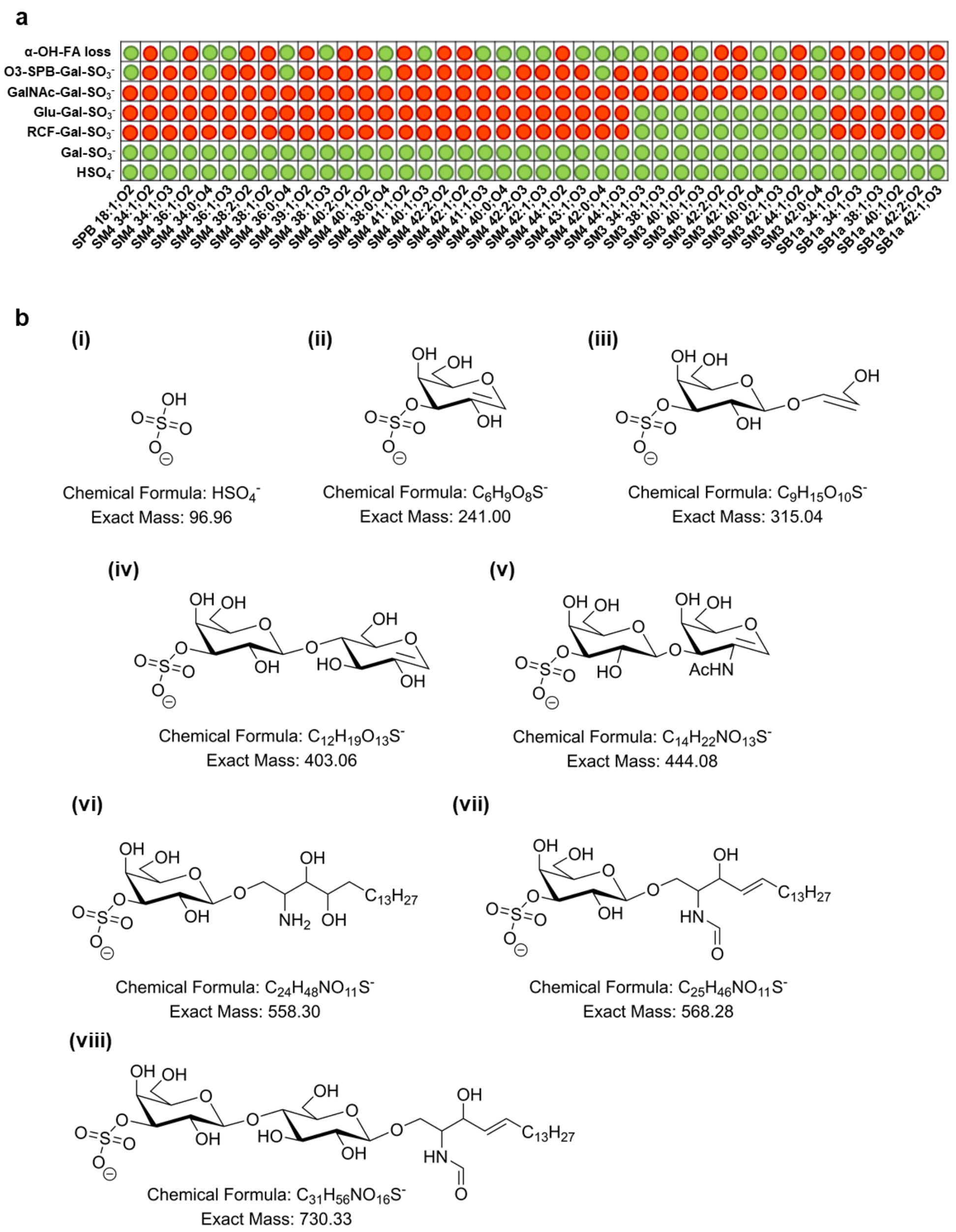

#### Supplementary Figure 19. Structural representation of characteristic sulfatide fragments.

**a,** Sulfatide fragments observed in TIMS-MSI and prm-PASEF analysis. **b,** Chemical structures of these fragments: **(i)**, cleavage of the sulfate moiety leads to fragment $HSO_4^-$ at *m/z* 96.96. **(ii)**, cleavage of the Gal-$SO_3$ moiety yields fragment at *m/z* 241.00. **(iii)**, ring-cross-fragmentation (RCF) within the glucose part of SM3 leads to fragment at *m/z* 315.04. **(iv)**, cleavage of the entire Glu-Gal-$SO_3$ moiety forms the fragment at *m/z* 403.06. **(v)**, for SB1a, cleavage of GalNAc-Gal-$SO_3$ leads to the fragment *m/z* 444.08. **(vi)**, neutral loss of the entire α-OH-FA moiety results in fragment *m/z* 558.30 for sulfatides consisting of a tri-hydroxylated sphingoid base (phytosphingosine, O3-SPB). **(vii)** and **(viii)**, neutral loss of the α-OH-FA leads to fragments at *m/z* 568.28 and *m/z* 730.33 for SM3 and SM4, respectively.

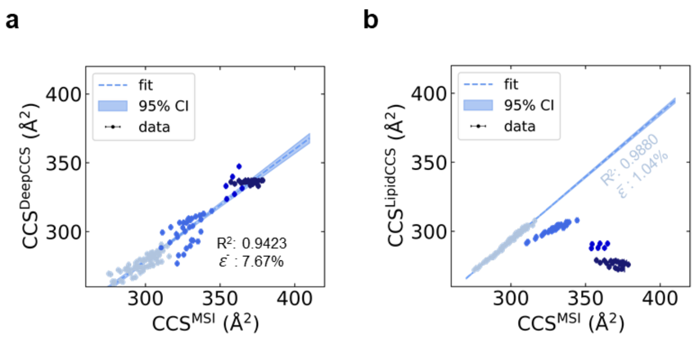

Supplementary Figure 20. Correlation of experimentally deduced CCS values (QCL IRI-guided MALDI-TIMS-MSI data) and CCS values predicted by IT tools: a, DeepCCS and b, LipidCCS. For all sulfatide CCS identified by MALDI-TIMS-MSI (timsTOF fleX) in this study, AllCCS2 yielded the most accurate prediction, i.e. mean relative error $\bar{\varepsilon}$ of about 1% (Fig. 4c). For individual sulfatide classes it can be increased, e.g. $\bar{\varepsilon}$ = 2.48% for SM2a. The accuracy of the predicted values for LipidCCS for the SM4 class is also on the level of 1%, but inaccurate for all other sulfatide classes. In contrast, $\bar{\varepsilon}$ is about 8% for CCS values predicted by DeepCCS.

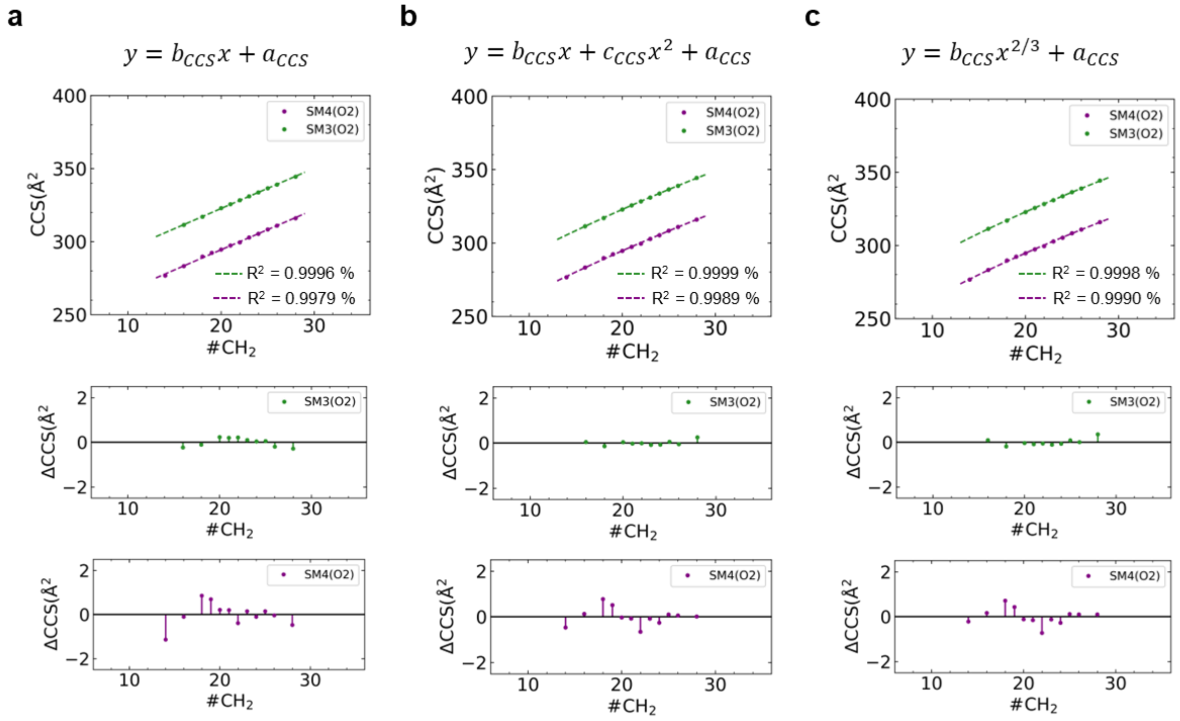

Supplementary Figure 21. Modelling of experimental CCS values as a function of the chain length of the *N*-acyl-linked fatty acid (FA). Linear data fit (**a**), 2nd order polynomial fit **(b)**, and $y = b_{CCS}x^{2/3} + a_{CCS}$ (**c**), a fit commonly used to describe CCS values of polymers[10]. A global least square fitting procedure was used with fixed amplitudes, $b_{CCS}$ and $c_{CCS}$, each. Residuals are shown, indicating best agreement between the data and the 2nd order polynomial fit **b**.

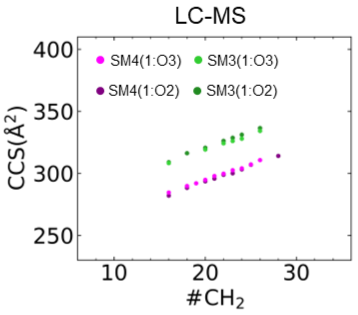

Supplementary Figure 22. Experimental (MALDI-TIMS-MSI and LC-MS) and predicted CCS values as a function of *N*-acyl-linked fatty acid chain length: Experimental LC-TIMS-MS**.** CCS values for the subclasses SM4 18+n:1;O3 (magenta), SM4 18+n:1;O2 (purple), SM3 18+n:1;O3 (light green), SM3 18+n:1;O2 (dark green), SM2a 18+n:1;O3 (orange), and SM2a 18+n:1;O2 (red). n denotes the chain length alteration.

**a**

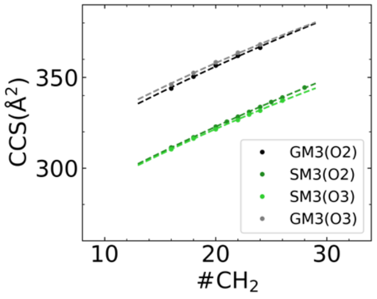

**b**

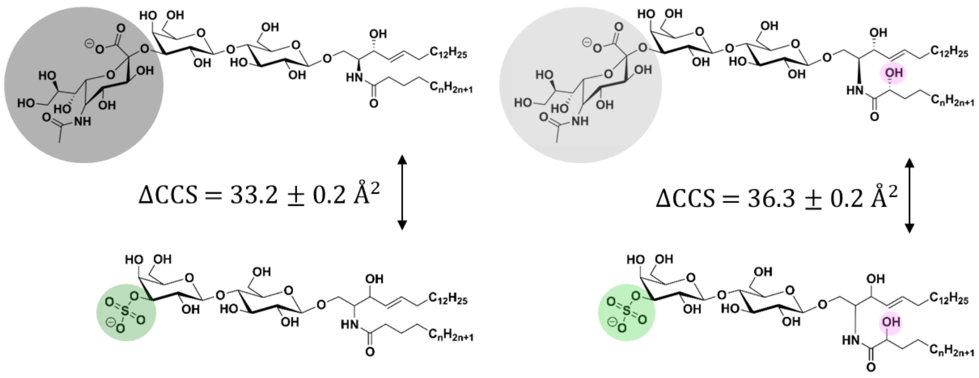

Supplementary Figure 23. Comparison of experimental MALDI-TIMS-MSI CCS values for SM3 and GM3 subclasses. **a,** Comparison of CCS values for ganglioside series GM3(O2) (black) and GM3(O3) (gray) against sulfatide series SM3(O2) (dark green) and SM3(O3) (light green) as a function of *N*-acyl-linked fatty acid chain length. Data is modelled by a 2nd order polynomial fit with fixed amplitudes of the linear and quadratic term of the polynomial fit. **b**, differences in CCS values associated with the contribution of the sulfate group (green) or the neuraminic acid group (gray) for SM3(O2) vs. GM3(O2) (left), and SM3(O3) vs. GM3(O3) (right). The difference is increased for SM3(O3) vs. GM3(O3), suggesting a possible influence of the α-hydroxyl group (magenta, right).

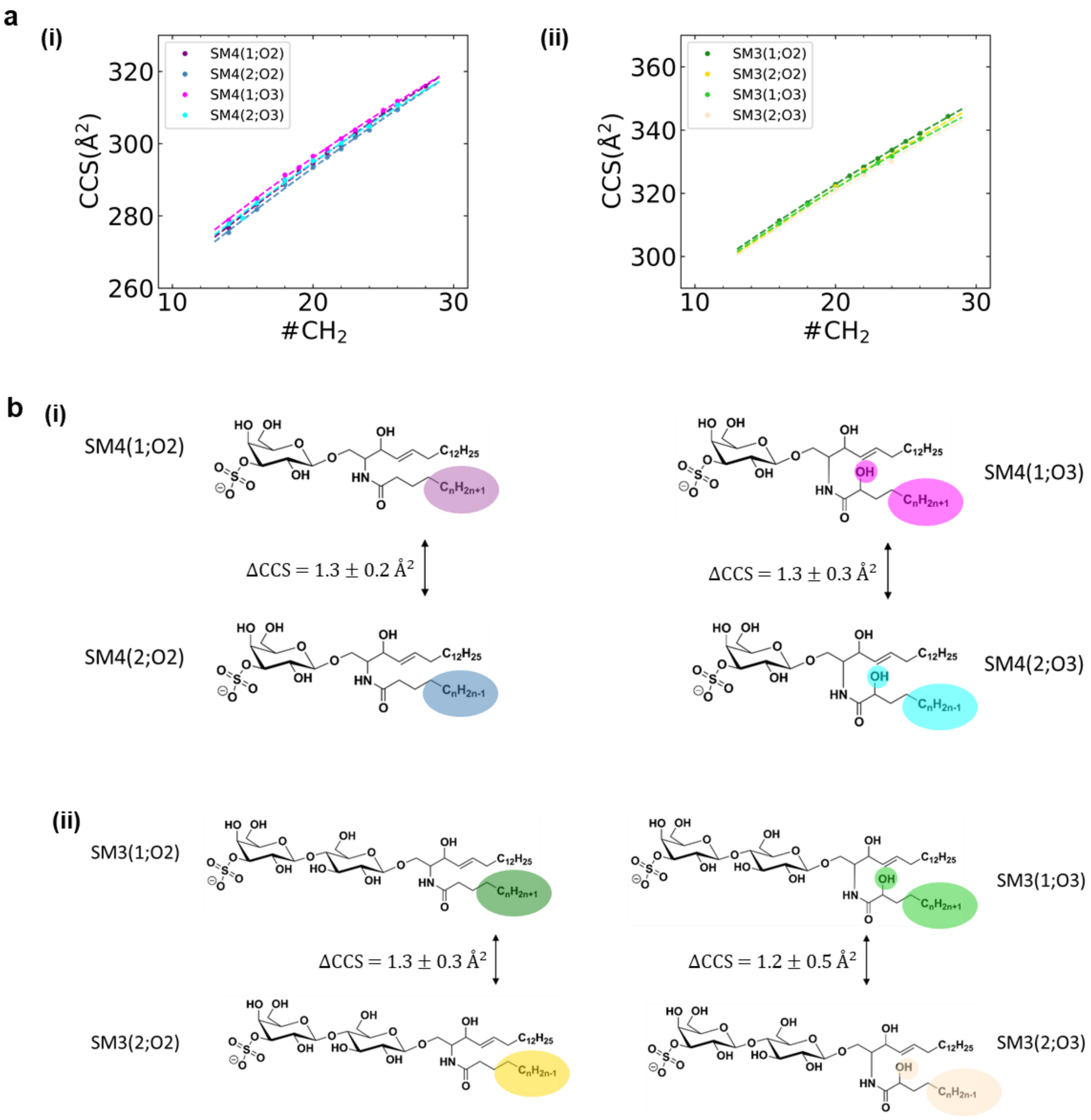

Supplementary Figure 24. Evolution of the relative CCS values between selected SM3 and SM4 subclasses incorporating either a saturated FA ($C_nH_{2n+1}$), or a mono-unsaturated FA ($C_nH_{2n-1}$). **a,** Comparison of CCS values for sulfatide series SM4(O2/O3) **(i)**, and SM3(O2/O3) **(ii)**, depending on the degree of unsaturation in their *N*-acyl-linked FAs. Data is modelled by a 2nd order polynomial fit with fixed amplitudes of the linear and quadratic term of the polynomial fit. **b**, The differences in CCS values associated to the contribution of the degree of unsaturation in the *N*-acyl-linked FA for SM4(O2/O3) **(i)**, and SM3(O2/O3) **(ii)** are shown. The mean relative difference was 1.3±0.2 Å$^2$.

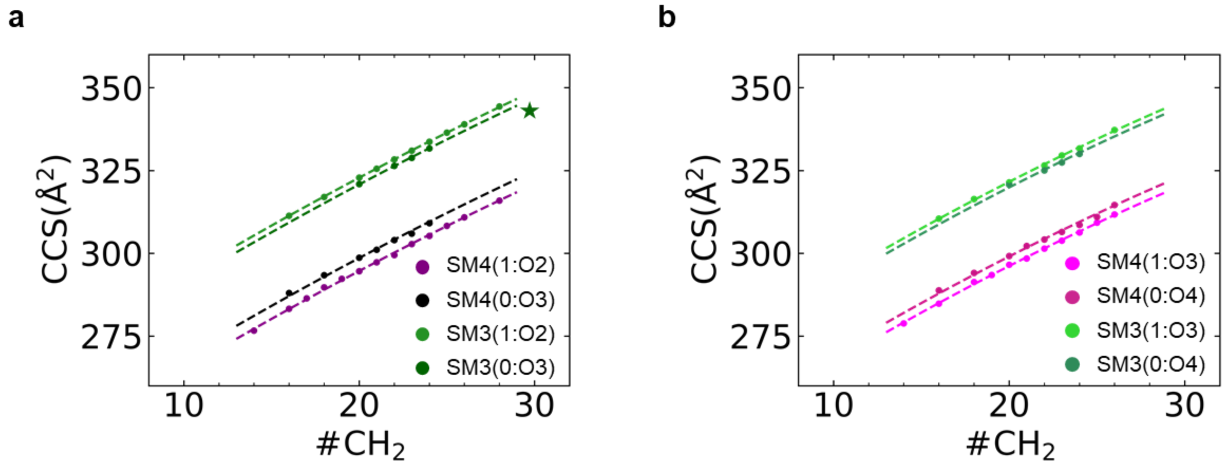

Supplementary Figure 25. Evolution of the relative CCS values between selected SM3 and SM4 subclasses. **a,** Comparison of CCS values for sulfatide series SM4/SM3(1:O3) against SM4/SM3(1:O2). **b**, Comparison of CCS values for sulfatide series SM4/SM3(0:O4) against SM4/SM3(1:O3). For SM3(0:O3), the TIMS-MSI data was taken, and a corrected offset was considered. Data is modelled by a $2^{nd}$ order polynomial fit with fixed amplitudes of the linear and quadratic term of the polynomial fit.

### Supplementary Tables

Supplementary Table 1. Overview of significant features (as determined by lasso method) in MCF and BCF. Uncertainties as standard deviations (*n*=18) in parentheses. All ions detected as $[M-H]^-$.

| Name | Chemical sum formula | *m/z* theoret. | 1/K0 [V·s/cm²] | CCS [Å²] | *m/z* timsTOF | ppm timsTOF |
|---|---|---|---|---|---|---|
| FA 20:5 | C20H30O2 | 301.217304 | 0.863(6) | 181(1) | 301.219(4) | 7.00 |
| lyso-PI 18:0 | C27H53O12P | 599.320187 | 1.176(7) | 241(1) | 599.320(1) | -0.66 |
| PE 32:1 | C37H72NO8P | 688.492278 | 1.289(7) | 263(1) | 688.494(1) | 2.73 |
| PE (P-36:4) | C41H74NO7P | 722.513014 | 1.330(7) | 271(1) | 722.512(1) | -0.94 |
| PA 38:2 | C41H77O8P | 727.528330 | 1.336(7) | 273(1) | 727.529(1) | 0.58 |
| PI 33:1 | C42H79O13P | 821.518553 | 1.427(7) | 291(1) | 821.516(2) | 2.74 |
| PI 34:1 | C43H81O13P | 835.534203 | 1.430(7) | 291(1) | 835.534(1) | -0.74 |
| PI 36:3 | C45H81O13P | 859.534203 | 1.444(7) | 294(1) | 859.533(1) | -1.16 |
| PI 40:4 | C49H87O13P | 913.581153 | 1.497(7) | 304(1) | 913.581(1) | -0.50 |

**Supplementary Table 2. Identified fragment ions of lyso-PI 18:0[M-H]$^-$ (*m/z* 599.317).** Isolated at $1/K_0$ = 1.171 Vs/cm² and fragmented with -40.0 eV. Fragment patterns were modelled according to Hsu and Turk, 2009[9].

| *m/z* measured | Fragment ion | Sum formula |
|---|---|---|
| 599.317 | [M-H]$^-$ | $C_{27}H_{52}O_{12}P^-$ |
| 419.254 | [M-inositol-H]$^-$ | $C_{21}H_{40}O_6P^-$ |
| 315.047 | [M-FA18:0-H]$^-$ | $C_9H_{16}O_{10}P^-$ |
| 283.262 | [FA18:0-H]$^-$ | $C_{18}H_{35}O_2^-$ |
| 241.010 | [inositolPO$_3$H$_2$-H]$^-$ | $C_6H_{10}O_8P^-$ |
| 152.994 | [glycerolPO$_3$H$_2$-H]$^-$ | $C_3H_6O_5P^-$ |

**Supplementary Table 3. Identified fragment ions of PE 32:1[M-H]$^-$ (*m/z* 688.489).** Isolated at $1/K_0$ = 1.281 Vs/cm² and fragmented with -40.0 eV. PE 16:0/16:1 was identified as the predominant isomeric structure[9].

| *m/z* measured | Fragment ion | Sum formula |
|---|---|---|
| 688.489 | [M-H]$^-$ | $C_{37}H_{71}NO_8P^-$ |
| 478.292 | [M-18:1ketene-H]$^-$ | $C_{23}H_{45}NO_7P^-$ |
| 460.266 | [M-FA18:1-H]$^-$ | $C_{23}H_{43}NO_6P^-$ |
| 452.276 | [M-16:1ketene-H]$^-$ | $C_{21}H_{43}NO_7P^-$ |
| 450.261 | [M-16:0ketene-H]$^-$ | $C_{21}H_{41}NO_7P^-$ |
| 434.266 | [M-FA16:1-H]$^-$ | $C_{21}H_{41}NO_6P^-$ |
| 432.250 | [M-FA16:0-H]$^-$ | $C_{21}H_{39}NO_6P^-$ |
| 424.245 | [M-14:0ketene-H]$^-$ | $C_{19}H_{39}NO_7P^-$ |
| 281.246 | [FA18:1-H]$^-$ | $C_{18}H_{33}O_2^-$ |
| 255.231 | [FA16:0-H]$^-$ | $C_{16}H_{31}O_2^-$ |
| 253.215 | [FA16:1-H]$^-$ | $C_{16}H_{29}O_2^-$ |
| 227.199 | [FA14:0-H]$^-$ | $C_{14}H_{27}O_2^-$ |

**Supplementary Table 4. Identified fragment ions of PE P-36:4[M-H]$^-$ (*m/z* 722.511).** Isolated at $1/K_0$ = 1.317 Vs/cm² and fragmented with -43.4 eV. PE P-18:0/18:4 was identified as the predominant isomeric structure[9].

| *m/z* measured | Fragment ion | Sum formula |
|---|---|---|
| 722.511 | [M-H]$^-$ | $C_{41}H_{73}NO_7P^-$ |
| 437.264 | [M-20:4ketene-H]$^-$ | $C_{21}H_{42}NO_5P^-$ |
| 419.254 | [M-FA20:4-H]$^-$ | $C_{21}H_{39}NO_4P^-$ |
| 303.230 | [FA20:4-H]$^-$ | $C_{20}H_{31}O_2^-$ |
| 301.214 | [FA20:5-H]$^-$ | $C_{20}H_{29}O_2^-$ |
| 283.263 | [FA18:0-H]$^-$ | $C_{18}H_{35}O_2^-$ |
| 281.247 | [FA18:1-H]$^-$ | $C_{18}H_{33}O_2^-$ |
| 255.232 | [FA16:0-H]$^-$ | $C_{16}H_{31}O_2^-$ |
| 152.994 | [MePO$_3$EtNH$_2$-H]- | $C_3H_8NO_4P^-$ |

**Supplementary Table 5. Identified fragment ions of PA 38:2[M-H]⁻ (*m/z* 727.523).** Isolated at $1/K_0$ = 1.330 Vs/cm² and fragmented with -44.5 eV. PA 20:1/18:1 was identified as the predominant isomeric structure[9].

| *m/z* measured | Fragment ion | Sum formula |
|---|---|---|
| 727.523 | $[M-H]^-$ | $C_{41}H_{76}O_8P^-$ |
| 463.281 | $[M-18:1ketene-H]^-$ | $C_{23}H_{44}NO_7P^-$ |
| 445.269 | $[M-FA18:1-H]^-$ | $C_{23}H_{42}NO_6P^-$ |
| 419.254 | $[M-FA20:2-H]^-$ | $C_{21}H_{40}NO_6P^-$ |
| 337.307 | $[FA22:1-H]^-$ | $C_{22}H_{41}O_2^-$ |
| 309.278 | $[FA20:1-H]^-$ | $C_{20}H_{37}O_2^-$ |
| 307.258 | $[FA20:2-H]^-$ | $C_{20}H_{35}O_2^-$ |
| 281.246 | $[FA18:1-H]^-$ | $C_{18}H_{33}O_2^-$ |
| 253.216 | $[FA16:1-H]^-$ | $C_{16}H_{29}O_2^-$ |
| 152.994 | $[glycerolPO_3H_2-H]^-$ | $C_3H_6O_5P^-$ |

**Supplementary Table 6. Identified fragment ions of PI 33:1[M-H]⁻ (*m/z* 821.517).** Isolated at $1/K_0$ = 1.416 Vs/cm² and fragmented with -53.5 eV. PI 15:0/18:1 and PI 16:0/17:1 were identified as the predominant isomeric structures[9].

| *m/z* measured | Fragment ion | Sum formula |
|---|---|---|
| 821.517 | $[M-H]^-$ | $C_{42}H_{78}O_{13}P^-$ |
| 599.320 | $[M-15:1ketene-H]^-$ | $C_{27}H_{52}O_{12}P^-$ |
| 597.303 | $[M-15:0ketene-H]^-$ | $C_{27}H_{50}O_{12}P^-$ |
| 585.306 | $[M-16:1ketene-H]^-$ | $C_{26}H_{50}O_{12}P^-$ |
| 583.286 | $[M-16:0ketene-H]^-$ | $C_{26}H_{48}O_{12}P^-$ |
| 581.306 | $[M-FA15:1-H]^-$ | $C_{27}H_{50}O_{11}P^-$ |
| 579.292 | $[M-FA15:0-H]^-$ | $C_{27}H_{48}O_{11}P^-$ |
| 571.285 | $[M-17:1ketene-H]^-$ | $C_{25}H_{48}O_{12}P^-$ |
| 567.290 | $[M-FA16:1-H]^-$ | $C_{26}H_{48}O_{11}P^-$ |
| 565.280 | $[M-FA16:0-H]^-$ | $C_{26}H_{46}O_{11}P^-$ |
| 557.309 | $[M-18:1ketene-H]^-$ | $C_{24}H_{46}O_{12}P^-$ |
| 553.277 | $[M-FA17:1-H]^-$ | $C_{25}H_{46}O_{11}P^-$ |
| 551.260 | $[M-FA17:0-H]^-$ | $C_{25}H_{44}O_{11}P^-$ |
| 539.261 | $[M-FA18:1-H]^-$ | $C_{24}H_{44}O_{11}P^-$ |
| 537.243 | $[M-FA18:0-H]^-$ | $C_{24}H_{42}O_{11}P^-$ |
| 437.262 | $[M-15:1ketene-inositol-H]^-$ | $C_{21}H_{42}O_7P^-$ |
| 423.248 | $[M-16:1ketene-inositol-H]^-$ | $C_{20}H_{40}O_7P^-$ |
| 419.254 | $[M-15:1FA-inositol-H]^-$ | $C_{21}H_{40}O_6P^-$ |
| 417.237 | $[M-15:0FA-inositol-H]^-$ | $C_{21}H_{38}O_6P^-$ |
| 409.234 | $[M-17:1ketene-inositol-H]^-$ | $C_{19}H_{38}O_7P^-$ |
| 405.238 | $[M-16:1FA-inositol-H]^-$ | $C_{20}H_{38}O_6P^-$ |
| 403.225 | $[M-16:0FA-inositol-H]^-$ | $C_{20}H_{36}O_6P^-$ |
| 395.253 | $[M-18:1ketene-inositol-H]^-$ | $C_{18}H_{36}O_7P^-$ |
| 391.223 | $[M-17:1FA-inositol-H]^-$ | $C_{19}H_{36}O_6P^-$ |
| 389.208 | $[M-17:0FA-inositol-H]^-$ | $C_{19}H_{34}O_6P^-$ |

| | | |
|---|---|---|
| 377.244 | [M-18:1FA-inositol-H]$^-$ | $C_{18}H_{34}O_6P^-$ |
| 375.230 | [M-18:0FA-inositol-H]$^-$ | $C_{18}H_{32}O_6P^-$ |
| 315.048 | [M-FA-ketene-H]$^-$ | $C_9H_{16}O_{10}P^-$ |
| 297.036 | [M-FA-FA-H]$^-$ | $C_9H_{14}O_9P^-$ |
| 283.261 | [FA18:0-H]$^-$ | $C_{18}H_{35}O_2^-$ |
| 281.246 | [FA18:1-H]$^-$ | $C_{18}H_{33}O_2^-$ |
| 269.246 | [FA17:0-H]$^-$ | $C_{17}H_{33}O_2^-$ |
| 267.230 | [FA17:1-H]$^-$ | $C_{17}H_{31}O_2^-$ |
| 259.090 | [inositolPO$_3$H$_2$-H]$^-$ | $C_6H_{12}O_9P^-$ |
| 255.231 | [FA16:0-H]$^-$ | $C_{16}H_{31}O_2^-$ |
| 253.215 | [FA16:1-H]$^-$ | $C_{16}H_{29}O_2^-$ |
| 241.216 | [FA15:0-H]$^-$ | $C_{15}H_{29}O_2^-$ |
| 241.010 | [inositolPO$_3$H$_2$-H$_2$O-H]$^-$ | $C_6H_{10}O_8P^-$ |
| 239.199 | [FA15:1-H]$^-$ | $C_{15}H_{27}O_2^-$ |
| 223.000 | [inositolPO$_3$H-2H$_2$O-H]$^-$ | $C_6H_8O_7P^-$ |
| 152.996 | [glycerolPO$_3$H$_2$-H]$^-$ | $C_3H_6O_5P^-$ |

Supplementary Table 7. Identified fragment ions of PI 34:1[M-H]$^-$ (3$^{rd}$ carbon isotope ($^{13}C_3$), *m/z* 838.541). Isolated at 1/K$_0$ = 1.427 Vs/cm² and fragmented with -54.5 eV.

| ***m/z* measured** | **Fragment ion** | **Sum formula** |
|---|---|---|
| 838.541 | [M($^{13}C_3$)-H]$^-$ | $C_{42}H_{78}O_{13}P^-$ |
| 598.317–602.331 | [M($^{13}C_{3-n}$)-16:1(0)ketene-H]$^-$ | $^{13}C_{3-n}C_{24+n}H_{52(50)}O_{12}P^-$ |
| 579.291–584.315 | [M($^{13}C_{3-n}$)-16:1(0)FA-H]$^-$ | $^{13}C_{3-n}C_{24+n}H_{50(48)}O_{11}P^-$ |
| 571.292–575.300 | [M($^{13}C_{3-n}$)-18:1(0)ketene-H]$^-$ | $^{13}C_{3-n}C_{22+n}H_{48(46)}O_{12}P^-$ |
| 552.263–557.321 | [M($^{13}C_{3-n}$)-18:1(0)FA-H]$^-$ | $^{13}C_{3-n}C_{22+n}H_{46(44)}O_{11}P^-$ |
| 417.234–422.260 | [M($^{13}C_{3-n}$)-16:1(0)FA-inositol-H]$^-$ | $^{13}C_{3-n}C_{18+n}H_{40(38)}O_6P^-$ |
| 389.207–393.232 | [M($^{13}C_{3-n}$)-18:1(0)FA-inositol-H]$^-$ | $^{13}C_{3-n}C_{16+n}H_{36(34)}O_6P^-$ |
| 315.045–318.056 | [M-FA-ketene-H]$^-$ | $^{13}C_{3-n}C_6H_{16}O_{10}P^-$ |
| 297.031–300.039 | [M-FA-FA-H]$^-$ | $^{13}C_{3-n}C_6H_{14}O_9P^-$ |
| 281.247–286.280 | [($^{13}C_{3-n}$)18:1(0)FA-H]$^-$ | $^{13}C_{3-n}C_{15}H_{34(36)}O_2^-$ |
| 259.022–262.036 | [inositolPO$_3$H$_2$-H]$^-$ | $^{13}C_{3-n}C_3H_{12}O_9P^-$ |
| 253.213–258.242 | [($^{13}C_{3-n}$)16:1(0)FA-H]$^-$ | $^{13}C_{3-n}C_{13}H_{30(32)}O_2^-$ |
| 241.009–244.016 | [inositolPO$_3$H$_2$-H$_2$O-H]$^-$ | $^{13}C_{3-n}C_3H_{10}O_8P^-$ |
| 223.000–226.006 | [inositolPO$_3$H-2H$_2$O-H]$^-$ | $^{13}C_{3-n}C_3H_8O_7P^-$ |
| 152.994–155.000 | [glycerolPO$_3$H$_2$-H]$^-$ | $^{13}C_{3-n}C_3H_6O_5P^-$ |

Supplementary Table 8. Identified fragment ions of PI 36:3[M-H]- (m/z 859.530). Isolated at $1/K_0$ = 1.438 Vs/cm² and fragmented with -55.3 eV. PI 18:0/20:3 was identified as the predominant isomeric structure[9].

| *m/z* measured | Fragment ion | Sum formula |
|---:|---:|---:|
| 859.530 | $[M\text{-}H]^-$ | $C_{42}H_{78}O_{13}P^-$ |
| 621.303 | $[M\text{-ketene16:0-}H]^-$ | $C_{29}H_{50}O_{12}P^-$ |
| 603.290 | $[M\text{-FA16:0-}H]^-$ | $C_{29}H_{48}O_{11}P^-$ |
| 599.319 | $[M\text{-ketene18:3-}H]^-$ | $C_{27}H_{52}O_{12}P^-$ |
| 597.302 | $[M\text{-ketene18:2-}H]^-$ | $C_{27}H_{50}O_{12}P^-$ |
| 595.299 | $[M\text{-ketene18:1-}H]^-$ | $C_{27}H_{48}O_{12}P^-$ |
| 593.265 | $[M\text{-ketene18:0-}H]^-$ | $C_{27}H_{46}O_{12}P^-$ |
| 581.306 | $[M\text{-FA18:3-}H]^-$ | $C_{27}H_{50}O_{11}P^-$ |
| 579.291 | $[M\text{-FA18:2-}H]^-$ | $C_{27}H_{48}O_{11}P^-$ |
| 577.277 | $[M\text{-FA18:1-}H]^-$ | $C_{27}H_{46}O_{11}P^-$ |
| 575.258 | $[M\text{-FA18:0-}H]^-$ | $C_{27}H_{44}O_{11}P^-$ |
| 571.293 | $[M\text{-20:3ketene-}H]^-$ | $C_{25}H_{48}O_{12}P^-$ |
| 553.276 | $[M\text{-FA20:3-}H]^-$ | $C_{26}H_{46}O_{11}P^-$ |
| 441.236 | $[M\text{-16:0FA-inositol-}H]^-$ | $C_{23}H_{38}O_6P^-$ |
| 419.256 | $[M\text{-18:3FA-inositol-}H]^-$ | $C_{21}H_{40}O_6P^-$ |
| 417.240 | $[M\text{-18:2FA-inositol-}H]^-$ | $C_{21}H_{38}O_6P^-$ |
| 415.224 | $[M\text{-18:1FA-inositol-}H]^-$ | $C_{21}H_{36}O_6P^-$ |
| 413.209 | $[M\text{-18:0FA-inositol-}H]^-$ | $C_{23}H_{34}O_6P^-$ |
| 391.230 | $[M\text{-20:3FA-inositol-}H]^-$ | $C_{19}H_{36}O_6P^-$ |
| 315.050 | $[M\text{-FA-ketene-}H]^-$ | $C_9H_{16}O_{10}P^-$ |
| 305.246 | $[\text{FA20:3-}H]^-$ | $C_{20}H_{33}O_2^-$ |
| 297.037 | $[M\text{-FA-FA-}H]^-$ | $C_9H_{14}O_9P^-$ |
| 283.262 | $[\text{FA18:0-}H]^-$ | $C_{18}H_{35}O_2^-$ |
| 281.247 | $[\text{FA18:1-}H]^-$ | $C_{18}H_{33}O_2^-$ |
| 279.231 | $[\text{FA18:2-}H]^-$ | $C_{18}H_{31}O_2^-$ |
| 277.214 | $[\text{FA18:3-}H]^-$ | $C_{18}H_{29}O_2^-$ |
| 259.022 | $[\text{inositol}PO_3H_2\text{-}H]^-$ | $C_6H_{12}O_9P^-$ |
| 255.231 | $[\text{FA16:0-}H]^-$ | $C_{16}H_{31}O_2^-$ |
| 241.010 | $[\text{inositol}PO_3H_2\text{-}H_2O\text{-}H]^-$ | $C_6H_{10}O_8P^-$ |
| 223.000 | $[\text{inositol}PO_3H\text{-}2H_2O\text{-}H]^-$ | $C_6H_8O_7P^-$ |
| 152.994 | $[\text{glycerol}PO_3H_2\text{-}H]^-$ | $C_3H_6O_5P^-$ |

**Supplementary Table 9. Identified fragment ions of PI 34:1[M-H]$^-$ (2$^{nd}$ carbon isotope ($^{13}C_2$), *m/z* 915.593).** Isolated at $1/K_0$ = 1.494 Vs/cm² and fragmented with -59.1 eV. Fragment ions marked with an asterisk result from additionally isolated PI 40:3.

| *m/z* measured | Fragment ion | Sum formula |
|---:|---:|---:|
| 915.593 | [M($^{13}C_3$)-H]$^-$ | $C_{42}H_{78}O_{13}P^-$ |
| 628.282–631.315 | [M($^{13}C_{2-n}$)-18:1(0)FA-H]$^-$<br>[M($^{13}C_{2-n}$)-20:4ketene-H]$^-$ | $^{13}C_{2-n}C_{29+n}H_{50(48)}O_{11}P^-$<br>$^{13}C_{2-n}C_{27+n}H_{56}O_{12}P^-$ |
| 609.343–611.347 | [M($^{13}C_{2-n}$)-20:4FA-H]$^-$ | $^{13}C_{2-n}C_{27+n}H_{54}O_{11}P^-$ |
| 597.300–603.348 | [M($^{13}C_{2-n}$)-22:4(3)ketene-H]$^-$<br>[M($^{13}C_{2-n}$)-20:4FA-H]$^-$ | $^{13}C_{2-n}C_{25+n}H_{52(50)}O_{12}P^-$<br>$^{13}C_{2-n}C_{27+n}H_{46}O_{11}P^-$ |
| 579.290–585.320 | [M($^{13}C_{2-n}$)-22:4(3)FA-H]$^-$ | $^{13}C_{2-n}C_{25+n}H_{50(48)}O_{11}P^-$ |
| 467.247–471.268 | [M($^{13}C_{2-n}$)-18:1(0)FA-inositol-H]$^-$ | $^{13}C_{2-n}C_{23+n}H_{42(40)}O_6P^-$ |
| 417.238–422.257 | [M($^{13}C_{2-n}$)-22:4(3)FA-inositol-H]$^-$ | $^{13}C_{2-n}C_{19+n}H_{40(38)}O_6P^-$ |
| 333.272 | [FA22:3-H]$^-$ | $C_{22}H_{37}O_2^-$ |
| 331.258 | [FA22:4-H]$^-$ | $C_{20}H_{35}O_2^-$ |
| 315.045 | [M-FA-ketene-H]$^-$ | $C_9H_{16}O_{10}P^-$ |
| 311.304 | [FA20:0-H]$^-$ | $C_{20}H_{39}O_2^-$ |
| *305.249 | [FA20:3-H]$^-$ | $C_{20}H_{33}O_2^-$ |
| 303.231 | [FA20:4-H]$^-$ | $C_{20}H_{31}O_2^-$ |
| 297.036 | [M-FA-FA-H]$^-$ | $C_9H_{14}O_9P^-$ |
| 283.263 | [FA18:0-H]$^-$ | $C_{18}H_{35}O_2^-$ |
| 281.246 | [FA18:1-H]$^-$ | $C_{18}H_{33}O_2^-$ |
| 259.020 | [inositolPO$_3$H$_2$-H]$^-$ | $C_6H_{12}O_9P^-$ |
| 255.230 | [FA16:0-H]$^-$ | $C_{16}H_{31}O_2^-$ |
| 253.215 | [FA16:1-H]$^-$ | $C_{16}H_{29}O_2^-$ |
| 241.009 | [inositolPO$_3$H$_2$-H$_2$O-H]$^-$ | $C_6H_{10}O_8P^-$ |
| 222.999 | [inositolPO$_3$H-2H$_2$O-H]$^-$ | $C_6H_8O_7P^-$ |
| 152.995 | [glycerolPO$_3$H$_2$-H]$^-$ | $C_3H_6O_5P^-$ |

**Supplementary Table 10. Overview of GM3-series gangliosides identified in ARSA-/- and ARSA+/+ kidney.** Uncertainties as standard deviation (*n*=4) in parentheses. All ions detected as [M-H]$^-$.

| Name | Chemical sum formula | m/z theo. | 1/K0 [V·s/cm²] | CCS [Å²] | m/z timsTOF | ppm tims TOF | m/z FTICR | ppm FTICR |
|---|---|---:|---:|---:|---:|---:|---:|---:|
| GM3 34:1;O2 | C57H104N2O21 | 1151.705882 | 1.697(1) | 343.9(3) | 1151.708(1) | 2.01 | 1151.7062(04) | 0.31 |
| GM3 34:1;O3 | C57H104N2O22 | 1167.700797 | 1.708(1) | 346.2(3) | 1167.701(1) | 0.20 | 1167.7021(13) | 1.13 |
| GM3 36:1;O2 | C59H108N2O21 | 1179.737182 | 1.730(1) | 350.4(2) | 1179.737(1) | 0.18 | 1179.7390(13) | 1.13 |
| GM3 36:1;O3 | C59H108N2O22 | 1195.732097 | 1.740(3) | 352.5(6) | 1195.733(3) | 0.36 | 1195.7331(08) | 0.66 |
| GM3 38:1;O2 | C61H112N2O21 | 1207.768482 | 1.760(2) | 356.5(3) | 1207.770(1) | 1.69 | 1207.7700(14) | 1.13 |
| GM3 38:1;O3 | C61H112N2O22 | 1223.763397 | 1.769(2) | 358.3(3) | 1223.764(1) | 0.41 | 1223.7620(17) | 1.38 |
| GM3 40:1;O2 | C63H116N2O21 | 1235.799782 | 1.787(2) | 361.9(4) | 1235.800(3) | -0.03 | 1235.8008(14) | 1.13 |
| GM3 40:1;O3 | C63H116N2O22 | 1251.794697 | 1.796(2) | 363.6(4) | 1251.798(1) | 2.66 | 1251.7960(12) | 0.99 |
| GM3 42:1;O2 | C65H120N2O21 | 1263.831082 | 1.810(2) | 366.4(4) | 1263.831(3) | -0.22 | 1263.8321(13) | 1.03 |
| GM3 42:1;O3 | C65H120N2O22 | 1279.825997 | 1.819(1) | 368.1(3) | 1279.829(1) | 1.94 | 1279.8269(13) | 1.02 |

#### Supplementary Table 11. List of all (kidney) sulfatides reported in ARSA-/- mice based on low-resolution MALDI-TOF-MSI (Marsching et al., 2011)[5].

| Lipid class | Name | Chemical sum formula | *m/z* theoretical |
|---|---|---|---|
| SM4 | SM4 32:1;O2 | C38H73NO11S | 750.483160 |
| SM4 | SM4 32:2;O3 | C38H71NO12S | 764.462422 |
| SM4 | SM4 32:1;O3 | C38H73NO12S | 766.478070 |
| SM4 | SM4 34:2;O2 | C40H75NO11S | 776.498807 |
| SM4 | SM4 34:1;O2 | C40H77NO11S | 778.514457 |
| SM4 | SM4 34:2;O3 | C40H75NO12S | 792.493722 |
| SM4 | SM4 34:1;O3 | C40H77NO12S | 794.509372 |
| SM4 | SM4 34:0;O3 | C40H79NO12S | 796.525022 |
| SM4 | SM4 36:2;O2 | C42H79NO11S | 804.530107 |
| SM4 | SM4 36:1;O2 | C42H81NO11S | 806.545757 |
| SM4 | SM4 34:0;O4 | C40H79NO13S | 812.519936 |
| SM4 | SM4 36:2;O3 | C42H79NO12S | 820.525022 |
| SM4 | SM4 36:1;O3 | C42H81NO12S | 822.540672 |
| SM4 | SM4 38:2;O2 | C44H83NO11S | 832.561407 |
| SM4 | SM4 38:1;O2 | C44H85NO11S | 834.577057 |
| SM4 | SM4 37:1;O3 | C43H83NO12S | 836.556322 |
| SM4 | SM4 38:2;O3 | C44H83NO12S | 848.556322 |
| SM4 | SM4 39:1;O2 | C45H87NO11S | 848.592707 |
| SM4 | SM4 38:1;O3 | C44H85NO12S | 850,571972 |
| SM4 | SM4 40:2;O2 | C46H87NO11S | 860.592707 |
| SM4 | SM4 40:1;O2 | C46H89NO11S | 862.608357 |
| SM4 | SM4 39:1;O3 | C45H87NO12S | 864.587622 |
| SM4 | SM4 38:0;O4 | C44H87NO13S | 868.582536 |
| SM4 | SM4 41:2;O2 | C47H89NO11S | 874.608357 |
| SM4 | SM4 40:2;O3 | C46H87NO12S | 876.587622 |
| SM4 | SM4 41:1;O2 | C47H91NO11S | 876.624007 |
| SM4 | SM4 40:1;O3 | C46H89NO12S | 878.603272 |
| SM4 | SM4 42:3;O2 | C48H89NO11S | 886.608357 |
| SM4 | SM4 42:2;O2 | C48H91NO11S | 888.624007 |
| SM4 | SM4 41:2;O3 | C47H89NO12S | 890.603272 |
| SM4 | SM4 42:1;O2 | C48H93NO11S | 890.639657 |
| SM4 | SM4 41:1;O3 | C47H91NO12S | 892.618922 |
| SM4 | SM4 40:0;O4 | C46H91NO13S | 896.613837 |
| SM4 | SM4 42:3;O3 | C48H89NO12S | 902.603272 |
| SM4 | SM4 42:2;O3 | C48H91NO12S | 904.618922 |
| SM4 | SM4 43:1;O2 | C49H95NO11S | 904.655307 |
| SM4 | SM4 42:1;O3 | C48H93NO12S | 906.634572 |
| SM4 | SM4 44:2;O2 | C50H95NO11S | 916.655307 |
| SM4 | SM4 44:1;O2 | C50H97NO11S | 918.670957 |
| SM4 | SM4 43:1;O3 | C49H95NO12S | 920.650222 |
| SM4 | SM4 42:1;O4 | C48H93NO13S | 922.629487 |
| SM4 | SM4 42:0;O4 | C48H95NO13S | 924.645137 |

| | | | |
|---|---|---|---|
| SM4 | SM4 44:1;O3 | C50H97NO12S | 934.665872 |
| SM3 | SM3 34:1;O2 | C46H87N1O16S1 | 940.567281 |
| SM3 | SM3 34:1;O3 | C46H87N1O17S1 | 956.562196 |
| SM3 | SM3 36:1;O2 | C48H91N1O16S1 | 968.598581 |
| SM3 | SM3 38:1;O2 | C50H95N1O16S1 | 996.629881 |
| SM3 | SM3 38:1;O3 | C50H95N1O17S1 | 1012.624796 |
| SM3 | SM3 38:0;O3 | C50H97N1O17S1 | 1014.640446 |
| SM3 | SM3 40:2;O2 | C52H97N1O16S1 | 1022.645531 |
| SM3 | SM3 40:1;O2 | C52H99N1O16S1 | 1024.661181 |
| SM3 | SM3 40:2;O3 | C52H97N1O17S1 | 1038.640446 |
| SM3 | SM3 41:1;O2 | C53H101N1O16S1 | 1038.676831 |
| SM3 | SM3 40:1;O3 | C52H99N1O17S1 | 1040.656096 |
| SM3 | SM3 40:0;O3 | C52H101N1O17S1 | 1042.671746 |
| SM3 | SM3 42:3;O2 | C54H99N1O16S1 | 1048.661181 |
| SM3 | SM3 42:2;O2 | C54H101N1O16S1 | 1050.676831 |
| SM3 | SM3 42:1;O2 | C54H103N1O16S1 | 1052.692481 |
| SM3 | SM3 41:1;O3 | C53H101N1O17S1 | 1054.671746 |
| SM3 | SM3 41:0;O3 | C53H103N1O17S1 | 1056.687396 |
| SM3 | SM3 40:0;O4 | C52H101N1O18S1 | 1058.666661 |
| SM3 | SM3 42:3;O3 | C54H99N1O17S1 | 1064.656096 |
| SM3 | SM3 42:2;O3 | C54H101N1O17S1 | 1066.671746 |
| SM3 | SM3 43:1;O2 | C55H105N1O16S1 | 1066.708131 |
| SM3 | SM3 42:1;O3 | C54H103N1O17S1 | 1068.687396 |
| SM3 | SM3 42:0;O3 | C54H105N1O17S1 | 1070.703046 |
| SM3 | SM3 44:1;O2 | C56H107N1O16S1 | 1080.723781 |
| SM3 | SM3 42:1;O4 | C54H103N1O18S1 | 1084.682311 |
| SM3 | SM3 42:0;O4 | C54H105N1O18S1 | 1086.697961 |
| SB1a | SB1a 34:1;O2 | C60H110N2O29S2 | 1305.699477 |
| SB1a | SB1a 34:1;O3 | C60H110N2O30S2 | 1321.694392 |
| SB1a | SB1a 36:1;O2 | C62H114N2O29S2 | 1333.730777 |
| SB1a | SB1a 38:1;O2 | C64H118N2O29S2 | 1361.762077 |
| SB1a | SB1a 38:0;O2 | C64H120N2O29S2 | 1363.777727 |
| SB1a | SB1a 39:1;O2 | C65H120N2O29S2 | 1375.777727 |
| SB1a | SB1a 38:1;O3 | C64H118N2O29S2 | 1377.756991 |
| SB1a | SB1a 38:0;O3 | C64H120N2O30S2 | 1379.772641 |
| SB1a | SB1a 40:2;O2 | C66H120N2O29S2 | 1387.777727 |
| SB1a | SB1a 40:1;O2 | C66H122N2O29S2 | 1389.793377 |
| SB1a | SB1a 39:1;O3 | C65H120N2O30S2 | 1391.772641 |
| SB1a | SB1a 41:1;O2 | C67H124N2O29S2 | 1403.809027 |
| SB1a | SB1a 40:1;O3 | C66H122N2O30S2 | 1405.788292 |
| SB1a | SB1a 40:0;O3 | C66H124N2O30S2 | 1407.803941 |
| SB1a | SB1a 42:2;O2 | C68H124N2O29S2 | 1415.809027 |
| SB1a | SB1a 42:1;O2 | C68H126N2O29S2 | 1417.824677 |
| SB1a | SB1a 41:1;O3 | C67H124N2O30S2 | 1419.803941 |
| SB1a | SB1a 42:2;O3 | C68H124N2O30S2 | 1431.803942 |
| SB1a | SB1a 42:1;O3 | C68H126N2O30S2 | 1433.819592 |
| SB1a | SB1a 42:0;O3 | C68H128N2O30S2 | 1435.835241 |

#### Supplementary Table 12. Overview of sulfatides identified in ARSA-/- kidney by LC-ESI-TIMS-TOF MS.

Internal standard marked with asterisk. SM4 and SM3 were detected as $[M-H]^{-}$, and SB1a were detected as $[M-2H]^{2-}$. Internal standard marked with asterisk.

| Name | Chemical sum formula | m/z theoret. | CCS [Å²] | $t_R$ | m/z timsTOF | ppm timsTOF |
|---|---|---|---|---|---|---|
| SM4 32:1;O3 | C38H73NO12S | 766.478072 | 280.3 | 8.74 | 766.477 | -1.41 |
| SM4 34:2;O2 | C40H75NO11S | 776.498807 | 284.2 | 9.39 | 776.498 | -0.52 |
| SM4 34:1;O2 | C40H77NO11S | 778.514457 | 285.5 | 10.17 | 778.515 | 0.43 |
| SM4 34:2;O3 | C40H75NO12S | 792.493722 | 285.4 | 9.10 | 792.493 | -1.04 |
| *SM4 35:1;O2 | C41H79NO11S | 792.530107 | 288.8 | 10.70 | 792.530 | 0.10 |
| SM4 34:1;O3 | C40H77NO12S | 794.509372 | 286.8 | 9.89 | 794.509 | 0.01 |
| SM4 34:0;O3 | C40H79NO12S | 796.525022 | 289.4 | 9.66 | 796.525 | -0.17 |
| SM4 36:2;O2 | C42H79NO11S | 804.530107 | 290.4 | 10.52 | 804.530 | -0.20 |
| SM4 36:1;O2 | C42H81NO11S | 806.545757 | 291.7 | 11.18 | 806.544 | -1.74 |
| SM4 36:0;O2 | C42H83NO11S | 808.561407 | 292.3 | 11.49 | 808.562 | 0.39 |
| SM4 36:2;O3 | C42H79NO12S | 820.525022 | 291.6 | 10.24 | 820.525 | 0.19 |
| SM4 36:1;O3 | C42H81NO12S | 822.540672 | 292.3 | 10.91 | 822.541 | 0.16 |
| SM4 38:2;O2 | C44H83NO11S | 832.561407 | 295.9 | 11.45 | 832.562 | 0.11 |
| SM4 38:1;O2 | C44H85NO11S | 834.577057 | 296.5 | 12.07 | 834.575 | -1.93 |
| SM4 37:1;O3 | C43H83NO12S | 836.556322 | 296.4 | 11.42 | 836.555 | -1.87 |
| SM4 36:0;O4 | C42H83NO13S | 840.551237 | 296.2 | 10.57 | 840.552 | 0.65 |
| SM4 38:2;O3 | C44H83NO12S | 848.556322 | 297.4 | 11.20 | 848.556 | -0.38 |
| SM4 39:1;O2 | C45H87NO11S | 848.592707 | 298.9 | 12.54 | 848.590 | -3.11 |
| SM4 38:1;O3 | C44H85NO12S | 850.571972 | 297.9 | 11.84 | 850.569 | 0.97 |
| SM4 40:2;O2 | C46H87NO11S | 860.592707 | 300.7 | 12.40 | 860.588 | -1.87 |
| SM4 40:1;O2 | C46H89NO11S | 862.608357 | 301.9 | 12.97 | 862.606 | -2.73 |
| SM4 39:1;O3 | C45H87NO12S | 864.587622 | 300.4 | 12.33 | 864.587 | -0.92 |
| SM4 38:0;O4 | C44H87NO13S | 868.582536 | 301.8 | 11.51 | 868.582 | -0.15 |
| SM4 41:2;O2 | C47H89NO11S | 874.608357 | 301.6 | 12.56 | 874.609 | 0.32 |
| SM4 40:2;O3 | C46H87NO12S | 876.587622 | 302.2 | 12.16 | 876.587 | -0.57 |
| SM4 41:1;O2 | C47H91NO11S | 876.624007 | 304.3 | 13.38 | 876.623 | -1.49 |
| SM4 40:1;O3 | C46H89NO12S | 878.603272 | 303.3 | 12.78 | 878.604 | 0.60 |
| SM4 42:3;O2 | C48H89NO11S | 886.608357 | 303.7 | 12.52 | 886.604 | -4.91 |
| SM4 42:2;O2 | C48H91NO11S | 888.624007 | 304.4 | 13.25 | 888.623 | -0.79 |
| SM4 41:2;O3 | C47H89NO12S | 890.603272 | 304.6 | 12.59 | 890.603 | -0.53 |
| SM4 42:1;O2 | C48H93NO11S | 890.639657 | 306.6 | 13.77 | 890.638 | -1.31 |
| SM4 41:1;O3 | C47H91NO12S | 892.618922 | 305.2 | 13.17 | 892.618 | -0.54 |
| SM4 40:0;O4 | C46H91NO13S | 896.613837 | 305.8 | 12.44 | 896.612 | -1,71 |
| SM4 42:3;O3 | C48H89NO12S | 902.603272 | 305.4 | 12.28 | 902.601 | -2.18 |
| SM4 42:2;O3 | C48H91NO12S | 904.618922 | 307.1 | 12.86 | 904.618 | -1.32 |
| SM4 42:2;O3 | C48H91NO12S | 904.618922 | 307.9 | 13.09 | 904.619 | 0.09 |
| SM4 43:1;O2 | C49H95NO11S | 904.655307 | 310.4 | 14.12 | 904.655 | -0.74 |
| SM4 42:1;O3 | C48H93NO12S | 906.634572 | 308.4 | 13.57 | 906.634 | -1.07 |
| SM4 44:2;O2 | C50H95NO11S | 916.655307 | 311.2 | 13.82 | 916.654 | -1.32 |
| SM4 44:1;O2 | C50H97NO11S | 918.670957 | 312.8 | 14.45 | 918.670 | -0.65 |
| SM4 43:1;O3 | C49H95NO12S | 920.650222 | 310.1 | 13.95 | 920.649 | -1.64 |
| SM4 42:1;O4 | C48H93NO13S | 922.629487 | 308.6 | 12.52 | 922.628 | -2.04 |
| SM4 42:0;O4 | C48H95NO13S | 924.645137 | 309.7 | 13.30 | 924.644 | -1.34 |
| SM4 44:1;O3 | C50H97NO12S | 934.665872 | 314.0 | 14.29 | 934.668 | 1.97 |
| SM4 46:1;O2 | C52H101NO11S | 946.702258 | 316.9 | 15.09 | 946.704 | 1.41 |
| SM3 34:1;O2 | C46H87N1O16S1 | 940.567281 | 313.0 | 10.11 | 940.566 | -1.17 |

| | | | | | | |
|---|---|---|---|---|---|---|
| SM3 34:1;O3 | C46H87N1O17S1 | 956.562196 | 311.6 | 9.78 | 956.562 | -0.52 |
| SM3 36:1;O2 | C48H91N1O16S1 | 968.598581 | 319.0 | 11.13 | 968.600 | 0.97 |
| SM3 38:1;O2 | C50H95N1O16S1 | 996.629881 | 324.0 | 12.05 | 996.629 | -0.96 |
| SM3 38:1;O3 | C50H95N1O17S1 | 1012.62480 | 321.7 | 11.80 | 1012.622 | -2.96 |
| SM3 40:2;O2 | C52H97N1O16S1 | 1022.64553 | 327.6 | 12.19 | 1022.644 | -1.49 |
| SM3 40:1;O2 | C52H99N1O16S1 | 1024.66118 | 329.8 | 12.91 | 1024.660 | -1.13 |
| SM3 40:0;O2 | C52H101NO16S1 | 1026.67683 | 330.7 | 13.22 | 1026.675 | -1.46 |
| SM3 40:2;O3 | C52H97N1O17S1 | 1038.64045 | 326.3 | 11.89 | 1038.640 | -0.49 |
| SM3 41:1;O2 | C53H101N1O16S1 | 1038.67683 | 332.5 | 13.33 | 1038.677 | -0.20 |
| SM3 40:1;O3 | C52H99N1O17S1 | 1040.65610 | 327.7 | 12.59 | 1040.654 | -2.01 |
| SM3 40:0;O3 | C52H101N1O17S1 | 1042.67175 | 328.0 | 12.50 | 1042.670 | -1.82 |
| SM3 42:3;O2 | C54H99N1O16S1 | 1048.66118 | 332.0 | 12.44 | 1048.660 | -0.89 |
| SM3 42:2;O2 | C54H101N1O16S1 | 1050.67683 | 333.5 | 13.03 | 1050.675 | -1.85 |
| SM3 42:1;O2 | C54H103N1O16S1 | 1052.69248 | 335.6 | 13.71 | 1052.692 | -0.27 |
| SM3 41:1;O3 | C53H101N1O17S1 | 1054.67175 | 330.1 | 13.08 | 1054.673 | 1.05 |
| SM3 42:0;O2 | C54H105NO16S1 | 1054.70813 | 336.8 | 13.95 | 1054.707 | -0.66 |
| SM3 41:0;O3 | C53H103N1O17S1 | 1056.68740 | 330.4 | 12.94 | 1056.689 | 1.04 |
| SM3 40:0;O4 | C52H101N1O18S1 | 1058.66666 | 326.4 | 12.29 | 1058.665 | -1,76 |
| SM3 42:2;O3 | C54H101N1O17S1 | 1066.67175 | 331.4 | 12.75 | 1066.669 | -2.86 |
| SM3 43:1;O2 | C55H105N1O16S1 | 1066.70813 | 337.5 | 14.05 | 1066.708 | 0.20 |
| SM3 42:1;O3 | C54H103N1O17S1 | 1068.68740 | 333.3 | 13.47 | 1068.686 | -1.71 |
| SM3 42:0;O3 | C54H105N1O17S1 | 1070.70305 | 333.3 | 13.34 | 1070.701 | -1.75 |
| SM3 44:1;O2 | C56H107N1O16S1 | 1080.72378 | 340.4 | 14.39 | 1080.722 | -1.21 |
| SM3 42:1;O4 | C54H103N1O18S1 | 1084.68231 | 330.3 | 12.40 | 1084.681 | -0.93 |
| SM3 42:0;O4 | C54H105N1O18S1 | 1086.69796 | 331.5 | 13.14 | 1086.696 | -2.08 |
| SB1a 34:1;O2 | C60H110N2O29S2 | 692.32451 | 395.1 | 8.77 | 692.325 | 0.15 |
| SB1a 34:1;O3 | C60H110N2O30S2 | 700.32187 | 396.1 | 8.50 | 700.321 | -1.19 |
| SB1a 36:1;O2 | C62H114N2O29S2 | 706.34016 | 400.9 | 9.85 | 706.339 | -1.07 |
| SB1a 37:1;O2 | C63H116N2O29S2 | 713.34798 | 402.8 | 10.36 | 713.348 | -0.48 |
| SB1a 36:1;O3 | C62H114N2O30S2 | 714.33762 | 401.5 | 9.60 | 714.337 | -0.71 |
| SB1a 38:1;O2 | C64H118N2O29S2 | 720.35581 | 405.4 | 10.83 | 720.355 | -1.06 |
| SB1a 38:0;O2 | C64H120N2O29S2 | 721.36363 | 405.1 | 11.15 | 721.364 | 0.00 |
| SB1a 38:2;O3 | C64H116N2O30S2 | 727.34544 | 405.4 | 9.94 | 727.345 | -0.19 |
| SB1a 39:1;O2 | C65H120N2O29S2 | 727.36363 | 406.7 | 11.32 | 727.362 | -1.74 |
| SB1a 38:1;O3 | C64H118N2O30S2 | 728.35327 | 406.0 | 10.61 | 728.353 | -0.90 |
| SB1a 38:0;O3 | C64H120N2O30S2 | 729.36109 | 407.3 | 10.42 | 729.360 | -0.86 |
| SB1a 40:2;O2 | C66H120N2O29S2 | 733.36363 | 409.3 | 10.97 | 733.363 | -0.53 |
| SB1a 40:1;O2 | C66H122N2O29S2 | 734.37146 | 410.3 | 11.71 | 734.371 | -0.35 |
| SB1a 39:1;O3 | C65H120N2O30S2 | 735.36109 | 408.0 | 11.08 | 735.362 | 0.58 |
| SB1a 41:1;O2 | C67H124N2O29S2 | 741.37928 | 411.9 | 12.17 | 741.379 | -0.34 |
| SB1a 40:1;O3 | C66H122N2O30S2 | 742.36892 | 410.2 | 11.50 | 742.369 | 0.60 |
| SB1a 40:0;O3 | C66H124N2O30S2 | 743.37674 | 411.8 | 11.35 | 743.377 | -0.04 |
| SB1a 42:2;O2 | C68H124N2O29S2 | 747.37928 | 414.2 | 11.77 | 747.379 | -0.53 |
| SB1a 42:1;O2 | C68H126N2O29S2 | 748.38711 | 414.1 | 12.57 | 748.387 | -0.59 |
| SB1a 41:1;O3 | C67H124N2O30S2 | 749.37674 | 413.1 | 11.94 | 749.375 | -1.85 |
| SB1a 42:3;O3 | C68H122N2O30S2 | 754.36892 | 413.1 | 11.03 | 754.368 | -1.31 |
| SB1a 42:2;O3 | C68H124N2O30S2 | 755.37674 | 415.1 | 11.61 | 755.377 | -0.09 |
| SB1a 42:1;O3 | C68H126N2O30S2 | 756.38457 | 415.1 | 12.38 | 756.385 | -0.09 |
| SB1a 42:0;O3 | C68H128N2O30S2 | 757.39239 | 416.4 | 12.21 | 757.393 | 0.46 |
| SB1a 44:1;O2 | C70H130N2O29S2 | 762.40276 | 419.6 | 13.32 | 762.402 | -1.10 |
| SB1a 42:1;O4 | C68H126N2O31S2 | 764.38202 | 417.0 | 11.32 | 764.383 | 0.80 |
| SB1a 42:0;O4 | C68H128N2O31S2 | 765.38985 | 416.7 | 12.04 | 765.389 | -1.51 |

#### Supplementary Table 13. Overview of sulfatides identified in ARSA-/- kidney by MALDI-MSI.

Uncertainties as standard deviation (*n*=4) in parentheses. Internal standard marked with asterisk. Sulfatides with predominant accumulation in cortex region are marked in *italics*.

| Name | Chemical sum formula | m/z theo. | 1/K0 [V·s/cm²] | CCS [Å²] | m/z timsTOF | ppm tims TOF | m/z FTICR | ppm FTICR |
|---|---|---|---|---|---|---|---|---|
| SM4 18:1;O2 | C24H47NO10S | 540.284791 | 1.115(1) | 228.9(3) | 540.285(1) | 1.13 | 540.28479 | -0.097 |
| SM4 18:0;O3 | C24H49NO11S | 558.295356 | 1.139(1) | 233.7(3) | 558.297(1) | 2.50 | | |
| SM4 32:2;O2 | C38H71NO11S | 748.467507 | 1.350(1) | 275.4(2) | 748.468(1) | 0.26 | | |
| SM4 32:1;O2 | C38H73NO11S | 750.483157 | 1.357(1) | 276.7(2) | 750.482(1) | -1.21 | | |
| SM4 32:2;O3 | C38H71NO12S | 764.462422 | 1.363(1) | 277.8(3) | 764.463(1) | 0.89 | | |
| SM4 32:1;O3 | C38H73NO12S | 766.478072 | 1.368(1) | 278.8(3) | 766.476(1) | -2.70 | | |
| SM4 34:2;O2 | C40H75NO11S | 776.498807 | 1.382(1) | 281.8(2) | 776.498(1) | -0.72 | 776.49881 | 0.061 |
| SM4 33:2;O3 | C39H73NO12S | 778.478072 | 1.371(1) | 279.3(3) | 778.477(1) | -1.44 | | |
| SM4 34:1;O2 | C40H77NO11S | 778.514457 | 1.390(1) | 283.2(3) | 778.515(1) | 0.09 | 778.51446 | 0.116 |
| SM4 34:2;O3 | C40H75NO12S | 792.493722 | 1.391(2) | 283.5(4) | 792.493(1) | -1.54 | | |
| *SM4 35:1;O2 | C41H79NO11S | 792.530107 | 1.406(2) | 286.4(3) | 792.531(1) | 0.91 | | |
| SM4 34:1;O3 | C40H77NO12S | 794.509372 | 1.398(1) | 284.8(2) | 794.509(1) | 0.00 | 794.50937 | 0.006 |
| SM4 34:0;O3 | C40H79NO12S | 796.525022 | 1.414(2) | 288.0(4) | 796.527(1) | 2.61 | 796.52502 | 0.008 |
| SM4 36:2;O2 | C42H79NO11S | 804.530107 | 1.417(1) | 288.6(3) | 804.529(1) | -0.82 | 804.53011 | -0.079 |
| SM4 36:1;O2 | C42H81NO11S | 806.545757 | 1.422(1) | 289.7(2) | 806.547(1) | 1.11 | 806.54576 | 0.124 |
| SM4 34:0;O4 | C40H79NO13S | 812.519936 | 1.418(2) | 288.8(4) | 812.520(1) | 0.23 | 812.51994 | 0.123 |
| SM4 36:2;O3 | C42H79NO12S | 820.525022 | 1.424(2) | 290.0(5) | 820.523(1) | -2.04 | 820.52502 | 0.077 |
| SM4 37:1;O2 | C43H83NO11S | 820.561407 | 1.435(1) | 292.3(3) | 820.561(2) | -0.86 | | |
| SM4 36:1;O3 | C42H81NO12S | 822.540672 | 1.431(1) | 291.3(3) | 822.541(1) | -0.15 | 822.54067 | -0.056 |
| SM4 36:0;O3 | C42H83NO12S | 824.556322 | 1.441(2) | 293.4(3) | 824.557(2) | 1.06 | | |
| SM4 38:2;O2 | C44H83NO11S | 832.561407 | 1.441(1) | 293.4(2) | 832.562(1) | 0.47 | 832.56141 | 0.084 |
| SM4 38:1;O2 | C44H85NO11S | 834.577057 | 1.447(2) | 294.6(3) | 834.577(1) | -0.40 | 834.57706 | 0.051 |
| SM4 37:1;O3 | C43H83NO12S | 836.556322 | 1.441(2) | 293.4(2) | 836.555(1) | -1.10 | 836.55632 | 0.006 |
| SM4 36:0;O4 | C42H83NO13S | 840.551237 | 1.445(1) | 294.1(3) | 840.551(1) | 0.22 | 840.55124 | -0.04 |
| SM4 39:2;O2 | C45H85NO11S | 846.577057 | 1.455(1) | 296.2(2) | 846.578(1) | 0.60 | | |
| SM4 38:2;O3 | C44H83NO12S | 848.556322 | 1.451(1) | 295.3(3) | 848.557(1) | 0.89 | 848.55632 | 0.075 |
| SM4 39:1;O2 | C45H87NO11S | 848.592707 | 1.461(2) | 297.3(3) | 848.591(1) | -1.69 | 848.59271 | 0.066 |
| SM4 38:1;O3 | C44H85NO12S | 850.571972 | 1.457(1) | 296.5(3) | 850.572(1) | -0.55 | 850.57197 | 0.154 |
| SM4 38:0;O3 | C44H87NO12S | 852.587622 | 1.468(1) | 298.7(2) | 852.587(2) | -1.08 | | |
| SM4 40:2;O2 | C46H87NO11S | 860.592707 | 1.467(2) | 298.5(3) | 860.594(1) | 0.92 | 860.59271 | 0.118 |
| SM4 40:1;O2 | C46H89NO11S | 862.608357 | 1.472(1) | 299.5(3) | 862.609(1) | 0.28 | 862.60836 | 0.057 |
| SM4 39:1;O3 | C45H87NO12S | 864.587622 | 1.467(2) | 298.4(3) | 864.588(1) | 0.15 | 864.58762 | 0.041 |
| SM4 39:0;O3 | C45H89NO12S | 866.603272 | 1.480(1) | 301.1(2) | 866.604(1) | 1.24 | | |
| SM4 38:0;O4 | C44H87NO13S | 868.582536 | 1.471(1) | 299.2(3) | 868.582(1) | -0.10 | 868.58254 | 0.132 |
| SM4 41:2;O2 | C47H89NO11S | 874.608357 | 1.483(2) | 301.8(3) | 874.606(1) | -2.35 | 874.60836 | 0.134 |
| SM4 40:2;O3 | C46H87NO12S | 876.587622 | 1.475(1) | 300.1(3) | 876.587(1) | -0.54 | 876.58762 | 0.162 |
| SM4 41:1;O2 | C47H91NO11S | 876.624007 | 1.488(2) | 302.8(4) | 876.623(1) | -0.81 | 876.62401 | 0.063 |
| SM4 40:1;O3 | C46H89NO12S | 878.603272 | 1.482(2) | 301.4(4) | 878.604(1) | 0.57 | 878.60327 | -0.077 |
| SM4 40:0;O3 | C46H91NO12S | 880.618922 | 1.495(1) | 304.0(2) | 880.619(1) | 0.26 | | |
| SM4 39:0;O4 | C45H89NO13S | 882.598187 | 1.486(1) | 302.2(1) | 882.599(1) | 0.35 | | |
| SM4 42:3;O2 | C48H89NO11S | 886.608357 | 1.489(3) | 302.9(5) | 886.608(1) | -0.52 | 886.60836 | 0.013 |
| SM4 42:2;O2 | C48H91NO11S | 888.624007 | 1.494(2) | 303.8(4) | 888.623(1) | -1.44 | 888.62401 | -0.122 |
| SM4 41:2;O3 | C47H89NO12S | 890.603272 | | | *890.603(1)* | *-0.53* | 890.60327 | -0.035 |
| SM4 42:1;O2 | C48H93NO11S | 890.639657 | 1.501(2) | 305.3(4) | 890.638(1) | -1.33 | 890.63966 | -0.103 |
| SM4 41:1;O3 | C47H91NO12S | 892.618922 | 1.494(2) | 303.8(4) | 892.619(1) | -0.02 | 892.61892 | -0.235 |
| SM4 41:0;O3 | C47H93NO12S | 894.634572 | 1.505(2) | 306.0(4) | 894.635(2) | 0.34 | | |
| SM4 40:0;O4 | C46H91NO13S | 896.613837 | 1.496(1) | 304.2(3) | 896.614(1) | 0.60 | 896.61384 | -0.119 |
| SM4 42:3;O3 | C48H89NO12S | 902.603272 | 1.493(2) | 303.5(4) | 902.603(1) | -0.72 | 902.60327 | 0.083 |
| SM4 42:2;O3 | C48H91NO12S | 904.618922 | 1.499(2) | 304.8(3) | 904.619(1) | -0.33 | 904.61892 | 0.149 |
| SM4 43:1;O2 | C49H95NO11S | 904.655307 | 1.516(2) | 308.3(5) | 904.654(1) | -1.17 | 904.65531 | 0.032 |
| SM4 42:1;O3 | C48H93NO12S | 906.634572 | 1.506(2) | 306.3(5) | 906.635(1) | 0.09 | 906.63457 | -0.073 |
| SM4 42:0;O3 | C48H95NO12S | 908.650222 | 1.520(1) | 309.1(1) | 908.651(1) | 1.30 | | |
| SM4 41:0;O4 | C47H93NO13S | 910.629487 | 1.507(2) | 306.5(4) | 910.630(1) | 0.62 | | |
| SM4 44:3;O2 | C50H93NO11S | 914.639657 | 1.514(2) | 307.7(3) | 914.640(1) | 0.18 | | |
| SM4 44:2;O2 | C50H95NO11S | 916.655307 | 1.522(2) | 309.3(5) | 916.656(1) | 0.65 | 916.65531 | -0.033 |
| SM4 44:1;O2 | C50H97NO11S | 918.670957 | 1.529(2) | 310.9(5) | 918.671(1) | 0.05 | 918.67096 | 0.087 |
| SM4 43:1;O3 | C49H95NO12S | 920.650222 | 1.521(3) | 309.3(7) | 920.650(1) | -0.43 | 920.65022 | -0.034 |

| | | | | | | | | |
|---|---|---|---|---|---|---|---|---|
| SM4 42:1;O4 | C48H93NO13S | 922.629487 | 1.514(2) | 307.8(4) | 922.629(1) | -0.26 | 922.62949 | 0.007 |
| SM4 42:0;O4 | C48H95NO13S | 924.645137 | 1.518(2) | 308.6(4) | 924.644(1) | -0.85 | 924.64514 | -0.046 |
| SM4 44:2;O3 | C50H95NO12S | 932.650222 | 1.529(3) | 310.7(5) | 932.651(1) | 1.02 | | |
| SM4 44:1;O3 | C50H97NO12S | 934.665872 | 1.534(2) | 311.8(5) | 934.666(1) | -0.16 | 934.66587 | -0.085 |
| SM4 43:0;O4 | C49H97NO13S | 938.660787 | 1.530(2) | 311.0(5) | 938.660(1) | -1.05 | | |
| SM4 46:1;O2 | C52H101NO11S | 946.702258 | 1.555(1) | 315.9(3) | 946.700(1) | -2.33 | 946.70226 | 0.323 |
| SM4 44:0;O4 | C50H99NO13S | 952.676438 | 1.549(1) | 314.6(1) | 952.674(1) | -2.51 | | |
| SM3 18:1;O2 | C30H57N1O15S1 | 702.337615 | 1.271(1) | 259.6(1) | 702.337(1) | -1.52 | | |
| SM3 18:0;O3 | C30H59N1O16S1 | 720.348180 | 1.285(1) | 262.3(1) | 720.347(1) | -2.19 | | |
| SM3 34:1;O2 | C46H87N1O16S1 | 940.567281 | 1.532(2) | 311.4(4) | 940.567(1) | -0.64 | 940.56728 | 0.100 |
| SM3 34:1;O3 | C46H87N1O17S1 | 956.562196 | 1.528(2) | 310.5(5) | 956.562(1) | -0.02 | 956.56220 | 0.113 |
| SM3 36:1;O2 | C48H91N1O16S1 | 968.598581 | 1.561(2) | 317.0(4) | 968.597(1) | -1.61 | 968.59858 | -0.054 |
| SM3 36:1;O3 | C48H91N1O17S1 | 984.593496 | 1.558(1) | 316.4(2) | 984.592(1) | -1.52 | | |
| SM3 38:2;O2 | C50H93N1O16S1 | 994.614231 | 1.587(2) | 322.1(4) | 994.612(1) | -2.22 | | |
| SM3 38:1;O2 | C50H95N1O16S1 | 996.629881 | 1.590(2) | 322.8(3) | 996.629(1) | -0.48 | 996.62988 | 0.053 |
| SM3 38:2;O3 | C50H93N1O17S1 | 1010.609146 | 1.583(1) | 321.3(1) | 1010.609(1) | -0.34 | | |
| SM3 39:1;O2 | C51H97N1O16S1 | 1010.645531 | 1.604(2) | 325.6(4) | 1010.643(1) | -2.11 | | |
| SM3 38:1;O3 | C50H95N1O17S1 | 1012.624796 | 1.584(2) | 321.5(3) | 1012.623(1) | -2.07 | 1012.62480 | 0.082 |
| SM3 38:0;O3 | C50H97N1O17S1 | 1014.640446 | | | *1014.639(1)* | *-1.13* | 1014.64044 | -0.226 |
| SM3 40:2;O2 | C52H97N1O16S1 | 1022.645531 | 1.613(2) | 327.4(4) | 1022.645(1) | -0.84 | 1022.64553 | 0.121 |
| SM3 40:1;O2 | C52H99N1O16S1 | 1024.661181 | 1.618(2) | 328.3(4) | 1024.662(1) | 1.17 | 1024.66118 | 0.068 |
| SM3 38:0;O4 | C50H97N1O18S1 | 1030.635360 | 1.580(1) | 320.7(2) | 1030.637(1) | 1.83 | | |
| SM3 40:2;O3 | C52H97N1O17S1 | 1038.640446 | 1.605(1) | 325.7(3) | 1038.640(2) | -0.24 | 1038.64044 | -0.207 |
| SM3 41:1;O2 | C53H101N1O16S1 | 1038.676831 | 1.631(2) | 331.0(4) | 1038.676(1) | -0.63 | 1038.67683 | -0.099 |
| SM3 40:1;O3 | C52H99N1O17S1 | 1040.656096 | 1.609(2) | 326.5(3) | 1040.657(1) | 0.82 | 1040.65610 | 0.144 |
| SM3 40:0;O3 | C52H101N1O17S1 | 1042.671746 | | | *1042.673(1)* | *0.82* | 1042.67175 | 0.036 |
| SM3 42:3;O2 | C54H99N1O16S1 | 1048.661181 | 1.631(1) | 330.8(3) | 1048.660(1) | -1.05 | 1048.66118 | -0.196 |
| SM3 42:2;O2 | C54H101N1O16S1 | 1050.676831 | 1.636(2) | 331.9(4) | 1050.676(1) | -1.15 | 1050.67683 | 0.163 |
| SM3 42:1;O2 | C54H103N1O16S1 | 1052.692481 | 1.645(2) | 333.7(4) | 1052.693(1) | 0.30 | 1052.69248 | -0.075 |
| SM3 41:1;O3 | C53H101N1O17S1 | 1054.671746 | 1.624(1) | 329.6(1) | 1054.671(1) | -1.09 | 1054.67175 | 0.101 |
| SM3 41:0;O3 | C53H103N1O17S1 | 1056.687396 | | | *1056.689(1)* | *1.49* | | |
| SM3 40:0;O4 | C52H101N1O18S1 | 1058.666661 | 1.602(2) | 325.0(4) | 1058.666(2) | -0.34 | 1058.66666 | 0.029 |
| SM3 42:3;O3 | C54H99N1O17S1 | 1064.656096 | 1.652(3) | 335.2(6) | 1064.666(3) | 9.23 | | |
| SM3 42:2;O3 | C54H101N1O17S1 | 1066.671746 | 1.627(2) | 330.1(4) | 1066.672(1) | -0.16 | 1066.67175 | 0.103 |
| SM3 43:1;O2 | C55H105N1O16S1 | 1066.708131 | 1.659(3) | 336.5(5) | 1066.709(1) | 0.63 | 1066.70813 | 0.071 |
| SM3 42:1;O3 | C54H103N1O17S1 | 1068.687396 | 1.635(1) | 331.7(3) | 1068.688(1) | 0.89 | 1068.68740 | 0.146 |
| SM3 42:0;O3 | C54H105N1O17S1 | 1070.703046 | | | *1070.702(1)* | *-0.98* | 1070.70305 | 0.073 |
| SM3 41:0;O4 | C53H103N1O18S1 | 1072.682311 | 1.614(1) | 327.5(2) | 1072.682(1) | -0.20 | | |
| SM3 44:2;O2 | C56H105N1O16S1 | 1078.708131 | 1.662(2) | 337.2(5) | 1078.708(1) | -0.05 | | |
| SM3 44:1;O2 | C56H107N1O16S1 | 1080.723781 | 1.671(2) | 339.0(4) | 1080.725(1) | 1.17 | 1080.72378 | 0.111 |
| SM3 42:1;O4 | C54H103N1O18S1 | 1084.682311 | 1.622(1) | 328.9(1) | 1084.681(1) | -1.35 | 1084.68231 | 0.743 |
| SM3 42:0;O4 | C54H105N1O18S1 | 1086.697961 | 1.627(2) | 330.0(3) | 1086.698(1) | -0.19 | 1086.69796 | 0.101 |
| SM3 44:1;O3 | C56H107N1O17S1 | 1096.718696 | 1.663(2) | 337.2(4) | 1096.718(1) | -0.50 | 1096.71870 | 0.009 |
| SM3 46:1;O2 | C58H111N1O16S1 | 1108.755082 | 1.698(3) | 344.4(6) | 1108.757(2) | 1.77 | | |
| SM2a 38:1;O2 | C58H108N2O21S1 | 1199.709253 | 1.750(2) | 354.4(5) | 1199.710(2) | 0.29 | | |
| SM2a 38:1;O3 | C58H108N2O22S1 | 1215.704168 | 1.747(2) | 353.9(4) | 1215.706(2) | 1.75 | | |
| SM2a 40:1;O2 | C60H112N2O21S1 | 1227.740553 | 1.776(2) | 359.7(4) | 1227.742(1) | 1.20 | | |
| SM2a 40:1;O3 | C60H112N2O22S1 | 1243.735468 | 1.770(2) | 358.4(4) | 1243.737(2) | 0.99 | | |
| SM2a 42:1;O2 | C62H116N2O21S1 | 1255.771853 | 1.801(2) | 364.7(4) | 1255.773(1) | 0.61 | | |
| SM2a 42:1;O3 | C62H116N2O22S1 | 1271.766768 | 1.792(1) | 362.7(1) | 1271.767(1) | 0.52 | | |
| SB1a 34:1;O2 | C60H110N2O29S2 | 1305.699477 | 1.766(2) | 357.3(3) | 1305.698(1) | -1.17 | 1305.69948 | 0.167 |
| SB1a 34:1;O3 | C60H110N2O30S2 | 1321.694392 | 1.778(1) | 359.8(1) | 1321.695(1) | 0.31 | 1321.69439 | -0.131 |
| SB1a 36:1;O2 | C62H114N2O29S2 | 1333.730777 | 1.792(1) | 362.7(1) | 1333.733(1) | 1.40 | 1333.73078 | 0.379 |
| SB1a 36:1;O3 | C62H114N2O30S2 | 1349.725692 | 1.800(1) | 364.1(1) | 1349.727(1) | 0.71 | 1349.72569 | 0.341 |
| SB1a 38:1;O2 | C64H118N2O29S2 | 1361.762077 | 1.814(1) | 366.9(2) | 1361.763(1) | 0.40 | 1361.76208 | 0.019 |
| SB1a 39:1;O2 | C65H120N2O29S2 | 1375.777727 | 1.824(1) | 368.9(2) | 1375.778(1) | 0.49 | 1375.77773 | 0.419 |
| SB1a 38:1;O3 | C64H118N2O29S2 | 1377.756991 | 1.819(5) | 367(1) | 1377.757(2) | -0.30 | | |
| SB1a 40:2;O2 | C66H120N2O29S2 | 1387.777727 | 1.831(2) | 370.2(3) | 1387.778(1) | 0.52 | 1387.77773 | 0.505 |
| SB1a 40:1;O2 | C66H122N2O29S2 | 1389.793377 | 1.836(3) | 371.3(6) | 1389.796(1) | 1.56 | 1389.79338 | 0.170 |
| SB1a 38:0;O4 | C64H120N2O31S2 | 1395.767556 | 1.809(1) | 365.9(2) | 1395.768(2) | 0.57 | | |
| SB1a 41:1;O2 | C67H124N2O29S2 | 1403.809027 | 1.847(3) | 373.2(3) | 1403.807(1) | -1.35 | 1403.80903 | 0.103 |
| SB1a 40:1;O3 | C66H122N2O30S2 | 1405.788292 | 1.845(3) | 373.2(7) | 1405.787(1) | -0.76 | 1405.78829 | -0.023 |
| SB1a 42:2;O2 | C68H124N2O29S2 | 1415.809027 | 1.855(3) | 375.2(6) | 1415.809(1) | 0.03 | 1415.80903 | -0.053 |
| SB1a 42:1;O2 | C68H126N2O29S2 | 1417.824677 | 1.859(8) | 376(2) | 1417.825(1) | -0.05 | 1417.82468 | 0.077 |
| SB1a 40:0;O4 | C66H124N2O31S2 | 1423.798857 | 1.830(1) | 370.0(1) | 1423.800(3) | 0.49 | | |
| SB1a 42:2;O3 | C68H124N2O30S2 | 1431.803942 | 1.858(1) | 375.6(2) | 1431.802(1) | -1.60 | 1431.80394 | -0.120 |

| | | | | | | | | |
|---|---|---|---|---|---|---|---|---|
| SB1a 42:1;O3 | C68H126N2O30S2 | 1433.819592 | 1.854(6) | 375(1) | 1433.820(1) | 0.62 | 1433.81959 | 0.087 |
| SB1a 41:0;O4 | C67H126N2O31S2 | 1437.814507 | 1.839(2) | 371.8(3) | 1437.817(3) | 1.84 | | |
| SB1a 44:1;O2 | C70H130N2O29S2 | 1445.855977 | 1.872(2) | 378.6(4) | 1445.857(1) | 0.60 | | |
| SB1a 42:0;O4 | C68H128N2O31S2 | 1451.830157 | 1.85(10) | 374(2) | 1451.834(1) | 2.58 | | |

#### Supplementary Table 14. Prm-PASEF fragmentation patterns of sulfatide isoforms identified by MALDI-timsTOF-MSI. Color code represents the degree of hydroxylation.

| | Name | [m/z] | 1/K0 | sample | CE [ -eV] | 96.96 m/z $[HSO_4]^-$ | 241.00 m/z $[Gal-SO_3]^-$ | 315.04 m/z $[RCF-Gal-SO_3]^-$ | 403.06 m/z $[Glu-Gal-SO_3]^-$ | 444.08 m/z $[GalNAc-Gal-SO_3]^-$ | 558.29 m/z $[O3-SPB-Gal-SO_3]^-$ | 568.28 m/z [α-OH-FA loss] |
|---|---|---|---|---|---|---|---|---|---|---|---|---|
| | SPB 18:1;O2 | 540,2850 | 1,116 | ISOM_1 | 65,0 | yes | yes | no | no | no | - | - |
| SM4 | SM4 32:2;O3 | 764,4610 | 1,362 | ISOM_1 | 81,0 | yes | yes | no | no | no | yes | yes |
| SM4 | SM4 32:1;O3 | 766,4770 | 1,368 | ISOM_2 | 81,7 | yes | yes | no | no | no | yes | yes |
| SM4 | SM4 34:1;O2 | 778,5150 | 1,393 | IMP_1 | 84,2 | yes | yes | no | no | no | no | no |
| SM4 | SM4 34:1;O3 | 796,5130 | 1,391 | IMP_2 | 85,0 | yes | yes | no | no | no | no | yes |
| SM4 | SM4 36:1;O2 | 806,5400 | 1,424 | IMP_1 | 89,4 | yes | yes | no | no | no | no | no |
| SM4 | SM4 34:0;O4 | 812,5200 | 1,410 | IMP_2 | 89,1 | yes | yes | no | no | no | yes | yes |
| SM4 | SM4 36:1;O3 | 822,5400 | 1,433 | ISOM_1 | 91,4 | yes | yes | no | no | no | no | yes |
| SM4 | SM4 38:2;O2 | 832,5600 | 1,443 | ISOM_2 | 93,5 | yes | yes | no | no | no | no | no |
| SM4 | SM4 38:1;O2 | 834,5750 | 1,448 | IMP_1 | 94,4 | yes | yes | no | no | no | no | no |
| SM4 | SM4 36:0;O4 | 840,5520 | 1,436 | IMP_2 | 94,5 | yes | yes | no | no | no | yes | yes |
| SM4 | SM4 38:2;O3 | 848,5573 | 1,458 | ISOM_1 | 95,2 | yes | yes | no | no | no | no | yes |
| SM4 | SM4 39:1;O2 | 848,5915 | 1,463 | ISOM_2 | 95,3 | yes | yes | no | no | no | no | no |
| SM4 | SM4 38:1;O3 | 850,5700 | 1,459 | ISOM_1 | 95,3 | yes | yes | no | no | no | no | yes |
| SM4 | SM4 40:2;O2 | 860,5920 | 1,470 | ISOM_2 | 95,6 | yes | yes | no | no | no | no | no |
| SM4 | SM4 40:1;O2 | 862,6070 | 1,473 | IMP_1 | 95,7 | yes | yes | no | no | no | no | no |
| SM4 | SM4 38:0;O4 | 868,5830 | 1,463 | IMP_2 | 95,8 | yes | yes | no | no | no | yes | yes |
| SM4 | SM4 41:1;O2 | 876,6240 | 1,476 | IMP_2 | 96,3 | yes | yes | no | no | no | no | no |
| SM4 | SM4 40:1;O3 | 878,6034 | 1,482 | ISOM_1 | 96,0 | yes | yes | no | no | no | no | yes |
| SM4 | SM4 42:2;O2 | 888,6300 | 1,494 | ISOM_2 | 96,4 | yes | yes | no | no | no | no | no |
| SM4 | SM4 42:1;O2 | 890,6400 | 1,504 | IMP_1 | 96,7 | yes | yes | no | no | no | no | no |
| SM4 | SM4 41:1;O3 | 892,6190 | 1,496 | ISOM_1 | 96,5 | yes | yes | no | no | no | no | yes |
| SM4 | SM4 40:0;O4 | 896,6140 | 1,487 | IMP_2 | 96,6 | yes | yes | no | no | no | yes | yes |
| SM4 | SM4 42:2;O3 | 904,6180 | 1,500 | ISOM_2 | 96,6 | yes | yes | no | no | no | no | yes |
| SM4 | SM4 42:1;O3 | 906,6350 | 1,510 | ISOM_1 | 97,0 | yes | yes | no | no | no | no | yes |
| SM4 | SM4 44:1;O2 | 918,6700 | 1,533 | IMP_1 | 97,7 | yes | yes | no | no | no | no | no |
| SM4 | SM4 43:1;O3 | 920,6500 | 1,523 | ISOM_2 | 97,4 | yes | yes | no | no | no | no | yes |
| SM4 | SM4 42:0;O4 | 924,6450 | 1,510 | IMP_2 | 97,4 | yes | yes | no | no | no | yes | yes |
| SM4 | SM4 44:1;O3 | 934,6660 | 1,536 | ISOM_1 | 97,9 | yes | yes | no | no | no | no | yes |
| SM3 | SM3 34:1;O3 | 956,5600 | 1,536 | ISOM_2 | 97,8 | yes | yes | yes | yes | no | no | *yes* |

| | | | | | | | | | | | | |
|---|---|---|---|---|---|---|---|---|---|---|---|---|
| | SM3 38:1;O3 | 1012,6240 | 1,585 | ISOM_2 | 99,5 | yes | yes | yes | yes | no | no | yes |
| | SM3 40:1;O2 | 1024,6600 | 1,619 | IMP_1 | 100,0 | yes | yes | yes | yes | no | no | no |
| | SM3 40:1;O3 | 1040,6560 | 1,613 | ISOM_2 | 100,0 | yes | yes | yes | yes | no | no | yes |
| | SM3 42:2;O3 | 1050,6760 | 1,638 | ISOM_1 | 100,0 | yes | yes | yes | yes | no | no | yes |
| | SM3 42:1;O2 | 1052,6900 | 1,647 | IMP_1 | 100,0 | yes | yes | yes | yes | no | no | no |
| | SM3 40:0;O4 | 1058,6700 | 1,590 | IMP_2 | 100,0 | yes | yes | yes | yes | no | yes | *yes* |
| | SM3 42:1;O3 | 1068,6880 | 1,638 | ISOM_2 | 100,0 | yes | yes | yes | yes | no | no | *yes* |
| | SM3 44:1;O2 | 1080,7250 | 1,674 | IMP_1 | 100,0 | yes | yes | yes | yes | no | no | no |
| | SM3 42:0;O4 | 1086,7000 | 1,616 | IMP_2 | 100,0 | yes | yes | yes | yes | no | *yes* | *yes* |
| SM2a | SM2a 40:1;O2 | 1227,7355 | 1,775 | ISOM | 100,0 | yes | yes | no | no | yes | no | no |
| SB1a | SB1a 34:1;O2 | 1305,6969 | 1,754 | ISOM | 100,0 | yes | yes | no | no | yes | no | no |
| | SB1a 34:1;O3 | 1321,6886 | 1,778 | ISOM | 100,0 | yes | yes | no | no | yes | no | no |
| | SB1a 38:1;O3 | 1377,7460 | 1,807 | ISOM | 100,0 | yes | yes | no | no | yes | no | no |
| | SB1a 40:1;O2 | 1389,7843 | 1,822 | ISOM | 100,0 | yes | yes | no | no | yes | no | no |
| | SB1a 42:2;O2 | 1415,8032 | 1,834 | ISOM | 100,0 | yes | yes | no | no | yes | no | no |
| | SB1a 42:1;O3 | 1433,8122 | 1,836 | ISOM | 100,0 | yes | yes | no | no | yes | no | no |

#### Supplementary Table 15. Relative values for structure-CCS-relationships of sulfatide sub-classes.

Relative differences in CCS values were obtained by a parallel line model using a 2nd order polynomial fit. The average CCS-reducing effect of single site FA unsaturation amounts to $\overline{IV}$ = 1.3 ± 0.2 Å$^2$. 1: The relative difference was obtained based on MALDI MSI and LC-TIMS-MS data. Therefore, the latter was shifted by the mean relative deviation.

| | a: *X*(1:O2) | b: *X*(1:O3) |
|---|---|---|
| ***I: SM4 – SM3*** | 28.2 ± 0.1 | 25.4 ± 0.2 |
| ***II: SM3 – SM2a*** | 31.3 ± 0.2 | 31.5 ± 0.5 |
| ***III: SM3 – GM3*** | 33.2 ± 0.2 | 36.3 ± 0.2 |
| | **c: *X*(1:O2) - *X*(2:O2)** | **d: *X*(1:O3) - *X*(2:O3)** |
| ***IV₁: SM4*** | *1.3 ± 0.2* | *1.3 ± 0.2* |
| ***IV₂: SM3*** | 1.3 ± 0.3 | 1.3 ± 0.2 |
| | **e: *X*(0:O3)[1]** | **f: *X*(0:O4)** |
| ***V: SM4 – SM3*** | 22.2 ± 0.2 | 20.9 ± 0.4 |
